## Supplementary File for "Best Paper awards lack transparency, inclusivity, and support for Open Science"

### SI_methods_results

ML

27/10/2023


- Supplementary Methods
  - Figure S1.
  - Project aim and general approach.
  - Award inclusion criteria.
  - Search and screening.
  - Screening cross-checking.
  - Data extraction.
  - Data extraction cross-checking.
  - Deviations from the protocol.
  - Data processing and analyses - an overview.
  - Project set up.
  - Loading data.
  - Meta-data tables.
    - Table S1.
    - Table S2.
    - Table S3.
    - Table S4.
- Supplementary Results
  - Characteristics of SCImago Subject Area (SA) journal lists and screening data.
    - Figure S2.
  - Overlap of journals across SCImago Subject Areas.
  - Award-level dataset characteristics.
    - Figure 1A - Screening effort: count numbers of journals screened to get 10 shortlisted awards:
    - Figure 1B - Counts of incuded awards with and without award description:
  - Award descriptions.
    - Figure 1C - Plot of award description lengths - overall:
    - Figure S3.
  - Award ages
    - Figure 1D - award age verall distribution:
    - Figure S4.
    - Figure 4 - Combine 4 panels and save:
    - Figure 2A - Awarding bodies:
    - Figure 2B - target awardee career stages:
    - Figure 2 - combine 2 panels and save:
    - Figure 3 - summary of extracted award characteristics:
    - Figure 3A - Summary plots of key award characteristics:
    - Figure 3B - Summary plots of key award characteristics:
    - Figure 3 - combine 2 panels and save:
  - Award characteristics by Subject Area.
    - Figure S5.
    - Figure S6.
    - Figure S7.
    - Figure S8.
    - Figure S9.
    - Figure S10.
    - Figure S11.
    - Figure S12.
    - Figure S13.
    - Figure S14.
    - Figure S15.
    - Figure S16.
    - Figure S17.
  - Text mining of award descriptions.
    - Figure 4A - frequencies of specific words - all mentions in award descriptions.
    - Figure 4B - frequencies of specific words - first mentions in award descriptions.
    - Figure 4 - combine 2 panels and save.
  - Characteristics of past winners.
    - Figure S18.
    - Figure S19.
    - Figure S20.
    - Figure S21.
  - Counts of extracted past awardees by decade:
  - Awardee gender.
    - Figure S22
    - Figure S23.
    - Figure S24.
    - Figure 5A - plot of gender counts by decade:
    - Figure 5B - plot of gender proportions by decade:
    - Figure S25.
    - Figure S26.
    - Figure S27.
  - Affiliation country.
    - Figure S28.
    - Figure S29.
    - Figure S30.
    - Figure 5C - plot of counts of top 10 country affiliations by decade:
    - Figure 5D - proportions of top 10 affiliation countries by decade:
    - Figure S31.
    - Figure 5 - combine 4 panels and save:
  - Country productivity context.
    - Figure S32.
    - Figure S33.
  - Analyses of past awardees first names.
    - Figure S34.
  - Affiliation institutions of past awardees.
    - Figure S35.
  - Cash and perks for the winners.
    - Figure 6A - mentions of cash awards:
    - Figure S36.
    - Figure 6B - disclosed values of cash awards:
    - Figure 6 - Combine 2 panels and save:
    - Figure S37.
    - Figure S37.
  - Text mining of the description of the perks and benefits.
    - Figure S38.
    - Figure S39.
    - Figure 7A - Individual winners data - presence of a personal profile:
    - Figure 7B - Individual winners data - presence of a personal photo:
    - Figure 7 - combine 2 panels and save:
  - Statistical analyses and visualisations of awardees data - using award years.
    - Figure S40.
    - Figure S41.
    - Figure S42.
  - Analyses of awards - using award age.
    - Figure S43.
    - Figure S44.
    - Figure S45.
    - Figure S46.
    - Figure S47.
    - Figure S48.
    - Figure S49.
    - Figure S50.
    - Figure S51.
    - Figure S52.
    - Figure S53.
  - Author contributions
    - Figure S54.
  - Session Info


### Supplementary Methods

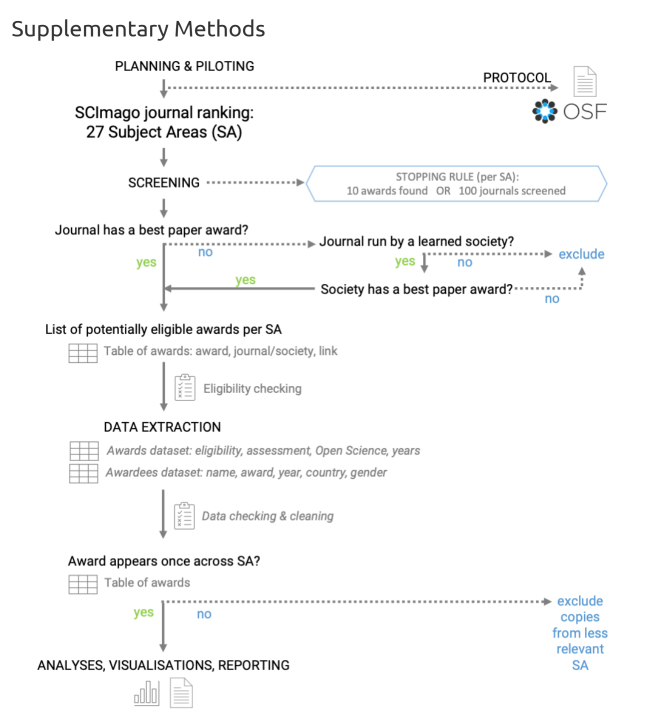


{#id .class width = 50% height = 50%}.

Figure S1.

*Project workflow.*

#### Project aim and general approach.

"Best paper" (or equivalent) awards are usually associated with specific journals / societies / publishers. We aimed to conduct a systematic-like search to collect a sample of highly regarded awards (from top-ranking journals and societies) across a broad range of disciplines. We conducted the process collaboratively and transparently using shared Google Sheets documents (file copies were provided and re-integrated for participants who were not able to access online Google Docs).

The initial project description and contribution guidelines were publicly available as GitHub pages: https://mlagisz.github.io/survey_best_paper_awards/. Project contributors were recruited via Open Science communities and conferences. They helped to pilot the project procedures and provided feedback on the project protocol, which was then updated to it second version. roject contributors self-assigned to the individual project tasks. Their actual contributions (completed tasks) were recorded on Google Sheets. Detailed project procedures were shared as a dedicated Project Manual on Google Docs. This document, with incorporated users' feedback, is available at https://docs.google.com/document/d/1aw_HhKawpi264njGi--Ms-x-bOs_uQJBiOepQKE89pQ/edit?usp=sharing.

#### Award inclusion criteria.

We followed pre-specified criteria outlined in the registered protocol (https://osf.io/7ez28) to increase data consistency and enable comparisons across disciplines:
1. We only included awards for a single published research contribution in a format of a research article (theses / dissertations are allowed if published as a journal article).
2. W only included awards managed by a journal, publisher, or a learned society.
3. We excluded awards that are discontinued, and awards that are restricted to applicants from underrepresented groups, e.g., women-only / minorities-only awards.

#### Search and screening.

Our starting point was from searching academic journals and societies/associations to identify those that award "Best Paper" prizes. To do so, we first used journal lists from 27 SCImago Subject Areas rankings (SCImago Journal & Country Rank (SJR) website at https://www.scimagojr.com/journalrank.php). To represent SAs evenly, we considered the top 100 journals from each SA list, or we will stop if we find 10 relevant awards before reaching this threshold.

During the intial screening, for each journal list representing a SSCImago Subject Area, we checked if journal websites or announcements include "best paper" awards , and if not, we checked if journals are run by learned societies (i.e. are offciel journals of a given learned society/ies). If so, we also checked if a relevant learned society awards a "best paper" prize that fits in a given subject area being assessed. If a society awards multiple relevant awards in journals in the same subject area, we choose the one from our top 100 SA list or an award that appears to be most prestigious (e.g., with highest monetary value, oldest).

For each subject area, we collated a preliminary list of 10-15 awards that meet our selection criteria for data extraction. If some of these awards got excluded during data extraction, we went back to the screening stage for that subject area to find replacement awards until we had 10 awards that are eligible for inclusion, or we ran out of journals to screen within the top 100 Subject Area list positions.

#### Screening cross-checking.

All screened journal lists from 27 SCImago Subject Areas rankings (https://www.scimagojr.com) were independently cross-checked to detect potential mistakes (it is easy to miss a note about awards on journals or organizational webpages, misinterpret eligibility of the award, etc.).

Project contributors self-assigned to one (or more) SCImago Subject Area - always different from the one they did initial Subject Areas screening.

The checking person confirmed if screening information and decision recorded by the initial screener is correct and accurate.

The checking person recorded in a new column (Award_name_extract) the name of the eligible award to be extracted and recorded any screening comments in additional columns.

Obvious mistakes were fixed and any unclear or disputable cases have been commented on and discussed before being resolved.

#### Data extraction.

For each 27 SCImago Subject Areas journal ranking (https://www.scimagojr.com), we extracted data on max. 10 awards that meet our selection criteria.

For each award, its eligibility (as outlined in "Selection criteria") for inclusion was confirmed during data extraction, and if deemed not eligible, no further data was be extracted.

Using a pre-piloted Google Form, we extracted relevant data on each award from the websites (e.g., journals, societies, publishers) or other publicly available documents (e.g., instructions for applicants).

The extracted pre-specified data included information on the granting journal / publisher / society, award eligibility criteria, award assessment criteria, and source of information on the past awardees (full metadata is provided alongside the data files).

We archived web pages with award descriptions by saving them to .pdf files named after award names (as used in data extraction documents).

The second stap of data extraction focused on the identities and characteristics of the past participants. For collecting data on individual winners, we had a data collection directly on a Google Sheet. We only collect data on winners from years 2001-2022. This data was used to infer patterns related to individual winners' gender and country of affiliation. Specifically: Gender (binary: F = female, M = male, ? - unassigned) was be assigned to the past awardee names using information (pronouns, images, names) available on award websites, personal or institutional websites, or using https://gender-api.com/, with an accuracy threshold of >95 for assigning gender from first names. We noted instances of non-binary gender (not applicable) and also when we are unable to assign binary gender. Affiliation information (institution, country) was be assigned to past winners' names for the year when the award was received using information available on award websites, personal or institutional websites, or information in the awarded paper. Additional information was also recorded: links to the websites with relevant information, type of source of information (award page, award announcement, article, other), whether award is shared (more than one author from one winning paper), whether photo or bio are provided on award page or award announcement, reference information and DOI for winning article, any additional notes.

#### Data extraction cross-checking.

Extracted main award characteristics data (awarding body, description, etc.) has been independently cross-checked by a second researcher (i.e. one that did not extract the data). Obvious mistakes were fixed and any unclear or disputable cases have been commented on and discussed before being resolved.

Extracted additional data (gender, affiliation country, and additional details on the award and winners) has been independently cross-checked by a second researcher (i.e. one that did not extract the data). Obvious mistakes were fixed and any unclear or disputable cases have been commented on and discussed before being resolved.

#### Deviations from the protocol.

The pre-registered protocol is available on OSF (https://osf.io/8mxpq/).
We acknowledge the following deviations from the protocol:

1. We coded whether the award goes to an individual author (explicitly named) or to an article (without mentioning individual author/s). Some awards only lists winning articles and other focus on specific authors who were the prizewinners (sometimes even not mentioning for which exact article they got an award).
2. In the protocol, we planned only to extract awardee names, gender and affiliations for winners from 2001 onwards. However, such data is only meaningful when an award is given to a specific individual researcher - thus, we did not extract such information for awards that only list winning papers (i.e. were awarded to the whole authorship team, not a particular person within the team).
3. We coded whether the individual winners got their photo and/or bio shared online by the journals/societies. Photographs and bios make assigning gender easier and provide some extra recognition and visibility for the winners.
4. We recorded the information (reference and DOI) on the winning article identity, where provided or findable. However, we did not use this information in the analyses, as it was solely for documentation and cross-checking purposes, since we extracted information on winner's affiliation from the winning article when affiliation it was not reported on the award webpage or announcement.
5. We recorded whether award criteria mention article impact indices (e.g. number of downloads or citations) and whether impact indices were the only criteria. Although impact indices give an impression of objectivity they are known to be biased and not necessarily reflect the quality of the published research. As such, we decided to include this information in our assessment of the award characteristics.
6. For the analyses, we did not remove Subject Areas that had data extracted from fewer than 5 awards, as initially planned. This is because our main results are presented as totals across all Subject Areas. Also, the two Subject Areas to be potentially excluded were Medicine and Immunology, and we felt that removing them would make our findings less complete.
7. For text mining of award descriptions, we ad hoc grouped the most common terms (truncated to word stems) into terms representing generic excellence and terms explicitly related to transparency and robustness.
8. We were not able to seek clarifications from the award committees or contact person, because most of the award descriptions lacked contact details (see our additional results on the availability of the contact details).
9. In our plots by decade, we also showed the incomplete decade 2021-2022 - in our protocol we stated that only decades 2001-2010 and 2011-2020 will be shown.
10. We used the year of earliest listed awardee as a proxy of award age and performed exploratory analyses of its associations with award characteristics.

#### Data processing and analyses - an overview.

The main steps were as follows:

1. Uploading cross-checked meta-data, screening data and extracted data into R.
2. Creating meta-data tables (variable descriptions) for each data set.
3. Data pre-processing (e.g. merging SA into a single data frame, removing excluded awards).
4. Summarizing data sets via tabulation and filtering.
5. Creating figures for the supplementary materials - shown as in-text figures.
6. Creating figures for the main manuscript text - saved as stand-alone files.

NOTE: Steps 3-6 were done for each data set in turn.

All code and results are provided below. Code for figures used in the main text is shown, but not evaluated for this supplementary document, because these figures are presented in the main text.

#### Project set up.

NOTE: project github page can be found at https://github.com/mlagisz/survey_best_paper_awards.

Loading packages and general settings.

```
#set global code chunk parameters for knitting:
knitr::opts_chunk$set(echo = TRUE)

#load libraries (install if absent):
pacman::p_load(tidyverse, #various tools
               here, #easy file paths
                DT, #data tables
                knitr, #document knitting
                ggimage, #adding images
                ggcharts, #plots functionality
                scales, #plots extra
                readxl, #loading Excel
                forcats, #category wrangling
                ggupset, #upset plots
                patchwork, #plot assembly
                stringr, #text wrangling
                ggbeeswarm, #beeswarm plots
                tidytext, #text processing
                stopwords, #text processing
                tokenizers, #text processing
                ggpatterns, #patterning for plots
                wacolors, #color-blind friendly color palettes
                lme4, #linear models
                sjPlot) #for tab_model reporting
```

```
## Warning: package 'ggpatterns' is not available for this version of R
##
## A version of this package for your version of R might be available elsewhere,
## see the ideas at
## https://cran.r-project.org/doc/manuals/r-patched/R-admin.html#Installing-packages
```

```
## Warning: 'BiocManager' not available.  Could not check Bioconductor.
##
## Please use `install.packages('BiocManager')` and then retry.
```

```
## Warning in p_install(package, character.only = TRUE, ...):
```

```
## Warning in library(package, lib.loc = lib.loc, character.only = TRUE,
## logical.return = TRUE, : there is no package called 'ggpatterns'
```

```
## Warning in pacman::p_load(tidyverse, here, DT, knitr, ggimage, ggcharts, : Failed to install/load:
## ggpatterns
```

```
## may need:
# install.packages("remotes")
# remotes::install_github("coolbutuseless/ggpattern")
library(ggpattern)

## set a global plotting theme with background transparency:
theme_set(theme_classic(base_size = 14) +
            theme(panel.background = element_blank(),
                  plot.background = element_blank(),
                  legend.background = element_blank(),
                  text = element_text(colour = "white")))
```

#### Loading data.

Load SCImago Subject Area -level data set and an associated meta-data table:

```
# accessing all the sheets
#sheet_names <- excel_sheets(here("data", "scimagojr 2021  Subject Areas.xlsx"))
sheet_names <- excel_sheets(here("data", "scimagojr 2021  Subject Areas_screening.xlsx"))
sheet_names <- sheet_names[-1] #remove first sheet with meta-data
SA_list_all <- lapply(sheet_names, function(x)
  {as.data.frame(read_excel(here("data", "scimagojr 2021  Subject Areas.xlsx"), sheet = x))}) #read all sheets
names(SA_list_all) <- sheet_names  #rename list elements
#lapply(SA_list_all, names) #extract column names
#unique(do.call(rbind, lapply(SA_list_all, names))) #making sure all have same column names
SAdata <- do.call(rbind, SA_list_all)
names(SAdata) <- gsub(" ","_", names(SAdata)) #change spaces to _ in the column names
names(SAdata) <- gsub("\\.","", names(SAdata)) #remove . the column names
SAdata <- SAdata %>% drop_na(Subject_area)
#table(is.na(SAdata$Title)) #note no empty journal names

SAmeta <- read_excel(here("data", "scimagojr 2021  Subject Areas_screening.xlsx"),
                     sheet = 1, skip = 1) #load SCImago SA meta-data
```

Load award-level data set and an associated meta-data table:

```
#BP load and prepare award-level meta-data
BPmeta <- read_excel(here("data", "Survey-Best_paper_awards (Responses)_SHAREDCOPY_checked.xlsx"),
                     sheet = 2, skip = 1) #load and skip first line

## load and prepare main extracted award-level data
BPdata <- read_excel(here("data","Survey-Best_paper_awards (Responses)_SHAREDCOPY_checked.xlsx"),
                     sheet = 1) #load main award data

names(BPdata) <- gsub(" ","_", names(BPdata)) #change spaces to _ in the column names

#rename selected data columns
BPdata <- BPdata %>%
rename(Extractor_name = "Name_of_the_extracting_person",
       Award_name = "Full_name_of_the_award",
       Journal_name = "Full_name_of_the_awarding_journal",
       Award_description = "Paste_the_information_text_describing_the_award",
       Eligible = "Confirm_eligibility_of_the_award",
       Awarding_journal = "Full_name_of_the_awarding_journal",
       Awarding_society = "Full_name_of_the_awarding_society",
       Awarding_other = "Full_name_of_the_awarding_publisher/other_body",
       Career_stage = "Target_career_stage_of_eligible_applicants",
       Flexible_eligibility = "Flexibility_of_the_eligibility_criteria",
       Assessors_transparency = "Assessor_transparency",
       Open_science = "Valuing_Open_Science",
       Self_nomination = "Self-nomination_allowed"
       )

##check for rows with empty "Scimago Subject Area" values
table(is.na(BPdata$Scimago_Subject_Area)) #4 rows from pilot screening
```

```
##
## FALSE  TRUE
##   264     4
```

```
##remove rows with pilot extractions and empty "Scimago Subject Area" values
BPdata <- BPdata[!is.na(BPdata$Scimago_Subject_Area), ]

#remove awards that were duplicate-extracted and excluded at extraction stages
BPdata <- BPdata[BPdata$Row_excluded == 0, ]
dim(BPdata) #222 rows left
```

```
## [1] 222  54
```

```
## [1] 222  54
##NOTE: total of 41 rows  removed as duplicates
##(19 awards were doubly- or triply-extracted)

#create new variable for awards with description or without
BPdata$Descr_available <- fct_collapse(BPdata$Open_science,
  available = c("no", "yes"),
  "not available" = "not available"
)
```

Load winners-level data set and an associated meta-data table:

```
#load individual winners data
BPindiv <- read_csv(here("data", "BP_awards_lists_SHAREDCOPY - indiv_winners_20230915.csv"),
                    skip = 1) #load individual winners data
```

```
## New names:
## Rows: 1079 Columns: 23
## ── Column specification
## ──────────────────────────────────────────────────────────── Delimiter: "," chr
## (21): award_SA, award_name, award_link, individual, awardee_name, awarde... dbl
## (2): record_count, award_year
## ℹ Use `spec()` to retrieve the full column specification for this data. ℹ
## Specify the column types or set `show_col_types = FALSE` to quiet this message.
## • `` -> `...23`
```

```
#load individual winners meta-data
BPindiv_meta <- read_csv(here("data", "BP_awards_lists_SHAREDCOPY - indiv_winners_meta-data.csv"),
                         skip = 1) #load individual winners data
```

```
## Rows: 22 Columns: 3
## ── Column specification ────────────────────────────────────────────────────────
## Delimiter: ","
## chr (3): column name, description, data type [options]
##
## ℹ Use `spec()` to retrieve the full column specification for this data.
## ℹ Specify the column types or set `show_col_types = FALSE` to quiet this message.
```

Load country-level productivity SCImago data set:

```
#load SCImago 2021 country productivity (documents) data downloaded from https://www.scimagojr.com/countryrank.php?
COprod <- read_csv(here("data", "scimagojr country rank 2021.csv"), skip = 0) #load data
```

```
## Rows: 235 Columns: 9
## ── Column specification ────────────────────────────────────────────────────────
## Delimiter: ","
## chr (2): Country, Region
## dbl (7): Rank, Documents, Citable documents, Citations, Self-citations, Cita...
##
## ℹ Use `spec()` to retrieve the full column specification for this data.
## ℹ Specify the column types or set `show_col_types = FALSE` to quiet this message.
```

```
#create meta-data with columns:
COprod_meta <- tibble("column name" = names(COprod),
                      "description" = c("Rank of a given country across all Scimago Subject Areas.",
                                        "Country name.",
                                        "Country location classification into one of the eight SCImago Major World 
                                        "Number of documents associated with a goven country published in 2021.",
                                        "Number of citable documents associated with a goven country published in 2
                                        "Number of citations to documents associated with a goven country published
                                        "Number of self-citations to documents associated with a goven country publ
                                        "Number of citations per document associated with a goven country published
                                        "H index of a country in 2021." ),
                      "data type [options]" = c("integer",
                                                "free text",
                                                "free text",
                                                "integer",
                                                "integer",
                                                "integer",
                                                "integer",
                                                "numeric",
                                                "integer"
                                                ))
```

Load aythor contributorship data set:

```
#load people and tasks log table country productivity
AUcontr <- read_csv(here("data", "BP_awards_lists_SHAREDCOPY - people_tasks log.csv"), skip = 0, n_max = 19) #load 
```

```
## Rows: 19 Columns: 18
## ── Column specification ────────────────────────────────────────────────────────
## Delimiter: ","
## chr  (2): name, author position
## dbl (16): piloting, documentation, register plan, checks and feedback, journ...
##
## ℹ Use `spec()` to retrieve the full column specification for this data.
## ℹ Specify the column types or set `show_col_types = FALSE` to quiet this message.
```

```
#simplify column names
names(AUcontr) <- c("name",
                    "piloting_documentation",
                    "register_plan",
                    "plan_checks_feedback",
                    "journal_screening_(SA)",
                    "journal_screening_checks_(SA)",
                    "awards_data_extraction_(SA)",
                    "awards_data_extraction_checks_(SA)",
                    "awardees_data_extraction_(SA)",
                    "awardees_data_extraction_checks (SA)",
                    "data_wrangling",
                    "data_wrangling_checks",
                    "draft_writing",
                    "draft_checks_feedback",
                    "finalising_manuscript",
                    "finalising_manuscript_checks_feedback",
                    "author_order",
                    "author_position")

#create meta-data for all columns:
AUcontr_meta <- tibble("column name" = names(AUcontr),
                       "description" = c("Name of the author",
                                         "Contribution to project piloting and documentation (1 = yes, 0 = no)",
                                         "Contribution to registering the project plan (1 = yes, 0 = no)",
                                         "Contribution to checks and feedback on the project plan (1 = yes, 0 = no)"
                                         "Contribution to journal screening and awards shortlisting (number of Subje
                                         "Contribution to cross-checking of journal screening and awards shortlistin
                                         "Contribution to awards data extractions (number of Subject Areas)",
                                         "Contribution to cross-checking of awards data extractiosn (number of Subje
                                         "Contribution to gender and affiliation data extraction (number of Subject 
                                         "Contribution to cross-checking of gender and affiliation data extraction (
                                         "Contribution to data cleaning and pilot analyses (1 = yes, 0 = no)",
                                         "Contribution to cross-checking ofdata cleaning and pilot analyses (1 = yes
                                         "Contribution to draft writing (1 = yes, 0 = no)",
                                         "Contribution to draft - checks and feedback (1 = yes, 0 = no)",
                                         "Contribution to finalising draft (1 = yes, 0 = no)",
                                         "Contribution to finalising draft - checks and feedback (1 = yes, 0 = no)",
                                         "Numeric position on the authorship list",
                                         "Position on the authorship list (first, middle, last)"),
                       "data type [options]" = c("text",
                                                 "integer",
                                                 "integer",
                                                 "integer",
                                                 "integer",
                                                 "integer",
                                                 "integer",
                                                 "integer",
                                                 "integer",
                                                 "integer",
                                                 "integer",
                                                 "integer",
                                                 "integer",
                                                 "integer",
                                                 "integer",
                                                 "integer",
                                                 "integer",
                                                 "text"
                                                 ))
```

#### Meta-data tables.

##### Table S1.

*Meta-data for the SCImago journal rankings by Subject Areas dataset.*

```
## making a html table:
kable(SAmeta, "html") #%>%
```

| column name | description | data type [options] |
|---|---|---|
| Subject_area | Name of the Scimago Subject Area (SA) representing 27 major thematic categories. | categorical [one of 27 Subject Areas] |
| Rank | Rank of a given Journal within a given Scimago Subject Area. | integer |
| Sourceid | Unique ID for the journal. | integer |
| Title | Name of the journal. | free text |
| Reviewer_name | Name of a person who reviews the journal trying to find associated eligible awards. | free text |
| Link_journal | Year when award was won/announced for a give individual winner, as reported on the award page or other documents with winner information. | web link |
| Society_name | Name of a person who won the award, as reported on the award page or other documents with winner information. | free text |
| Has_relevant_award | Gender assigned to an individual past awardee names using information from pronouns, images, names) available on award websites, personal or institutional websites, or from first names. | categorical [yes; no] |
| Link_award | Main source of information for assigning gender. Order of preference: pronouns, photo, name. | web link |
| Notes_award | Affiliation institution (usually university) assigned to a winner when the award was received using information available on award websites, personal or institutional websites, or information in the awarded paper. | free text |
| Checker_name | Name of the person who cross-checked a given row of screening data. | free text |
| Check_pass | Main source of the winner's affiliation information (institution, country) for the year when the award was received or announced. | categorical [yes; no] |
| Check_notes | Any comments for awardee, e.g. award sub-category, or link to additional info from Internet searches. | free text |
| Award_name_extract | Whether the award is shared with other authors of the same article. | categorical [yes; no] |
| Extract | Whether a bio/profile of the winner is shown on the award page/announcement (with more extended information than affiliation information only). | categorical [yes; no] |
| Award_extractor_name | Name of the person who extracted award data . | categorical [yes; no] |
| Extracted | Whether a given award has been extracted. | categorical [yes; no] |
| Comments | Anny comments regarding screened and extracted data in a given row. | free text |
| Archived | DOI of the winning paper. DOI code in a format starting with "10.". | DOI |
| Issn | ISSN number of teh journal | free text |
| SJR | SCImago Journal Rank indicator expresses the average number of weighted citations received in the selected year by the documents published in the selected journal in the three previous years, i.e. weighted citations received in year X to documents published in the journal in years X-1, X-2 and X-3 | free text |
| SJR Best Quartile | Anny comments regarding extracted data in a given row. | categorical [Q1; Q2; Q3; Q4] |
| H index | The h index expresses the journal's number of articles (h) that have received at least h citations. It quantifies both journal scientific productivity and scientific impact and it is also applicable to scientists, countries, etc. | number |
| Total Docs. (2021) | Output of the selected period. All types of documents are considered, including citable and non citable documents. | integer |
| Total Docs. (3years) | Published documents in the three previous years (selected year documents are excluded), i.e.when the year X is selected, then X-1, X-2 and X-3 published documents are retrieved. All types of documents are considered, including citable and non citable documents. | integer |
| Total Refs. | Total number of all the bibliographical references in a journal in the selected period. | integer |
| Total Cites (3years) | Number of citations received in the seleted year by a journal to the documents published in the three previous years, –i.e. citations received in year X to documents published in years X-1, X-2 and X-3. All types of documents are considered. | integer |
| Citable Docs. (3years) | Number of citable documents published by a journal in the three previous years (selected year documents are excluded). Exclusively articles, reviews and conference papers are considered. | integer |
| Cites / Doc. (2years) | Average citations per document in a 2 year period. It is computed considering the number of citations received by a journal in the current year to the documents published in the two previous years, –i.e. citations received in year X to documents published in years X-1 and X-2. | number |
| Ref. / Doc. | References per Document is the average number of references per document in the selected year. | number |
| Country | Country where journal is indexed by Scopus. | categorical |
| Region | Eight Major World Regions in used to facilitate sectorial analyses. | categorical |
| Publisher | name of teh journal publisher. | free text |
| Coverage | Year range for which publication data is available on Scopus (?). | free text |
| Categories | Tags for relevant 309 specific subject categories according to Scopus® Classification. | categorical |

```
  #kable_styling("striped", position = "left") #%>%
  #scroll_box(width = "100%", height = "1500px")
```

##### Table S2.

*Meta-data for the Best Paper awards dataset.*

```
## making a html table:
kable(BPmeta, "html") #%>%
```

| column name | description | data type [options] |
|---|---|---|
| Timestamp | System-recorded data and time of when the given row of data was submitted via the Google Form. | time |
| Name of the extracting person | Name of the extracting person: Please use your name in one consistent form (should be matching the name you gave us on EOI and recorded on GoogleSheet for award screening). It is best to always use your first and last name, to avoid confusion. | free text |
| Scimago Subject Area | Scimago Subject Area: Record name of the Scimago Subject Area (SA) you are doing extractions for - as in the Subject_area column in the GoogleSheet with journal and awards lists. Should be exactly as used in the original list of Subject Areas. | categorical [one of 27 Subject Areas] |
| Full name of the award | Full name of the award: Record specific award name, as in the Award_name_extract column in the GoogleSheet with journal and awards lists. | free text |
| Source of the information on the award | Source of the information on the award: Ideally, enter a link to an online page/document with information on award eligibility and assessment criteria. If not available, could be also a link to any document describing the award. If you cannot find any information about the award enter "NA". You can paste more than one link separated by ";". | weblink |
| Paste the information text describing the award | Paste the information text describing the award: Ideally, copy and paste the section of the online page/document with information on award eligibility and assessment criteria only. This description text will be used for text-mining analyses. Do not copy and paste list of past winners or generic information about the journal or society. Do not copy and paste any images. If you cannot find any information about the award enter "NA". | free text |
| Confirm eligibility of the award | Confirm eligibility of the award: · This award is for a single published research contribution in a format of a research article (theses / dissertation are allowed if published as a journal article). · This award is managed by a journal, publisher, or a learned society. · This award is NOT discontinued or restricted to applicants from underrepresented groups, e.g., women-only / minorities-only. Options: yes (eligible award), no (not eligible - make note on the reasons below and enter NA to the following compulsory questions, then submit), unclear (make note on the reasons below) | categorical [yes; no; unclear] |
| Comment on the award eligibility | Comment on the award eligibility: Any comments, e.g. reasons for excluding this award. | free text |
| Full name of the awarding journal | Full name of the awarding journal: Record if award is associated with a specific journal. Enter "NA" if not associated with a specific journal. | free text |
| Full name of the awarding society | Full name of the awarding society: Record if award is associated with a specific society. Enter "NA" if not associated with a specific society. | free text |
| Full name of the awarding publisher/other body | Full name of the awarding publisher/other body: Record if award is associated with a specific publisher (group of journals) or awarding organisation other than learned society or journal. Enter "NA" if not associated with a specific publisher. | free text |
| Comment on the awarding body | Comment on the awarding body: Any comments, e.g. if more than one journal considered for the award. | free text |
| Award cash | Award cash: Code whether there is any cash coming with the award. - Select "yes" if award description mentions any cash given to the winners. - Select "no" if award description does not mention any cash given to the winners. Travel awards, publishing discounts, plenary talks, etc. do not count as cash awards. - Select "not available" if there is no page/document describing the award (e.g. only the list of winners is publicly published). NOTE: In the next question ("Inclusivity phrasing") copy and paste relevant text from the award document or make a note if no such document/information is available. Options: yes, no, not available. | free text |
| [illegible] | Comment on the award cash: Add comments on cash amount, if provided, and note any other | [illegible] |

| | | |
|---|---|---|
| Comment on the award cash | [illegible] benefits winners receive, as relevant (e.g., travel awards, publishing discounts, plenary talks) | free text |
| Target career stage of eligible applicants | Target career stage of eligible applicants, as stated in the award information: As stated in the award information. More than one choice is possible. If it is for any paper in a journal/s, select "any career stage". Options: student, early-career, mid-career, any career stage, unclear. | categorical [student; early-career; mid-career; any career stage; unclear] |
| Comment on the career stage | Comment on the career stage: Any comments, e.g. paste the phrasing of who is eligible for the award. | free text |
| Flexibility of the eligibility criteria | Flexibility of the eligibility criteria: Code whether explicitly allowing for career interruptions in eligibility timeframes: - Select "not applicable" if description only states that published research has to be performed while studying for a degree OR if there is no eligibility limit in terms of years since an event (e.g. a PhD or author's age. - Select "yes" if the description says that there is an eligibility limit of years after a degree to apply for the. award and this limit can be extended in special cases". - Select "no" if the description says that there is an eligibility limit of years after a degree to apply for the award but it does not mention that this limit can be extended in special cases. - Select "not available" if there is no page/document describing the award (e.g. only the list of winners is publicly published). NOTE: In the next question ("Eligibility phrasing") copy and paste relevant text from the award document or make a note if no such document/information is available. Options: yes, no, not applicable, not available. | categorical [yes; no; not applicable; not available] |
| Eligibility phrasing | Eligibility phrasing: Wording of the eligibility criteria in relation to career stage in the relevant documentation, if available. | free text |
| Inclusivity statement | Inclusivity statement: Code whether underrepresented/minority groups are encouraged to apply for the award (this does not mean that the award is restricted to underrepresented groups, e.g., women-only) or award information includes declarations of commitment to equity / diversity / inclusivity (EDI): - Select "yes" if award description mentions underrepresented/minority groups or EDI. - Select "no" if award description does not mention anything about underrepresented/minority groups or EDI. - Select "not available" if there is no page/document describing the award (e.g. only the list of winners is publicly published). NOTE: Extract this information from award descriptions only - do not include information from other documents not directly related to the award, e.g. journal/society/institutional policies or mission statements. In the next question ("Inclusivity phrasing") copy and paste relevant text from the award document or make a note if no such document/information is available. | categorical [yes; no; not available] |
| Inclusivity phrasing | Inclusivity phrasing: Wording of the inclusivity statement in the relevant documentation, if available. | free text |
| Assessor transparency | Assessor transparency: Code whether information is provided on who will be conducting assessments of candidate papers (for example, that editors-in-chief will be deciding): - Select "yes" if information is provided, e.g. names, or stating that journal editors will be doing this, or mentioning specific existing committee that is described somewhere else (e.g. on a society webpage). - Select "no" if description does not mention any information on the assessors. - Select "not available" if there is no page/document describing the award (e.g. only the list of winners is publicly published). In the next question ("Assessor phrasing") copy and paste relevant text from the award document or make a note if no such document/information is available. | categorical [yes; no; not available] |
| Assessor phrasing | Assessor phrasing: Wording of the information on who will be conducting the assessments, if available. | free text |
| Process transparency | Process transparency: Code whether breakdown of the applicants / candidates by gender or geographic region is publicly available: - Select "yes" if such information is provided (e.g. on a society webpage). - Select "no" if such information is not provided. - Select "not available" if there is no page/document describing the award (e.g. only the list of winners is publicly published). NOTE: In the next question ("Process phrasing") copy and paste relevant text from the award document or make a note if no such document/information is available. Options: yes, no, not available. | categorical [yes; no; not available] |
| Process phrasing | Process phrasing: Wording of the information on the transparency of the process, e.g. if breakdown of the applicants / candidates by gender or geographic region is publicly available. | free text |
| Feedback availability | Feedback availability: Code whether award information included an offer of constructive feedback for unsuccessful applicants: - Select "yes" if such feedback can be provided (e.g. on request). - Select "no" if such feedback is not provided or not mentioned. - Select "not available" if there is no page/document describing the award (e.g. only the list of winners is publicly published). NOTE: In the next question ("Feedback phrasing") copy and paste relevant text from the award document or make a note if no such document/information is available. Options: yes, no, not available. | categorical [yes; no; not available] |
| Feedback phrasing | Feedback phrasing: Wording of the information on whether/how feedback will be provided, if available. | free text |
| Criteria transparency | Criteria transparency: Code whether assessment criteria are detailed (more than one vague sentence) or vague (often one vague sentence, e.g. "assessed on innovation and novelty"): - Select "yes" if assessment criteria are more than one vague sentence (e.g., mentions things like readability of the language, sample size, performing alternative tests/analyses to check robustness of the results, sharing data or code, registering the study/protocol - whatever is relevant to a given research field). - Select "no" if assessment criteria are only one vague sentence (e.g. award is for "best paper", "outstanding contribution", "excellent research", "novel insights", etc.). - Select "not available" if there is no page/document describing the award (e.g. only the list of winners is publicly published). NOTE: In the next question ("Criteria phrasing") copy and paste relevant text from the award document or make a note if no such document/information is available. Options: yes, no, not available. | categorical [yes; no; not available] |
| Criteria phrasing | Criteria phrasing: Wording of the information on the assessment criteria, if available. | free text |
| Valuing Open Science | Valuing Open Science: Code whether any Open Science practices (data, code, materials sharing, preregistration, transparency of reporting, etc.) are explicitly included in the assessment criteria: - Select "yes" if award description mentions valuing any of the Open Science practices (e.g., sharing data or code, registering the study/protocol, performing replication studies - whatever is relevant to a given research field). - Select "no" if the award description does not mention valuing any of the Open Science practices. - Select "not available" if there is no page/document describing the award (e.g. only the list of winners is publicly published). NOTE: In the next question ("Valuing Open Science phrasing") copy and paste relevant text from the award document or make a note if no such document/information is available. Options: yes, no, not available. | categorical [yes; no; not available] |
| Valuing Open Science phrasing | Valuing Open Science phrasing: Wording of the information on the assessment criteria valuing Open Science practices, if available. | free text |
| Self-nomination allowed | Self-nomination allowed: Code whether candidates can self-nominate for the award. - Select "yes" if the award description explicitly mentions that candidates can self-nominate. - Select "no" if the award description states that candidates/papers need to be nominated by somebody else or does not explicitly mention that candidates can self-nominate. - Select "not available" if there is no page/document describing the award (e.g. only the list of winners is publicly published). NOTE: In the next question ("Self-nomination phrasing") copy and paste relevant text from the award document or make a note if no such document/information is available. Options: yes, no, not available, | categorical [yes; no; not available] |
| Self-nomination phrasing | Self-nomination phrasing: Wording of the information on how to nominate (e.g., using a checkbox on the manuscript submission form), if available. | free text |
| Letter required | Letter required: Code whether candidates are required to provide nomination/recommendation letter/s: - Select "yes" if award description explicitly mentions that candidates have to provide nomination or support letters from others. - Select "no" if the award description states that candidates/papers do not need to provide letters from somebody else or does not explicitly mention anything about such letters. - Select "not available" if there is no page/document describing the award (e.g. only the list of winners is publicly published). NOTE: In the next question ("Letter requirement phrasing") copy and paste relevant text from the award document or make a note if no such document/information is available. Options: yes, no, not available. | categorical [yes; no; not available] |
| Letter requirement phrasing | Letter requirement phrasing: Wording of the information on the requirement for written nominations / reference letters, if available. | free text |
| Awardee list source | Awardee list source: Source of the information on the past winners - paste a link to a webpage, file name, personal information, etc., showing a list of past winners (any year span). Write "not available" if no such list exists. | weblink |
| Awardee list most recent year | Awardee list most recent year: The most recent year for which information on past awardees is available (enter whole number, e.g. 2022). Leave empty if no such list exists. | integer |
| Awardee list earliest year | Awardee list earliest year: The earliest year for which information on past awardees is available (enter whole number, e.g. 1999). Leave empty if no such list exists. | integer |
| Comments on awardees list | Comments on awardees list: Any comments for awardees list, e.g. if Internet searches might be needed to locate announcements with the names of the past winners. | free text |
| Comments_general | Comments_general: Any other notes and comments on issues, assumptions, or seeking additional information from the award committees / contacts. | free text |
| Checked | Whether a given row of extracted data has been cross-checked. | categorical [yes; no] |
| Checker_name | Name of the person who cross-checked a given row of extracted data. | free text |
| Checker_comments | Anny comments regarding extracted data in a given row. | free text |
| Row_excluded | Whether a given row of extracted data has been cross-checked. | integer [1 = yes; 0 = no] |
| Award_individual | Whether a given award recognises individual (selected) authors rather than all authors of a winning paper. | categorical [yes; no] |
| Award_impact_metrics_mentioned | Whether award criteria mention impact metrics (number of citations, downloads) as basis of winner selection. | categorical [yes; no] |
| Award_impact_metrics_only | Whether impact metrics (number of citations, downloads) ar the sole basis of winner selection. | categorical [yes; no] |
| Award_impact_metrics_comment | Quote relevent sections of award description if impact metrics are mentioned. Add any comments regarding award criteria and impact metrics. | free text |
| Award_contact_provided | Whether specific contact information (email) is provided on the award webpage or publicly available application/description documents. | categorical [yes; no] |
| Award_integrity_mentioned | Whether award description states how potential conflictes of interests will be managed | categorical [yes; no; not available] |
| Award_integrity_comment | Quote relevent sections of award description on how potential conflicts of interests will ba managed. | free text |
| Award_cash_max_USD_pperson | The maximum reported amount of cash awarded per winning paper, recalculated online into USD. | Integer |
| N_winners_extracted | Number of individual winners from years 2001-2022 extracted for this awards (in indiv_winners data set). | Integer |
| Archived_files | Whether award website and over information have been archived as pdf files in the dedicated project folder. | categorical [yes; no] |
| Comment_extraction | Anny comments regarding extracted data in a given row. | free text |

```
  #kable_styling("striped", position = "left") #%>%
  #scroll_box(width = "100%", height = "2500px")
```

Table S3.

*Meta-data for the individual winners dataset.*

```
## making a html table:
kable(BPindiv_meta, "html") #%>%
```

| column name | description | data type [options] |
|---|---|---|
| award_SA | Name of the Scimago Subject Area (SA) you are doing extractions for - as in the Subject_area column in the GoogleSheet with journal and awards lists. Should be exactly as used in the original list of Subject Areas. | categorical [one of 27 Subject Areas] |
| record_count | Subsequent numbers for counting extracted winners within each award. | integer |
| award_name | Name of the award - as in the Award_name column in the main GoogleSheet with awards main data extraction. | free text |
| award_link | Weblink to the main Internet page with award description / information. | weblink |
| individual | Whether an award is individual or article-focused. Individual awards are defined as awards that are usually not shared equally between all article authors. For example the award names as a winner only the ECR authors or only the first author (unless there are more than one ECR/first authors who contributed equally). | categorical [yes; no] |
| award_year | Year when award was won/announced for a give individual winner, as reported on the award page or other documents with winner information. | integer |
| awardee_name | Name of a person who won the award, as reported on the award page or other documents with winner information. | free text |
| awardee_gender | Gender assigned to an individual past awardee names using information from pronouns, images, names) available on award websites, personal or institutional websites, or from first names. | categorical [F = female; M = male; ? - unassigned] |
| gender_source | Main source of information for assigning gender. Order of preference: pronouns, photo, name. | categorical [pronouns; photo; name] |
| affiliation_institution | Affiliation institution (usually university) assigned to a winner when the award was received using information available on award websites, personal or institutional websites, or information in the awarded paper. | free text |
| affiliation_country | Affiliation country assigned to a past winner for the year when the award was received using information available on award websites, personal or institutional websites, or information in the awarded paper. | free text |
| affiliation_info_source | Main source of the winner's affiliation information (institution, country) for the year when the award was received or announced. | categorical [award page; award announcement; article; other] |
| awardee_comment1 | Any comments for awardee, e.g. award sub-category, or link to additional info from Internet searches. | free text |
| shared | Whether the award is shared with other authors of the same article. | categorical [yes; no] |
| awardee_profile_shown | Whether a bio/profile of the winner is shown on the award page/announcement (with more extended information than affiliation information only). | categorical [yes; no] |
| awardee_photo_shown | Whether a photo of the winner is shown on the award page/announcement. | categorical [yes; no] |
| awardee_comment2 | Any other comments on the awardee or awarded paper, e.g. award sub-category, or link to additional info from Internet searches. | free text |
| awarded_paper_ref | Bibliographic reference of the winning paper. | free text [reference in any long format] |
| awarded_paper_doi | DOI of the winning paper. DOI code in a format starting with "10.". | DOI |
| checked | Whether a given row of extracted data has been cross-checked. | categorical [yes; no] |
| checker_name | Name of the person who cross-checked a given row of extracted data. | free text |
| checker_comment | Anny comments regarding extracted data in a given row. | free text |

```
  #kable_styling("striped", position = "left") #%>%
  #scroll_box(width = "100%", height = "1500px")
```

Table S4.

*Meta-data for the author contributions dataset.*

```
## making a html table:
kable(AUcontr_meta, "html") #%>%
```

| column name | description | data type [options] |
|---|---|---|
| name | Name of the author | text |
| piloting_documentation | Contribution to project piloting and documentation (1 = yes, 0 = no) | integer |
| register_plan | Contribution to registering the project plan (1 = yes, 0 = no) | integer |
| plan_checks_feedback | Contribution to checks and feedback on the project plan (1 = yes, 0 = no) | integer |
| journal_screening_(SA) | Contribution to journal screening and awards shortlisting (number of Subject Areas) | integer |
| journal_screening_checks_(SA) | Contribution to cross-checking of journal screening and awards shortlisting (number of Subject Areas) | integer |
| awards_data_extraction_(SA) | Contribution to awards data extractions (number of Subject Areas) | integer |
| awards_data_extraction_checks_(SA) | Contribution to cross-checking of awards data extractiosn (number of Subject Areas) | integer |
| awardees_data_extraction_(SA) | Contribution to gender and affiliation data extraction (number of Subject Areas) | integer |
| awardees_data_extraction_checks (SA) | Contribution to cross-checking of gender and affiliation data extraction (number of Subject Areas) | integer |
| data_wrangling | Contribution to data cleaning and pilot analyses (1 = yes, 0 = no) | integer |
| data_wrangling_checks | Contribution to cross-checking ofdata cleaning and pilot analyses (1 = yes, 0 = no) | integer |
| draft_writing | Contribution to draft writing (1 = yes, 0 = no) | integer |
| draft_checks_feedback | Contribution to draft - checks and feedback (1 = yes, 0 = no) | integer |
| finalising_manuscript | Contribution to finalising draft (1 = yes, 0 = no) | integer |
| finalising_manuscript_checks_feedback | Contribution to finalising draft - checks and feedback (1 = yes, 0 = no) | integer |
| author_order | Numeric position on the authorship list | integer |
| author_position | Position on the authorship list (first, middle, last) | text |

```
  #kable_styling("striped", position = "left") #%>%
  #scroll_box(width = "100%", height = "1500px")
```

### Supplementary Results

#### Characteristics of SCImago Subject Area (SA) journal lists and screening data.

Plot Number of journals per SCImago Subject Area:

```
##count journals per SA
# count_j_perSA <- SAdata %>%
#     count(Subject_area) %>%
#     arrange(desc(n))

#table(SAdata$Subject_area)
#min(table(SAdata$Subject_area))
#max(table(SAdata$Subject_area))

#all SA as a simple barplot
SAdata %>%
  #filter(!is.na(Subject_area)) %>%
  count(Subject_area) %>%
  arrange(Subject_area) %>%
  ggplot(aes(x = reorder(Subject_area, desc(Subject_area)), y = n)) +
  geom_bar(stat = "identity", position = position_dodge(0.9)) +
  coord_flip()
```

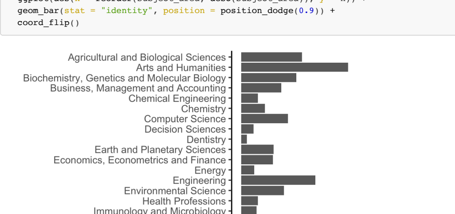


Figure S2.

*Counts of journal Titles per SCImago ranking lists by Subject Area.*

NOTE: The numbers of journals per Subject Area vary widely reflecting the differences in sizes of different research disciplines. Thus, random sampling of journals across Subject Areas could result in uneven representation of disciplines, potrentially biasing the findings towards the largest disciplines. By sampling similar number of journals with similar levels of "ranking" in each Subject Area, we were able to get a rough estimate of the frequency of the awards across disciplines.

#### Overlap of journals across SCImago Subject Areas.

```
# length(unique(SAdata$Title)) #number of unique journals
# length(unique(SAdata$Title))/length(SAdata$Title) #60% unique

#reduce to top 100 from each SA
SAdata %>%
  #filter(!is.na(Subject_area)) %>%
  group_by(Subject_area) %>%
  top_n(100, Title) -> SAdata100

#count number of duplicated journal titles across all SA top 100 journal titles:
#sum(duplicated(SAdata100$Title)) #644 duplicated journal Titles (out of 2700)
#sum(duplicated(SAdata100$Title))/length(SAdata100$Title) #24% duplicated journal Titles (out of 2700)
```

Across the top 100 journals in each Subject Area ranking list there are 644 duplicated journal titles (24%). This indicates that one journal can often be classified into one or more related Subject Areas (e.g. "Trends in Cognitive Sciences" is included both in Neuroscience and Psychology Subject Areas). This overlap implies that the related awards will also often be relevant to more than one Subject Area. Thus, after the screening step, we checked for duplicate inclusions and data extractions and assigner each award to only one Subject Area for the analyses to avoid double-counting.

#### Award-level dataset characteristics.

Overall summary:

Total number of unique names of awards: 222.
Total number of unique names of awards: 27.

Total number of awards without award description: 11 (5%).

Total number of individual winners records extracted: 1079.
Total number of unique names of individual winners records extracted: 1065 (99%).

Awarding bodies:

Total number of awards associated with a journal: 176 (83%).
Total number of awards associated with a society: 144 (68%).
Total number of awards associated with other awarding body: 39 (18%).
Note: one oward could be associated with more than one category from above.

Overlap of awarding journals across SA: 0 (0%).
Overlap of of awarding societies across SA: 18 (9%).
Overlap of awarding other bodies across SA: 18 (9%).

```
##### Initial data sanity checks (Note: code chunk not evaluated on knitting)

##Check award names - same award could have been extracted in different SA, or different awards have same names
# table(duplicated(BPdata$Award_name)) #0 duplicated, 222 unique

## Simple counts

# #eligible awards per SA
# count_awards_perSA <- BPdata %>%
#     count(Scimago_Subject_Area) #%>%
#     #arrange(desc(n))

# #count number of unique awards per year:
# BPindiv %>%
#   #filter(!is.na(affiliation_country)) %>%
#   group_by(award_year) %>%
#   summarise(count = n_distinct(award_name))

##How many awards per Scimago_Subject_Area
# table(BPdata$Scimago_Subject_Area) #note many SA with <10 extracted awards
#- this is because of many journals being shared between SA

table(BPdata$Awarding_journal == "NA") #46 not associated with a journal
table(BPdata$Awarding_society == "NA") #78 not associated with a learned society
table(BPdata$Career_stage, useNA = "always")
table(BPdata$Award_individual, useNA = "always") #66 yes
table(BPdata$Flexible_eligibility, useNA = "always") #only 5 - "yes"
#View(BPdata[BPdata$Flexible_eligibility == "yes", ]) #see rows with "yes"

#two-way table for the Career_stage and Flexible_eligibility
table(BPdata$Career_stage, BPdata$Flexible_eligibility, useNA = "always") #most "not applicable" is for "any career

#two-way table for the Flexible_eligibility and Award_individual
table(BPdata$Flexible_eligibility, BPdata$Award_individual, useNA = "always") #flexible ones are for ECRs only

#two-way table for the Career_stage and Award_individual
table(BPdata$Career_stage, BPdata$Award_individual, useNA = "always") #individual are mostly for ECRs
#BPdata$Award_name[BPdata$Career_stage == "unclear"]
#BPdata$Award_name[BPdata$Career_stage == "early-career" & BPdata$Award_individual == "no"]
#BPdata$Award_name[BPdata$Career_stage == "early-career" & BPdata$Award_individual == "no"]
#length(BPdata$Award_name[BPdata$Career_stage == "any career stage" & BPdata$Award_individual == "no"]) #150

#three-way table
table(BPdata$Career_stage, BPdata$Flexible_eligibility, BPdata$Award_individual) #most "not applicable" is for "any

length(BPdata$Award_name[BPdata$Award_individual == "no" &
                           BPdata$Career_stage == "any career stage" &
                           BPdata$Flexible_eligibility == "not applicable"]) #147 not applicable flexibility

length(BPdata$Award_name[BPdata$Award_individual == "no" &
                           BPdata$Career_stage == "any career stage" &
                           BPdata$Flexible_eligibility == "not available"]) #3 with no description

length(BPdata$Award_name[BPdata$Award_individual == "no" &
                           BPdata$Career_stage == "any career stage" &
                           BPdata$Flexible_eligibility == "yes"]) #0 with flexibility

length(BPdata$Award_name[BPdata$Award_individual == "yes" &
                           BPdata$Career_stage == "any career stage"]) #28 not limited to specific career stage

length(BPdata$Award_name[BPdata$Award_individual == "yes" &
                           BPdata$Flexible_eligibility == "not applicable"]) #38 not applicable flexibility

length(BPdata$Award_name[BPdata$Award_individual == "yes" &
                           BPdata$Flexible_eligibility == "not available"]) #2 with no description

length(BPdata$Award_name[BPdata$Award_individual == "yes" &
                           BPdata$Flexible_eligibility == "yes"]) #5 with flexibility

table(BPdata$Inclusivity_statement, useNA = "always") #only 2 with "yes"

#BPdata$Award_name[BPdata$Inclusivity_statement == "yes"] #see rows with "yes"

table(BPdata$Assessors_transparency, useNA = "always") #114 "yes"
#View(BPdata[BPdata$Assessors_transparency == "yes", ]) #see rows with "yes"

# sum(str_count(tolower(BPdata$Assessor_phrasing), "editor"), na.rm = TRUE) #162 - counting all mentions
# sum(str_detect(BPdata$Assessor_phrasing, "editor"), na.rm = TRUE) #36 - counting once per award

table(BPdata$Process_transparency, useNA = "always") #only 2 "yes"
#BPdata$Award_name[BPdata$Process_transparency == "yes"] #see award name with "yes"
#View(BPdata[BPdata$Process_transparency == "yes",]) #see award name with "yes"

table(BPdata$Self_nomination, useNA = "always") #28 yes
#View(BPdata[BPdata$Self_nomination == "yes", ]) #see rows with "yes"

table(BPdata$Letter_required, useNA = "always") #38 yes
#View(BPdata[BPdata$Letter_required == "yes", ]) #see rows with "yes"

table(BPdata$Letter_required, BPdata$Self_nomination, useNA = "always") #two-way table
dim(BPdata[BPdata$Letter_required == "yes" & BPdata$Self_nomination == "yes", ])[1] #mentioning letter and self-nom
dim(BPdata[BPdata$Letter_required == "no" & BPdata$Self_nomination == "no", ])[1] #not mentioning letter and self-n

table(BPdata$Feedback_availability, useNA = "always") #0 "yes"

table(BPdata$Award_contact_provided, useNA = "always") #38 "yes"

table(BPdata$Criteria_transparency, useNA = "always") #40 yes
#View(BPdata[BPdata$Criteria_transparency == "yes", ]) #see rows with "yes"

table(BPdata$Award_impact_metrics_mentioned, useNA = "always") #21 "yes"

table(BPdata$Award_impact_metrics_only, useNA = "always") #8 "yes"

table(BPdata$Open_science, useNA = "always") #1 "yes"
#View(BPdata[BPdata$Open_science == "yes", ]) #see rows with "yes": Social Sciences: Review Of Research Award  by A

table(BPdata$Awardee_list_most_recent_year, useNA = "always") #a few <2020 and 19 NA
#View(BPdata[is.na(BPdata$Awardee_list_most_recent_year), ]) #see rows with NA

table(BPdata$Awardee_list_earliest_year, useNA = "always") #some old, 19 NA
#View(BPdata[is.na(BPdata$Awardee_list_earliest_year), ]) #see rows with NA
#hist(BPdata$Awardee_list_earliest_year)
length(BPdata$Awardee_list_earliest_year[BPdata$Awardee_list_earliest_year > 2010 & !is.na(BPdata$Awardee_list_earl
```

Figure 1A - Screening effort: count numbers of journals screened to get 10 shortlisted awards:

```
#load journal screening summary dataset:
BPscreening <- read_csv(here("data","BP_awards_lists_SHAREDCOPY - scimagojr_2021_SA.csv")) #load award screening su
```

```
## Rows: 29 Columns: 28
## ── Column specification ────────────────────────────────────────────────────────
## Delimiter: ","
## chr (22): Subject_Area, Reveiwer_name, Reviewer_email, Screening_status (ass...
## dbl  (4): Nr, N_records_imported, N_records_screened, Extraction1_N_awards_e...
## lgl  (2): Extraction1_comments, Extraction2_winners_checking_comments
##
## ℹ Use `spec()` to retrieve the full column specification for this data.
## ℹ Specify the column types or set `show_col_types = FALSE` to quiet this message.
```

```
BPscreening <- BPscreening[!is.na(BPscreening$Subject_Area), ] #remove 2 extra rows without Subject_Area
count_awards_perSA <- BPdata %>% count(Scimago_Subject_Area) #count included awards per SA
BPscreening$N_included <- count_awards_perSA$n #add included award counts per SA
BPscreening$N_excluded <- BPscreening$N_records_screened - BPscreening$N_included
BPscreening$Subject_Area <- as.factor(BPscreening$Subject_Area) #convert to factor

## reshape into long dataframe format:
BPscreening %>% select(Subject_Area, N_included, N_excluded) %>% gather(key = status, count, N_included:N_excluded)

#reorder status levels and edit labels:
BPscreening_long <- BPscreening_long %>%
  mutate(status = factor(status, levels = rev(c("N_excluded", "N_included"))))

BPscreening_long$status <- as.character(BPscreening_long$status)  #convert to character
#BPscreening_long$status[BPscreening_long$count <= 5] <- "N_included_low" #use to mark SA with 5 or less awards inc

#wacolors$san_juan #palette to be used

figure1A <- BPscreening_long %>%
    mutate(status = recode(status,
                           'N_excluded' = 'not found',
                           'N_included' = 'found and included')) %>%
    ggplot(aes(x = reorder(Subject_Area, desc(Subject_Area)),
               y = count, fill = status)) +
    geom_col(width = 0.8,
             position = position_stack(reverse = TRUE)) +
    coord_flip() +
    scale_y_continuous(breaks = c(0,25,50,75,100)) +
    theme_bw() +
    scale_fill_manual(values = c("#CA884C", "#BAAF9F")) +
    labs(x = "",
         y = "Count of screened journals",
         fill = "Award: ") +
    theme(legend.position = "top",
          legend.justification = c(0, 1),
          legend.title = element_text(size = rel(0.7)),
          legend.text = element_text(size = rel(0.7)),
          axis.title.x = element_text(size = 10, hjust = 0.5),
          plot.margin = unit(c(0, 0, 0, 0), "pt")) +
    geom_hline(yintercept = 10,
               linetype = "dotted",
               color = "black",
               linewidth = 0.7) +
    scale_x_discrete(position = "top")
```

Figure 1B - Counts of incuded awards with and without award description:

```
figure1B <- BPdata %>%
    count(Scimago_Subject_Area, Descr_available) %>%
    arrange(desc(n)) %>%
    ggplot(aes(x = reorder(Scimago_Subject_Area, desc(Scimago_Subject_Area)),
               y = n,
               fill = Descr_available)) +
    geom_col(width = 0.8,
             position = position_stack(reverse = TRUE)) +
    coord_flip() +
    scale_y_continuous(breaks = c(0,5,10)) +
    theme_bw() +
    scale_fill_manual(values = c("#3A5775", "#C9DCE2")) +
    labs(x = "",
         y = "Count of included awards",
         fill = "Award description:") +
    scale_x_discrete(labels = NULL) +
    labs(x = "")  + #used to remove vertical labels, also breaks = NULL
    theme(legend.position = "top",
          legend.title = element_text(size = rel(0.7)),
          legend.text = element_text(size = rel(0.7)),
          axis.title.x = element_text(size = 10),
          plot.margin = unit(c(0, 0, 0, 0), "pt"))
```

#### Award descriptions.

Word counts per award description:

```
#add new variable with counts of words in the award description:
BPdata$Award_description_wordcount <- str_count(BPdata$Award_description, "\\w+")
#hist(str_count(BPdata$Award_description, "\\w+"), breaks = 20)

dim(BPdata[is.na(BPdata$Scimago_Subject_Area), ])[1] #no NA, but no description was coded as NA, so if 1 then chang
```

```
## [1] 0
```

```
BPdata$Award_description_wordcount[BPdata$Award_description_wordcount < 2] <- 0 #if 1 then change to 0 (no descript
length(BPdata$Award_description_wordcount[BPdata$Award_description_wordcount == 0]) #11 awards without description
```

```
## [1] 11
```

```
#dim(BPdata[BPdata$Award_description_wordcount < 100, ])[1] #descriptions have less than 100 words! (includes 11 wi
```

Number of awards with less than 100-word description: 90 (42.7% of 211 awards with descriptions).

Median length of award description: 123.

Figure 1C - Plot of award description lengths - overall:

```
figure1C <- BPdata %>%
  filter(!is.na(Award_description_wordcount)) %>%
  ggplot(aes(x = Award_description_wordcount)) +
  geom_histogram(binwidth = 20,
                 fill = "#21281D") +
  theme_bw() +
  labs(x = "Number of words in award description",
       y = "Award count") +
  scale_x_continuous(breaks = c(seq(0, 1000, by = 100))) +
  theme(legend.position = "none",
        axis.title.x = element_text(size = 10))
```

Plot description word count by Subject Area:

```
BPdata %>%
  filter(!is.na(Award_description_wordcount)) %>%
  ggplot(aes(x = Award_description_wordcount)) +
  geom_histogram(binwidth = 20,
                 fill = "#21281D") +
  theme_bw() +
  labs(x = "Number of words in award description",
       y = "Award count") +
  scale_x_continuous(breaks = c(seq(0, 1000, by = 100))) +
  theme(legend.position = "none",
        axis.title.x = element_text(size = 10)) +
  facet_wrap(~Scimago_Subject_Area,
             ncol = 3)
```

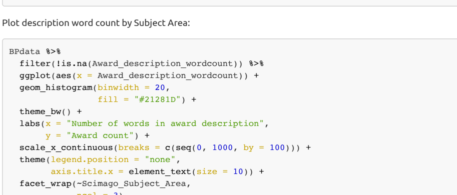


Figure S3.

Plot of award description lengths - by Subject Area.

#### Award ages

NOTE: Using the year of oldest listed past winner as a proxy.

Figure 1D - award age verall distribution:

```
#table(!is.na(BPdata$Awardee_list_earliest_year)) # values available

#plot overall - beeswarm
figure1D <- BPdata %>%
  filter(!is.na(Awardee_list_earliest_year)) %>%
  ggplot(aes(y = Awardee_list_earliest_year,
             x = Row_excluded)) +
  geom_beeswarm(color = "#21281D",
                alpha = 0.2,
                cex = 3.3) +
  coord_flip() +
  theme_bw() +
  labs(y = "Year first winner listed",
       x = "") +
  scale_x_discrete(labels = NULL) +
  labs(x = "")  + #used to remove vertical labels, also breaks = NULL
  scale_y_continuous(breaks = c(seq(1900, 2030, by = 10))) +
  theme(legend.position = "none",
        axis.title.x = element_text(size = 10)) +
  annotate("rect",
           ymin = 2001,
           ymax = 2022,
           xmin = -1.5,
           xmax = 1.5,
           alpha = .1,
           fill = "#008B00")
```

Plot award ages by Subject Area:

```
#plot by SA - beeswarm
BPdata %>%
  filter(!is.na(Awardee_list_earliest_year)) %>%
  ggplot(aes(x = reorder(Scimago_Subject_Area, desc(Scimago_Subject_Area)), y = Awardee_list_earliest_year)) +
  geom_beeswarm(color = "#21281D", alpha = 0.2) +
  coord_flip() +
  theme_bw() +
  labs(y = "Year first winner listed",
       x = "Scimago Subject Area") +
  scale_y_continuous(breaks = c(seq(1900, 2030, by = 10))) +
  theme(legend.position = "none",
        axis.title.x = element_text(size = 10))
```

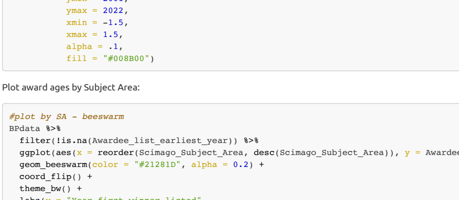


Figure S4.

*Plot of award earliest extracted winner award year - by Subject Area.*

Figure 4 - Combine 4 panels and save:

```
#assemble the panels using patchwork package
figure1 <- (figure1A + figure1B) / figure1C / figure1D +
  plot_layout(widths = c(1, 1),
              heights = c(3, 1, 1)) +
  plot_annotation(tag_levels = "A")

#ggsave(plot = figure1, here("plots", "Fig1ABCD_screening_descriptions_v1.png"), width = 18.6, height = 18, units =
#ggsave(plot = figure1, here("plots", "Fig1ABCD_screening_descriptions_v1.pdf"), width = 18.6, height = 18, units =
```

Summary by award age:
- Award year of the earliest listed winner used as a proxy: `r` min(BPdata$Awardee_list_earliest_year, na.rm = TRUE)`.
- Number of awards with the earliest listed winner known: 203.
- Number of awards with the earliest listed winner in or before 2000: 51.
- Number of awards with the earliest listed winner within 2001-2010: 45.
- Number of awards with the earliest listed winner within 2011-2023: 107.

Figure 2A - Awarding bodies:

```
# table(BPdata$Awarding_journal == "NA")
# table(BPdata$Awarding_society == "NA")
# table(BPdata$Awarding_other == "NA")

#create new variable with simple categorisation of awarding body type mentioned:
BPdata2 <- BPdata %>%
   mutate(
     Awarded_by = case_when(Awarding_journal != "NA" & Awarding_society == "NA" & Awarding_other == "NA" ~ "journal
     Awarding_journal == "NA" & BPdata$Awarding_society != "NA" & BPdata$Awarding_other == "NA" ~ "society",
     Awarding_journal == "NA" & BPdata$Awarding_society == "NA" & BPdata$Awarding_other != "NA" ~ "other",
     Awarding_journal != "NA" & BPdata$Awarding_society != "NA" & BPdata$Awarding_other == "NA" ~ "journal&society"
     Awarding_journal != "NA" & BPdata$Awarding_society != "NA" & BPdata$Awarding_other != "NA" ~ "journal&society&
     Awarding_journal != "NA" & BPdata$Awarding_society == "NA" & BPdata$Awarding_other != "NA" ~ "journal&other",
     Awarding_journal == "NA" & BPdata$Awarding_society != "NA" & BPdata$Awarding_other != "NA" ~ "society&other",
     TRUE ~ "none")
   )

#table(BPdata$Awarded_by)
```

```
#create new variable for awards with description or without
BPdata2 %>%
  select(Award_name, Awarding_journal, Awarding_society, Awarding_other) %>%
  mutate(
    across(c(Awarding_journal, Awarding_society, Awarding_other), ~ if_else(. == "NA", FALSE, TRUE))
  ) -> BPdata2

#make upset plot using library(ggupset)
figure2A <- BPdata2 %>%
  column_to_rownames(var = "Award_name") %>%
  as_tibble(rownames = "Award_name") %>%
  gather(Body, Award, -Award_name) %>%
  filter(Award) %>%
  select(-Award) %>%
  mutate(Body = recode(Body,
                       'Awarding_journal' = 'journal',
                       'Awarding_society' = 'society',
                       'Awarding_other' = 'other')) %>%
  group_by(Award_name) %>%
  summarize(Awarding_bodies = list(Body)) %>%
  ggplot(aes(x = Awarding_bodies)) +
    geom_bar(fill = "#21281D", width = 0.8) +
    scale_y_continuous(limits = c(0, 200)) +
    scale_x_upset(order_by = c("freq")) +
    theme_combmatrix(combmatrix.panel.striped_background = FALSE,
                     combmatrix.panel.point.color.fill = "#21281D",
                     combmatrix.panel.line.size = 0,
                     plot.title = element_text(family = "sans", size = 20, face = "plain", color = "black"),
                     axis.title.x = element_text(family="sans", size = 14, color = "black", face="plain"),
                     axis.title.y = element_text(family="sans", size = 14, color = "black", face = "plain", vjust = 
                     ) +
    labs(title = "A",
         x = "Awarding bodies combination",
         y = "Award count")
```

Figure 2B - target awardee career stages:

```
#table(BPdata$Career_stage) #mai options: student, early-career, mid-career, any career stage, unclear

#create new variable with separate career stage values as a list
BPdata2 %>%
  mutate(
    Career_stage_list = str_split(BPdata$Career_stage, pattern = ", ")
  ) -> BPdata2

#BPdata$Career_stage_list <- str_split(BPdata$Career_stage, pattern = ", ") #sam as above but modifies original dat

#make upset plot using library(ggupset)
figure2B <- BPdata2 %>%
  ggplot(aes(x = Career_stage_list)) +
    geom_bar(fill = "#21281D",width = 0.8) +
    scale_y_continuous(limits = c(0, 200)) +
    scale_x_upset(order_by = c("freq")) +
    theme_combmatrix(combmatrix.panel.striped_background = FALSE,
                     combmatrix.panel.point.color.fill = "#21281D",
                     combmatrix.panel.line.size = 0,
                     plot.title = element_text(family = "sans", size = 20, face = "plain", color = "#21281D"),
                     axis.title.x = element_text(family="sans", size = 14, color = "#21281D", face="plain"),
                     axis.title.y = element_text(family="sans", size = 14, color = "#21281D", face = "plain", vjust 
                     ) +
    labs(title = "B",
         x = "Target career stages combination",
         y = "")
```

Figure 2 - combine 2 panels and save:

```
#assemble the panels using patchwork package
figure2 <- figure2A / figure2B +
  plot_layout(ncol = 2, nrow = 1, widths = c(1, 1)) #+  plot_annotation(tag_levels = "A") #does not work with this 

#ggsave(plot = figure2, here("plots", "Fig2AB_award_descriptions_v1.png"), width = 18, height = 8, units = "cm", bg
#ggsave(plot = figure2, here("plots", "Fig2AB_award_descriptions_v1.pdf"), width = 18, height = 8, units = "cm", sc
```

Summary of included awards by awarding or funding body type:
#names(BPdata) - Total number of awards associated with a journal or a few related journals: 176 (79%).
- Total number of awards associated with a learned society: 144 (65%).
- Total number of awards associated with other organisation (e.g. a publisher, university, charity): 39 (18%).
- Most commonly mentioned other organisations: 15 mention "Elsevier", 6 mention "Wiley".

Summary by target career stage:

- Total number of awards for 'any career stage' authors: 179 (81).
- Total number of awards for 'early career' authors: 0 (0).
- Total number of awards for 'student' authors: 27 (12).
- Total number of awards for 'student' and/or 'early career' authors: 20 (9).
- Total number of awards for 'mid-career'authors: 5 (2).

Figure 3 - summary of extracted award characteristics:

```
#BPdata %>%
#    count(Open_science) %>%
#    ggplot(aes(x = Open_science, y = n, fill = Open_science)) +
#    geom_col(width = 0.7) +
#    geom_text(position = position_stack(vjust = 0.5), aes(label = n)) +
#    theme_classic() +
#    coord_flip() +
#    labs(x = "Open Science valued", y = "Article count", fill = "Open_science") +
#    theme(legend.position = "none", axis.title.x = element_text(size = 10))
```

Figure 3A - Summary plots of key award characteristics:

```
BPdata %>%
  select(Award_individual, Flexible_eligibility, Inclusivity_statement, Assessors_transparency, Award_integrity_men

## reshape into long dataframe format:
BPdata3A %>% gather(key = var_name, value = value, 1:ncol(BPdata3A)) -> BPdata3Along

#reorder var_name levels
# BPdata3Along <- BPdata3Along %>% mutate(var_name = factor(var_name, levels = rev(c("Award_individual",
#                                                                              "Flexible_eligibility",
#                                                                              "Inclusivity_statement",
#                                                                              "Assessors_transparency",
#                                                                              "Award_integrity_mentioned",
#                                                                              "Process_transparency",
#                                                                              "Self_nomination",
#                                                                              "Letter_required",
#                                                                              "Feedback_availability",
#                                                                              "Award_contact_provided"
#                                                                              ))))


#reorder value levels:
# BPdata3Along <- BPdata3Along %>% mutate(value = factor(value, levels = rev(c("no",
#                                                                              "yes",
#                                                                              "not applicable",
#                                                                              "not available"))))

# colSums(BPdata3A == "no") #check how many "no" values each column has

BPdata3Along <- BPdata3Along %>% mutate(var_name = factor(var_name, levels = rev(c("Flexible_eligibility",
                                                                           "Assessors_transparency",
                                                                           "Award_individual",
                                                                           "Letter_required",
                                                                           "Self_nomination",
                                                                           "Award_contact_provided",
                                                                           "Award_integrity_mentioned",
                                                                           "Inclusivity_statement",
                                                                           "Process_transparency",
                                                                           "Feedback_availability"
                                                                           ))))

BPdata3Along <- BPdata3Along %>% mutate(value = factor(value, levels = rev(c("yes",
                                                                           "not available",
                                                                           "no"))))
#wacolors$palouse #palette to be used

figure3A <- BPdata3Along %>%
  group_by(var_name) %>%
  count(var_name, value) %>%
  ggplot(aes(x = var_name,
             y = n,
             fill = value,
             pattern = value,
             pattern_angle = value)) +
  geom_bar_pattern(stat = "identity",
    pattern_density = 0.1,
    pattern_spacing = 0.024,
    pattern_key_scale_factor = 0.8,
    pattern_alpha = 0.5,
    pattern_fill    = 'white',
    pattern_colour  = 'white') +
  coord_flip() +
  scale_fill_manual(values = c("#8A6172", "#C0A43D",  "#CCBA98", "#748A52")) +
  scale_x_discrete(labels = rev(c("Eligibility career timeline is flexible",
                              "Assessors are revealed",
                              "Award is focused on individual authors",
                              "Nomination letter is required",
                              "Self-nomination is allowed",
                              "Award contact point is provided",
                              "Award integrity mentioned",
                              "Inclusivity statement or encouragement is provided",
                              "Process is transparent",
                              "Feedback is availabile"
                              ))) +
  labs(x = "Award characteristics",
       y = "Award count") +
  theme_classic()
```

Figure 3B - Summary plots of key award characteristics:

```
BPdata %>% select(Criteria_transparency,
                  Award_impact_metrics_mentioned,
                  Award_impact_metrics_only,
                  Open_science) -> BPdata3B

## reshape into long dataframe format:
BPdata3B %>%
  gather(key = var_name, value = value, 1:ncol(BPdata3B)) -> BPdata3Blong
#str(BPdata3long)

#reorder var_name levels
BPdata3Blong <- BPdata3Blong %>%
  mutate(var_name = factor(var_name, levels = rev(c( "Criteria_transparency",
                                                     "Award_impact_metrics_mentioned",
                                                     "Award_impact_metrics_only",
                                                     "Open_science"))))

#reorder value levels:
BPdata3Blong <- BPdata3Blong %>%
  mutate(value = factor(value, levels = rev(c("yes",
                                              "not available",
                                              "no"))))
figure3B <- BPdata3Blong %>%
  group_by(var_name) %>%
  count(var_name, value) %>%
  #ggplot(aes(x = var_name, y = n, fill = value)) +
  ggplot(aes(x = var_name,
             y = n,
             fill = value,
             pattern = value,
             pattern_angle = value)) +
  geom_bar_pattern(stat = "identity",
    pattern_density = 0.1,
    pattern_key_scale_factor = 0.4,
    pattern_alpha = 0.5,
    pattern_fill    = 'white',
    pattern_colour  = 'white') +
  coord_flip() +
  scale_fill_manual(values = c("#8A6172", "#CCBA98", "#748A52" )) +
  #geom_bar(aes(fill = value), stat = "identity") +
  #coord_flip() +
  labs(x = "Award criteria",
     y = "Award count") +
  scale_x_discrete(labels = rev(c("Assessment criteria are detailed out",
                              "Impact metrics are mentioned in criteria",
                              "Impact metrics are only criteria",
                              "Open Science practices are valued"))) +
  theme_classic()
```

Figure 3 - combine 2 panels and save:

```
#assemble the panels using patchwork package
figure3 <- figure3A / figure3B +
  plot_layout(ncol = 1, nrow = 2, heights = c(2, 1)) +
  plot_annotation(tag_levels = "A")

#ggsave(plot = figure3, here("plots", "Fig3AB_award_descriptions_v1.png"), width = 18, height = 10, units = "cm", b
#ggsave(plot = figure3, here("plots", "Fig3AB_award_descriptions_v1.pdf"), width = 18, height = 10, units = "cm", s
```

Summary by focus on individuals ("individual award") or whole article ("team award"):

- Total number of awards for individual authors: 66 (30%).
- Total number of awards for individual authors with any career stage being eligible: 28 (42%).
- Total number of awards for individual authors with inflexible time limits for eligibility: 21 (32% of all individual awards).
- Total number of awards for individual authors with flexible time limits for eligibility: 5 (8% of all individual awards). Note: remaining are "not applicable", usually because there were no time-related limits for eligibility, e.g. due to any career stage being eligible.

Summary by basic award characteristics:

- Total number of awards with an inclusivity statement: 2 (1%).
- Total number of awards with assessor transparency: 114 (54%).
- Total number of awards with assessment integrity mentioned: 19 (9%).
- Total number of awards with process transparency: 2 (1%).
- Total number of awards with self-nomination allowed: 28 (13%).
- Total number of awards with required nomination letter:38 (18%).
- Total number of awards with feedback available: 0 (0%).
- Total number of awards with contact details available: 38 (18%).

Summary by award criteria characteristics:

- Total number of awards with non-vague criteria: 40 (19%).
- Total number of awards mentioning impact metrics: 21 (10%).
- Total number of awards based only on impact metrics: 8 (4%).
- Total number of awards which value Open Science: 1 (0%).
- Total number of awards with cash award: 112 (53%).

### Award characteristics by Subject Area.

Plot Award_individual by Subject Area:

```
#table(BPdata$Award_individual) #as a table
#table(BPdata$Scimago_Subject_Area, BPdata$Award_individual) #as a table - 9 disciplines have 0!

BPdata %>%
    mutate(Award_individual = factor(Award_individual,
                                     levels = c("yes",
                                                "no"))) %>% #reorder value levels
    count(Scimago_Subject_Area, Award_individual) %>%
    ggplot(aes(x = reorder(Scimago_Subject_Area, desc(Scimago_Subject_Area)),
               y = n,
               fill = Award_individual)) +
    geom_col(width = 0.8, position = position_stack(reverse = TRUE)) +
    coord_flip() +
    scale_y_continuous(breaks = c(0, 5, 10)) +
    theme_classic() +
    scale_fill_manual(values = c("#748A52", "#8A6172")) +
    labs(x = "Scimago Subject Area",
         y = "Award count",
         fill = "Award is focused on individual authors:") +
    theme(legend.position = "top",
          axis.title.x = element_text(size = 10))
```

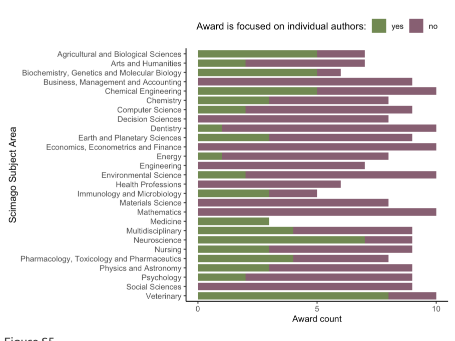


Figure S5.

*Individual-focused awards - by Subject Area.*

Plot Flexible_eligibility by SA:

```
#table(BPdata$Flexible_eligibility, useNA = "always") #as a table
#table(BPdata$Scimago_Subject_Area, BPdata$Flexible_eligibility, useNA = "always") #as a table

BPdata %>%
    mutate(Flexible_eligibility = factor(Flexible_eligibility,
                                         levels = c("yes",
                                                    "not available",
                                                    "not applicable",
                                                    "no"))) %>%
    count(Scimago_Subject_Area, Flexible_eligibility) %>%
    ggplot(aes(x = reorder(Scimago_Subject_Area, desc(Scimago_Subject_Area)),
               y = n,
               fill = Flexible_eligibility)) +
    geom_col(width = 0.8, position = position_stack(reverse = TRUE)) +
    coord_flip() +
    scale_y_continuous(breaks = c(0, 5, 10)) +
    theme_classic() +
    scale_fill_manual(values = c("#748A52", "#CCBA98", "#C0A43D", "#8A6172")) +
    #scale_fill_manual(values = c("#FA8072", "#FA807210", "#F1FFC1", "#C1FFC1")) +
    labs(x = "Scimago Subject Area",
         y = "Award count",
         fill = "Eligibility career timeline is flexible:") +
    theme(legend.position = "top",
          axis.title.x = element_text(size = 10))
```

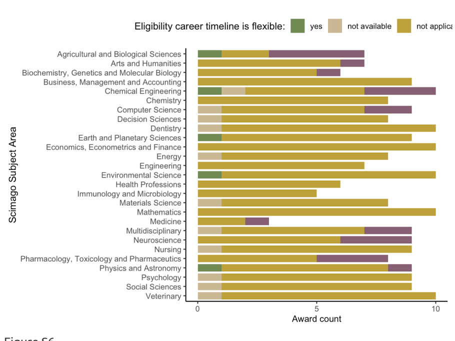


Figure S6.

*Award eligibility timeline flexibility - by Subject Area.*

Plot Inclusivity_statement by SA:

```
#table(BPdata$Scimago_Subject_Area, BPdata$Inclusivity_statement) #as a table

BPdata %>%
    mutate(Inclusivity_statement = factor(Inclusivity_statement,
                                          levels = (c("yes",
                                                      "not available",
                                                      "no")))) %>% #reorder value levels
    count(Scimago_Subject_Area, Inclusivity_statement) %>%
    ggplot(aes(x = reorder(Scimago_Subject_Area, desc(Scimago_Subject_Area)),
               y = n,
               fill = Inclusivity_statement)) +
    geom_col(width = 0.8, position = position_stack(reverse = TRUE)) +
    coord_flip() +
    scale_y_continuous(breaks = c(0, 5, 10)) +
    theme_classic() +
    scale_fill_manual(values = c("#748A52", "#CCBA98", "#8A6172")) +
    labs(x = "Scimago Subject Area",
         y = "Award count",
         fill = "Inclusivity statement or encouragement is provided:") +
    theme(legend.position = "top",
          axis.title.x = element_text(size = 10))
```

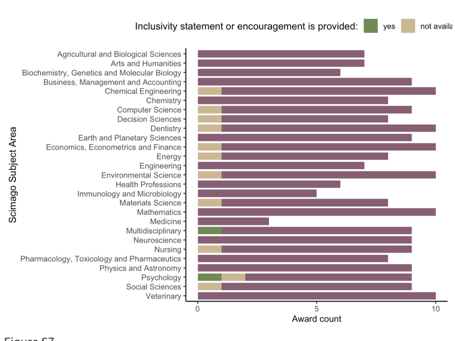


Figure S7.

*Award inclusivity statement or encouragement - by Subject Area.*

Plot Assessors_transparency by SA:

```
#table(BPdata$Scimago_Subject_Area, BPdata$Assessors_transparency) #as a table

BPdata %>%
    mutate(Assessors_transparency = factor(Assessors_transparency, levels = (c("yes",
                                                                              "not available",
                                                                              "no")))) %>% #reorder value levels
    count(Scimago_Subject_Area, Assessors_transparency) %>%
    ggplot(aes(x = reorder(Scimago_Subject_Area, desc(Scimago_Subject_Area)),
               y = n,
               fill = Assessors_transparency)) +
    geom_col(width = 0.8, position = position_stack(reverse = TRUE)) +
    coord_flip() +
    scale_y_continuous(breaks = c(0, 5, 10)) +
    theme_classic() +
    scale_fill_manual(values = c("#748A52", "#CCBA98", "#8A6172")) +
    labs(x = "Scimago Subject Area",
         y = "Award count",
         fill = "Assessors are revealed (names or as journal editors):") +
    theme(legend.position = "top",
          axis.title.x = element_text(size = 10))
```

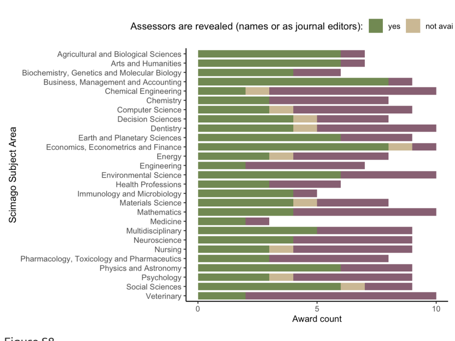


Figure S8.

*Award assessors transparency - by Subject Area.*

Plot Process_transparency by SA:

```
#table(BPdata$Scimago_Subject_Area, BPdata$Process_transparency) #as a table

BPdata %>%
    mutate(Process_transparency = factor(Process_transparency, levels = (c("yes",
                                                                          "not available",
                                                                          "no")))) %>% #reorder value levels
    count(Scimago_Subject_Area, Process_transparency) %>%
    ggplot(aes(x = reorder(Scimago_Subject_Area, desc(Scimago_Subject_Area)),
               y = n,
               fill = Process_transparency)) +
    geom_col(width = 0.8, position = position_stack(reverse = TRUE)) +
    coord_flip() +
    scale_y_continuous(breaks = c(0, 5, 10)) +
    theme_classic() +
    scale_fill_manual(values = c("#748A52", "#CCBA98", "#8A6172")) +
    labs(x = "Scimago Subject Area",
         y = "Award count",
         fill = "Process is transparent:") +
    theme(legend.position = "top",
          axis.title.x = element_text(size = 10))
```

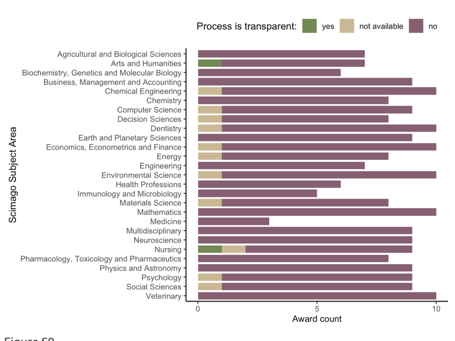


Figure S9.

*Award process transparency - by Subject Area.*

Plot Self_nomination by SA:

```
#table(BPdata$Scimago_Subject_Area, BPdata$Self_nomination) #as a table

BPdata %>%
    mutate(Self_nomination = factor(Self_nomination, levels = (c("yes",
                                                                "not available",
                                                                "no")))) %>% #reorder value levels
    count(Scimago_Subject_Area, Self_nomination) %>%
    ggplot(aes(x = reorder(Scimago_Subject_Area, desc(Scimago_Subject_Area)),
               y = n,
               fill = Self_nomination)) +
    geom_col(width = 0.8, position = position_stack(reverse = TRUE)) +
    coord_flip() +
    scale_y_continuous(breaks = c(0, 5, 10)) +
    theme_classic() +
    scale_fill_manual(values = c("#748A52", "#CCBA98", "#8A6172")) +
    labs(x = "Scimago Subject Area",
         y = "Award count",
         fill = "Self-nomination is allowed:") +
    theme(legend.position = "top",
          axis.title.x = element_text(size = 10))
```

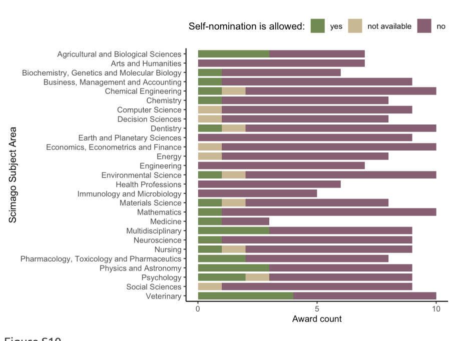


Figure S10.

*Award self-nomination explicitly allowed - by Subject Area.*

Plot Letter_required by SA:

```
#table(BPdata$Scimago_Subject_Area, BPdata$Letter_required) #as a table

BPdata %>%
    mutate(Letter_required = factor(Letter_required, levels = (c("yes",
                                                                "not available",
                                                                "no")))) %>% #reorder value levels
    count(Scimago_Subject_Area, Letter_required) %>%
    ggplot(aes(x = reorder(Scimago_Subject_Area, desc(Scimago_Subject_Area)),
               y = n,
               fill = Letter_required)) +
    geom_col(width = 0.8, position = position_stack(reverse = TRUE)) +
    coord_flip() +
    scale_y_continuous(breaks = c(0, 5, 10)) +
    theme_classic() +
    scale_fill_manual(values = c("#748A52", "#CCBA98", "#8A6172")) +
    labs(x = "Scimago Subject Area",
         y = "Award count",
         fill = "Nomination letter is required:") +
    theme(legend.position = "top",
          axis.title.x = element_text(size = 10))
```

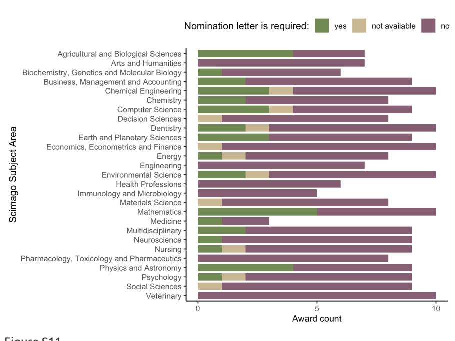


Figure S11.

*Award nomination letter required - by Subject Area.*

Plot Feedback_availability by SA:

```
#table(BPdata$Scimago_Subject_Area, BPdata$Feedback_availability) #as a table

BPdata %>%
    mutate(Feedback_availability = factor(Feedback_availability, levels = (c("yes",
                                                                            "not available",
                                                                            "no")))) %>% #reorder value levels
    count(Scimago_Subject_Area, Feedback_availability) %>%
    ggplot(aes(x = reorder(Scimago_Subject_Area, desc(Scimago_Subject_Area)),
               y = n,
               fill = Feedback_availability)) +
    geom_col(width = 0.8, position = position_stack(reverse = TRUE)) +
    coord_flip() +
    scale_y_continuous(breaks = c(0, 5, 10)) +
    theme_classic() +
    scale_fill_manual(values = c("#CCBA98", "#8A6172")) +
    labs(x = "Scimago Subject Area",
         y = "Award count",
         fill = "Feedback is availabile:") +
    theme(legend.position = "top",
          axis.title.x = element_text(size = 10))
```

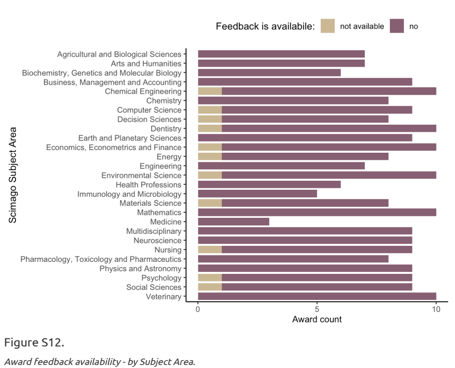


Figure S12.

*Award feedback availability - by Subject Area.*

Plot Award_contact_provided by SA:

```
#table(BPdata$Scimago_Subject_Area, BPdata$Award_contact_provided) #as a table

BPdata %>%
    mutate(Award_contact_provided = factor(Award_contact_provided, levels = (c("yes",
                                                                              "not available",
                                                                              "no")))) %>% #reorder value levels
    count(Scimago_Subject_Area, Award_contact_provided) %>%
    ggplot(aes(x = reorder(Scimago_Subject_Area, desc(Scimago_Subject_Area)),
               y = n,
               fill = Award_contact_provided)) +
    geom_col(width = 0.8, position = position_stack(reverse = TRUE)) +
    coord_flip() +
    scale_y_continuous(breaks = c(0, 5, 10)) +
    theme_classic() +
    scale_fill_manual(values = c("#748A52", "#CCBA98", "#8A6172")) +
    labs(x = "Scimago Subject Area",
         y = "Award count",
         fill = "Award contact point is provided:") +
    theme(legend.position = "top",
          axis.title.x = element_text(size = 10))
```

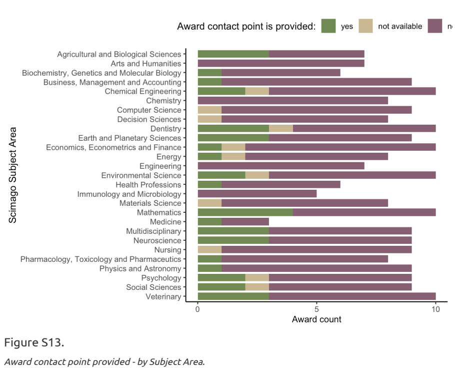


Figure S13.

*Award contact point provided - by Subject Area.*

Plot Criteria_transparency by SA:

```
#table(BPdata$Scimago_Subject_Area, BPdata$Criteria_transparency) #as a table

BPdata %>%
    mutate(Criteria_transparency = factor(Criteria_transparency, levels = (c("yes",
                                                                            "not available",
                                                                            "no")))) %>% #reorder value levels
    count(Scimago_Subject_Area, Criteria_transparency) %>%
    ggplot(aes(x = reorder(Scimago_Subject_Area, desc(Scimago_Subject_Area)),
               y = n,
               fill = Criteria_transparency)) +
    geom_col(width = 0.8, position = position_stack(reverse = TRUE)) +
    coord_flip() +
    scale_y_continuous(breaks = c(0, 5, 10)) +
    theme_classic() +
    scale_fill_manual(values = c("#748A52", "#CCBA98", "#8A6172")) +
    labs(x = "Scimago Subject Area",
         y = "Award count",
         fill = "Assessment criteria are detailed out:") +
    theme(legend.position = "top",
          axis.title.x = element_text(size = 10))
```

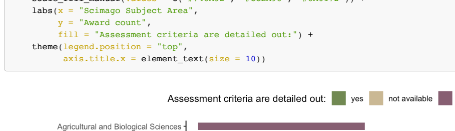

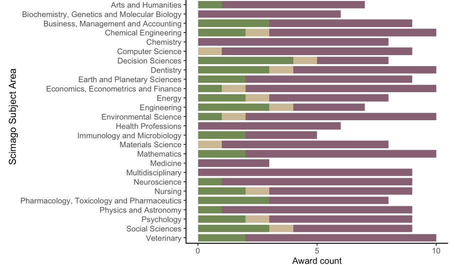


Figure S14.

*Detailed award assessment criteria - by Subject Area.*

Plot Award_impact_metrics_mentioned by SA:

```
#table(BPdata$Scimago_Subject_Area, BPdata$Award_impact_metrics_mentioned) #as a table

BPdata %>%
    mutate(Award_impact_metrics_mentioned = factor(Award_impact_metrics_mentioned, levels = (c("yes",
                                                                                               "not available",
                                                                                               "no")))) %>% #reorde
    count(Scimago_Subject_Area, Award_impact_metrics_mentioned) %>%
    ggplot(aes(x = reorder(Scimago_Subject_Area, desc(Scimago_Subject_Area)),
               y = n,
               fill = Award_impact_metrics_mentioned)) +
    geom_col(width = 0.8, position = position_stack(reverse = TRUE)) +
    coord_flip() +
    scale_y_continuous(breaks = c(0, 5, 10)) +
    theme_classic() +
    scale_fill_manual(values = c("#748A52", "#CCBA98", "#8A6172")) +
    labs(x = "Scimago Subject Area",
         y = "Award count",
         fill = "Impact metrics are mentioned in criteria:") +
    theme(legend.position = "top",
          axis.title.x = element_text(size = 10))
```

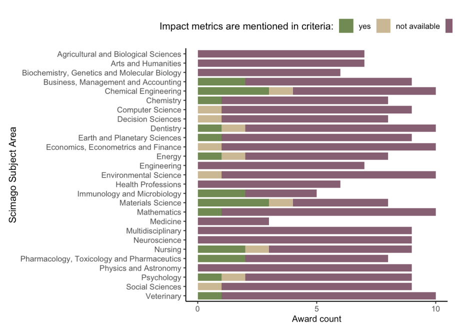


Figure S15.

*Award assessment criteria with impact metrics - by Subject Area.*

Plot Award_impact_metrics_only by SA:

```
#table(BPdata$Scimago_Subject_Area, BPdata$Award_impact_metrics_only) #as a table

BPdata %>%
    mutate(Award_impact_metrics_only = factor(Award_impact_metrics_only, levels = (c("yes",
                                                                                     "not available",
                                                                                     "no")))) %>% #reorder value le
    count(Scimago_Subject_Area, Award_impact_metrics_only) %>%
    ggplot(aes(x = reorder(Scimago_Subject_Area, desc(Scimago_Subject_Area)),
               y = n,
               fill = Award_impact_metrics_only)) +
    geom_col(width = 0.8, position = position_stack(reverse = TRUE)) +
    coord_flip() +
    scale_y_continuous(breaks = c(0, 5, 10)) +
    theme_classic() +
    scale_fill_manual(values = c("#748A52", "#CCBA98", "#8A6172")) +
    labs(x = "Scimago Subject Area",
         y = "Award count",
         fill = "Impact metrics are only criteria:") +
    theme(legend.position = "top",
          axis.title.x = element_text(size = 10))
```

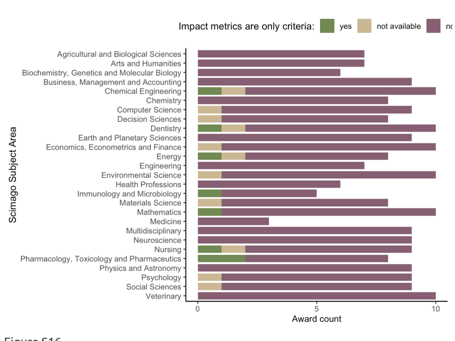


Figure S16.

*Impact metrics as only criteria - award scores by Subject Area.*

Plot Open_science by SA:

```
#table(BPdata$Scimago_Subject_Area, BPdata$Open_science) #as a table

BPdata %>%
    mutate(Open_science = factor(Open_science, levels = (c("yes",
                                                           "not available",
                                                           "no")))) %>% #reorder value levels
    count(Scimago_Subject_Area, Open_science) %>%
    ggplot(aes(x = reorder(Scimago_Subject_Area, desc(Scimago_Subject_Area)),
               y = n,
               fill = Open_science)) +
    geom_col(width = 0.8, position = position_stack(reverse = TRUE)) +
    coord_flip() +
    scale_y_continuous(breaks = c(0, 5, 10)) +
    theme_classic() +
    scale_fill_manual(values = c("#748A52", "#CCBA98", "#8A6172")) +
    labs(x = "Scimago Subject Area",
         y = "Award count",
         fill = "Open Science practices are valued:") +
    theme(legend.position = "top",
          axis.title.x = element_text(size = 10))
```

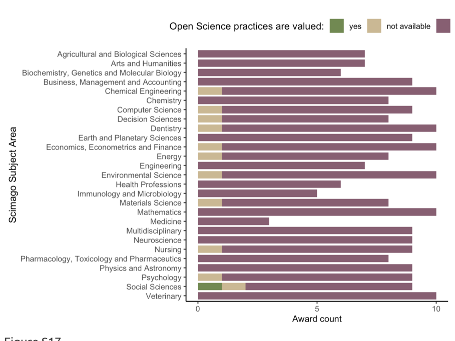


Figure S17.

*Open Science practices - award scores by Subject Area.*

### Text mining of award descriptions.

Award descriptions - frequencies of words:

```
#Using packages tidytext and stopwords
Award_description_txt <- tibble(txt = tolower(BPdata$Award_description))
Award_description_txt <- Award_description_txt %>% unnest_tokens(output = word, input = txt, token = "words", to_lo
Award_description_txt <- Award_description_txt %>% anti_join(get_stopwords()) #remove stop words from library(stopw
```

```
## Joining with `by = join_by(word)`
```

```
## Joining with `by = join_by(word)`
Award_description_txt <- Award_description_txt %>% anti_join(get_stopwords()) #remove stop words from library(stopw
```

```
## Joining with `by = join_by(word)`
```

```
## Joining with `by = join_by(word)`
Award_description_txt$word <- tokenize_word_stems(Award_description_txt$word) #make all word stems lowercase
word.freq <- Award_description_txt %>% count(word, sort = TRUE) #count words
```

Award descriptions - count mentions of specific words (stemmed):

```
#create list of specific words (stemmed) to count within strings
specific.words <- c("best", "outstand", "impact", "original", "significant", "merit", "excell", "innovat", "novel",

#prepare award descriptions as a single lowercase string
descriptions <- BPdata %>%
  filter(!is.na(Award_description)) %>%
  select(Award_description) %>%
  tolower() #single lowercase string

#sum of all mentions for each word
specific.words.mentions <- specific.words %>%
  map_int(~ str_count(tolower(descriptions), .x))

#prepare award descriptions while keeping them separate for each award
descriptions2 <- tolower(BPdata$Award_description) #vector of lowercase strings

#sum of mentions per award for each word (counts only one mention per award)
specific.words.mentions2 <- specific.words %>%
  map_int(~ sum(str_detect(descriptions2, .x), na.rm = TRUE))

# ## doing the same as above, but manually:
# #count all mentions of words(parts) individually, e.g.:
# sum(str_count(tolower(BPdata$Award_description), "best"), na.rm = TRUE) #169
# #counting once per award, e.g.
# sum(str_detect(BPdata$Award_description, "best"), na.rm = TRUE) #82
```

Figure 4A - frequencies of specific words - all mentions in award descriptions.

```
words.df <- tibble(Words = specific.words,
                   Count_all = specific.words.mentions,
                   Count_once = specific.words.mentions2)

#wacolors$palouse #palette to be used

figure4A <- words.df %>%
    ggplot(aes(x = reorder(Words, Count_all),
               y = Count_all)) +
    geom_col(width = 0.8,
             fill = "#2D3F4A") +
    coord_flip() +
    scale_y_continuous(breaks = c(0, 50, 100, 150),
                       limits = c(0, 200)) +
    theme_bw() +
    labs(x = "Word stem", y = "Count of all mentions") +
    annotate(geom = "rect",xmin = 8.5, xmax = Inf, ymin = -Inf, ymax = Inf, fill = "#8A6172", alpha = 0.2) +
    annotate(geom = "rect",xmin = -Inf, xmax = 8.5, ymin = -Inf, ymax = Inf, fill = "#748A52", alpha = 0.2) +
    theme(legend.position = "none",
          axis.title.x = element_text(size = 10))
```

Figure 4B - frequencies of specific words - first mentions in award descriptions.

```
figure4B <- words.df %>%
    ggplot(aes(x = reorder(Words, Count_all),
               y = Count_once)) +
    geom_col(width = 0.8,
             fill = "#2D3F4A",
             alpha = 0.7) +
    coord_flip() +
    scale_y_continuous(breaks = c(0, 50, 100, 150),
                       limits = c(0, 200)) +
    theme_bw() +
    labs(x = "Word stem",
         y = "Count of first mention per award") +
    scale_x_discrete(labels = NULL) +
    labs(x = "")  + #used to remove vertical labels, also breaks = NULL
    annotate(geom = "rect",xmin = 8.5, xmax = Inf, ymin = -Inf, ymax = Inf, fill = "#8A6172", alpha = 0.2) +
    annotate(geom = "rect",xmin = -Inf, xmax = 8.5, ymin = -Inf, ymax = Inf, fill = "#748A52", alpha = 0.2) +
    theme(legend.position = "none",
          axis.title.x = element_text(size = 10))
```

Figure 4 - combine 2 panels and save.

```
#assemble the panels using patchwork package
figure4 <- figure4A / figure4B +
  plot_layout(ncol = 2, nrow = 1) +
  plot_annotation(tag_levels = "A")
#ggsave(plot = figure4, here("plots", "Fig4AB_words_counts_v1.png"), width = 18, height = 8, units = "cm", dpi = "r
#ggsave(plot = figure4, here("plots", "Fig4AB_words_counts_v1.pdf"), width = 18, height = 8, units = "cm", scale =
```

### Characteristics of past winners.

```
#length(unique(BPdata$Award_name[BPdata$N_winners_extracted != 0])) #61 - number from the records in the award-leve
#length(unique(BPindiv$award_name)) #61 - number from records in past winner's data (matching)
```

Total number of extracted awards: 61.
Number of Subject Areas with 0 extracted award winners: 8.

Plot numbers of awards extracted per SA:

```
# #from BPindiv:
# BPindiv %>%
#   group_by(award_SA) %>%
#   summarise(count = n_distinct(award_name))

# #from BPindiv:
# BPindiv %>%
#   group_by(award_SA) %>%
#   summarise(count = n_distinct(award_name)) %>% summarise(mean(count)) #3

#from BPdata - with winners data:
BPdata %>%
  group_by(Scimago_Subject_Area) %>%
  filter(N_winners_extracted != 0) %>%
  count(Scimago_Subject_Area) -> awards_perSA

#No data for:
setdiff(unique(BPdata$Scimago_Subject_Area), unique(BPdata$Scimago_Subject_Area[BPdata$N_winners_extracted != 0]))

#combine two tibbles
awards_perSA <- bind_rows(awards_perSA ,tibble(Scimago_Subject_Area = awards_perSA_none, n = rep(0, length(awards_p


# #plot counts where >0 awards per SA
# p_count.awards.SA <- BPindiv %>%
#   group_by(award_SA) %>%
#   summarise(count = n_distinct(award_name)) %>%
#   ggplot(aes(x = reorder(award_SA, desc(award_SA)),
#              y = count)) +
#   geom_bar(stat = "identity",
#            position = position_dodge(0.9)) +
#   coord_flip() +
#   theme_bw() +
#   labs(x = "Scimago Subject Area",
#        y = "Award count")

#plot counts per SA for all SA
p_count.awards.SA <- awards_perSA %>%
  ggplot(aes(x = reorder(Scimago_Subject_Area, desc(Scimago_Subject_Area)),
             y = n)) +
  geom_bar(stat = "identity",
           position = position_dodge(0.9)) +
  coord_flip() +
  theme_bw() +
  labs(x = "Scimago Subject Area",
       y = "Award count")

p_count.awards.SA
```

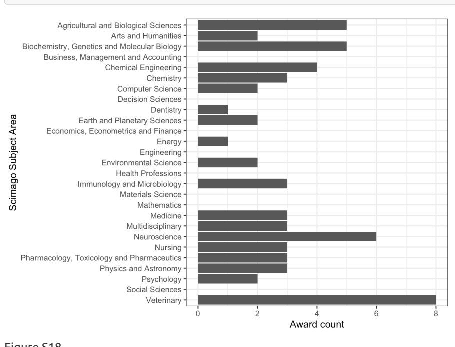


Figure S18.

*Number of awards with extracted individual winners data - counts by Subject Area.*

Total number of extracted records (awardee names): 1079.

```
#sum(BPdata$N_winners_extracted, na.rm = TRUE) #number from the records in the award-level data set
#dim(BPindiv)[1] #number of records in past winner's data (matching)
```

Plot numbers of records (names) extracted per SA:

```
#from BPdata:
BPdata %>%
  group_by(Scimago_Subject_Area) %>%
  select(N_winners_extracted) %>%
  filter(!is.na(N_winners_extracted)) %>%
  summarise(sum_winners = sum(N_winners_extracted), .groups = 'drop') -> winners_perSA1 #use for plotting all SA
```

```
## Adding missing grouping variables: `Scimago_Subject_Area`
```

```
#dim(winners_perSA1[winners_perSA1$sum_winners != 0, ])[1] #number of SA with any extracted names
#winners_perSA1[winners_perSA1$sum_winners != 0, ] #table of SA with any extracted names

##Using BPindiv
##Adding missing grouping variables: Scimago_Subject_Area
#table(BPindiv$award_SA)
#dim(table(BPindiv$award_SA, useNA = "always")) #19 SA with any extracted names
# BPindiv %>%
#   group_by(award_SA) %>% count()
#   -> winners_perSA1_extracted #does not show SA with 0 extracted
#median(winners_perSA1_extracted$n) #42

# #plot counts where >0 winners per SA
# p_count.awardees.SA <-
#   BPindiv %>%
#   group_by(award_SA) %>%
#   count() %>%
#   ggplot(aes(x = reorder(award_SA, desc(award_SA)),
#              y = n)) +
#   geom_bar(stat = "identity",
#            position = position_dodge(0.9)) +
#   coord_flip() +
#   theme_bw() +
#   #scale_x_discrete(labels = NULL) + labs(x = "")  + #used to remove vertical labels, also breaks = NULL
#   labs(x = "Scimago Subject Area",
#        y = "Awardee count")

#plot counts per SA for all SA
p_count.awardees.SA <- winners_perSA1 %>%
  ggplot(aes(x = reorder(Scimago_Subject_Area, desc(Scimago_Subject_Area)),
             y = sum_winners)) +
  geom_bar(stat = "identity",
           position = position_dodge(0.9)) +
  coord_flip() +
  theme_bw() +
  labs(x = "Scimago Subject Area",
       y = "Awardee count")

p_count.awardees.SA
```

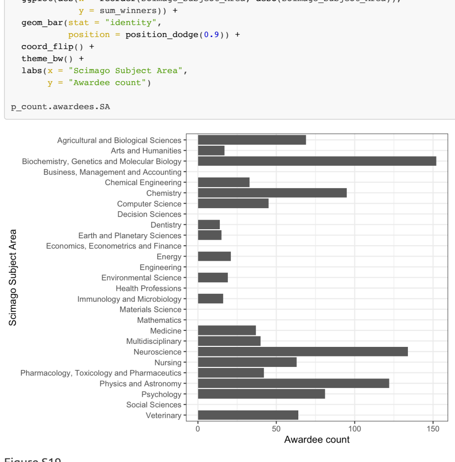


Figure S19.

*Numbers of extracted individual winners - counts by Subject Area.*

Number of Subject Areas without any extracted individual winners: 8.

Plot numbers of awards with individual winner data per year:

```
BPindiv %>%
  group_by(award_year) %>%
  summarise(n = n_distinct(award_name)) %>%
  ggplot(aes(x = award_year,
             y = n)) +
  geom_bar(stat = "identity",
           position = position_dodge(0.9)) +
  theme_bw() +
  labs(x = "Award year", y = "Award count")
```

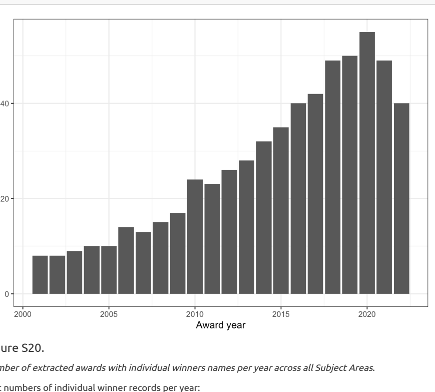


Figure S20.

*Number of extracted awards with individual winners names per year across all Subject Areas.*

Plot numbers of individual winner records per year:

```
#table(BPindiv$award_year, useNA = "always")

BPindiv %>%
  group_by(award_year) %>%
  count() %>%
  ggplot(aes(x = award_year,
             y = n)) +
  geom_bar(stat = "identity",
           position = position_dodge(0.9)) +
  theme_bw() +
  labs(x = "Award year",
       y = "Awardee count")
```

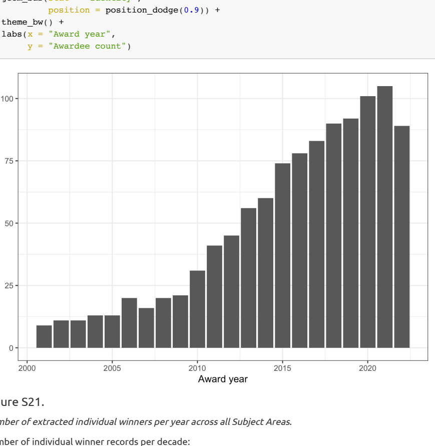


Figure S21.

*Number of extracted individual winners per year across all Subject Areas.*

Number of individual winner records per decade:

```
#create decade variable 2001-2010, 2011-2020, 2021-2022
BPindiv %>%  mutate(Decade = case_when(
    award_year >= 2001 & award_year <= 2010 ~ "2001-2010",
    award_year >= 2011 & award_year <= 2020 ~ "2011-2020",
    award_year >= 2021 & award_year <= 2022 ~ "2021-2022",
    award_year == "Veterinary" ~ "Biology",
    TRUE ~ NA)
    ) -> BPindiv

#table(BPindiv$Decade, useNA = "always")

## 2001-2010 2011-2020 2021-2022      <NA>
##       165       720       194         0
```

### Counts of extracted past awardees by decade:

- 2001-2010: 165.
- 2011-2020: 720.
- 2021-2022: 194.

```
#table(BPindiv$shared, useNA = "always") #156 shared (14% of names), usually shared by 2 authors)
#table(BPindiv$Decade, BPindiv$shared) #more common recently
#table(BPindiv$award_SA, BPindiv$shared) # sharing most frequent in Computer Science
```

Number of individuals sharing awards: 156 (14%).

NOTE: during the first round of data extraction, we located around 3/4 of the winning articles - these were articles where title or DOI were available on the award page or announcement. Often article information was publicly listed only for recently awarded articles and absent for the older ones. Our additional Internet searches increased the number of identified articles.

Number of awards with article information (only reference): 880 (82).
Number of awards with article information (only DOI): 879 (81).
Number of awards with article information (reference or DOI): 879 (81%).

### Awardee gender.

Source of information on gender:
- Pronouns: 187 (17%).
- Photo: 265 (25%).
- Name only:627 (58%).

Total number of past awardees without gender assigned: 0.
Total number of past awardees assigned as a woman: 424 (39%).
Total number of past awardees assigned as a man: 655 (61%).

Plot gender counts overall:

```
#wacolors$san_juan

## set the levels in order we want:
BPindiv$Gender <- factor(BPindiv$awardee_gender)
levels(BPindiv$Gender) <- c("female", "male")

## plot for a all SA and years - not stacked, with count values:
BPindiv %>%
#  group_by(award_SA) %>%
  count(Gender) %>%
  ggplot(aes(x = Gender,
             y = n)) +
  geom_bar(aes(fill = Gender),
           stat = "identity",
           position = position_dodge(0.9)) +
  scale_fill_manual(values = c("#CA884C", "#3A5775")) +
  theme_bw() +
  theme(legend.position = "none",
        legend.box = "horizontal",
        axis.text = element_text(size = 10)) +
  labs(fill = "Awardee gender:") +
  labs(x = "Awardees",
       y = "Awardee count")
```

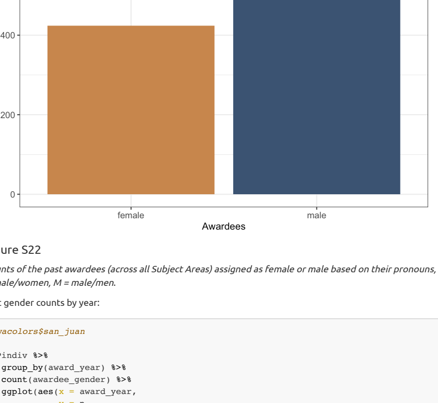


Figure S22

*Counts of the past awardees (across all Subject Areas) assigned as female or male based on their pronouns, photos and names. F = female/women, M = male/men.*

Plot gender counts by year:

```
#wacolors$san_juan

BPindiv %>%
  group_by(award_year) %>%
  count(awardee_gender) %>%
  ggplot(aes(x = award_year,
             y = n,
             fill = awardee_gender)) +
  geom_bar(stat = "identity", position = position_dodge(0.9)) +
  theme_bw() +
  scale_fill_manual(values = c("#CA884C", "#3A5775")) +
  theme(legend.position = "top",
        legend.box = "horizontal",
        axis.text = element_text(size = 10)) +
  labs(fill = "Awardee gender:") +
  labs(x = "Award year",
       y = "Awardee count")
```

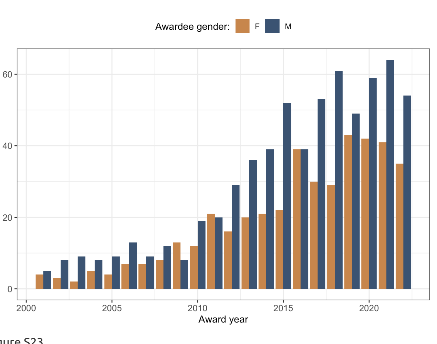


Figure S23.

*Counts of the past awardees (across all Subject Areas) assigned as female or male based on their pronouns, photos and names - by year. F = female/women, M = male/men.*

Plot gender proportion by year:

```
#wacolors$san_juan

BPindiv %>%
  group_by(award_year) %>%
  count(awardee_gender) %>%
  ggplot(aes(x = award_year,
             y = n,
             fill = awardee_gender)) +
  geom_bar(stat = "identity",
           position = "fill") +
  theme_bw() +
  scale_fill_manual(values = c("#CA884C", "#3A5775")) +
  theme(legend.position = "top",
        legend.box = "horizontal",
        axis.text = element_text(size = 10)) +
  labs(fill = "Awardee gender:") +
  labs(x = "Award year",
       y = "Awardee proportion")
```

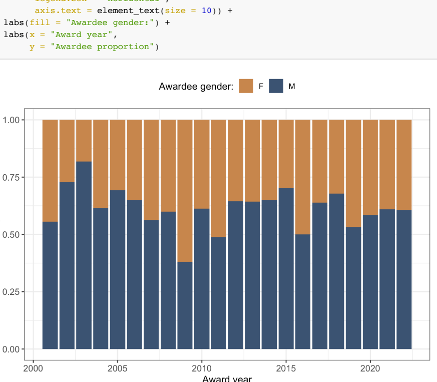


Figure S24.

*Proportions of the past awardees (across all Subject Areas) assigned as female or male based on their pronouns, photos and names - by year. F = female/women, M = male/men.*

Figure 5A - plot of gender counts by decade:

```
##See the actual counts
# BPindiv %>%
#   group_by(Decade) %>%
#   count(awardee_gender) %>%
#   mutate(decade_n = sum(n)) %>%
#   mutate(pct = n / decade_n *100)

## plot for all Subject Areas by Decade - horizontal, not stacked, with count values:
figure5A <- BPindiv %>%
  group_by(Decade) %>%
  count(Gender) %>%
```

```r
  ggplot(aes(y = Decade,
             x = n,
             fill = Gender)) +
  geom_bar(aes(fill = Gender),
           stat = "identity") +
  scale_fill_manual(values = c("#CA884C", "#3A5775")) +
  theme_bw() +
  theme(legend.position="top",
        legend.text = element_text(size = 8),
        legend.title = element_text(size = 8),
        axis.text = element_text(size = 10),
        legend.box = "horizontal",
        legend.margin = margin()) +
  guides(fill = guide_legend(nrow = 1, byrow = TRUE, reverse = TRUE)) +
  labs(fill = "Awardee gender:") +
  labs(y = "Decade",
       x = "Awardee counts")
```

##### Figure 5B - plot of gender proportions by decade:

```r
## plot for a all Subject Areas by Decade - horizontal, stacked, with proportion values:
figure5B <- BPindiv %>%
  group_by(Decade) %>%
  count(Gender) %>%
  ggplot(aes(x = Decade,
             y = n,
             fill = Gender)) +
  geom_bar(aes(fill = Gender),
           stat = "identity",
           position = "fill") + # use , position = "fill" for proportion plot
  coord_flip() +
  scale_fill_manual(values = c("#CA884C", "#3A5775")) +
  theme_bw() +
  theme(legend.position = "none",
        legend.box = "horizontal",
        axis.text = element_text(size = 10)) +
  labs(fill = "Awardee gender:") +
  labs(x = "Decade",
       y = "Awardee proportion")
```

Plot gender counts by Subject Area:

```r
##See the counts by Subject Area
# BPindiv %>%
#    group_by(award_SA) %>%
#    count(awardee_gender) %>%
#    mutate(SA_n = sum(n)) %>%
#    mutate(pct = n / SA_n *100)

## plot for a all awards by SA - not stacked, with count values:
BPindiv %>%
  group_by(award_SA) %>%
  count(Gender) %>%
  ggplot(aes(x = reorder(award_SA, desc(award_SA)),
             y = n,
             fill = Gender)) +
  geom_bar(aes(fill = Gender),
           stat = "identity") + # use , position = "fill" for proportion plot
  coord_flip() +
  scale_fill_manual(values = c("#CA884C", "#3A5775")) +
  theme_bw() +
  theme(legend.position="top",
        legend.box = "horizontal",
        axis.text = element_text(size = 10)) +
  labs(fill = "Awardee gender:") +
  labs(x = "Scimago Subject Area",
       y = "Awardee count")
```

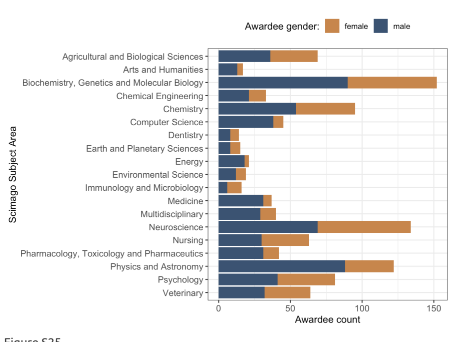


Figure S25.

*Counts of the past awardees assigned as female or male based on their pronouns, photos and names - by Subject Area. F = female/women, M = male/men.*

Plot gender proportions by Subject Area:

```r
## plot for a all awards by SA -stacked, with proportion values:
BPindiv %>%
  group_by(award_SA) %>%
  count(Gender) %>%
  ggplot(aes(x = reorder(award_SA, desc(award_SA)),
             y = n,
             fill = Gender)) +
  geom_bar(aes(fill = Gender),
           stat = "identity",
           position = "fill") +
  coord_flip() +
  scale_fill_manual(values = c("#CA884C", "#3A5775")) +
  theme_bw() +
  theme(legend.position="top",
        legend.box = "horizontal",
        axis.text = element_text(size = 10)) +
  labs(fill = "Awardee gender:") +
  labs(x = "Scimago Subject Area",
       y = "Awardee proportion")
```

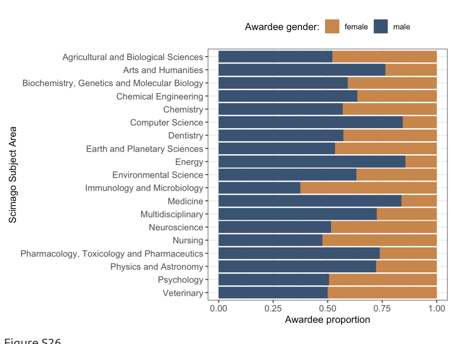


Figure S26.

*Proportions of the past awardees assigned as female or male based on their pronouns, photos and names - by Subject Area. F = female/women, M = male/men.*

Plot awardee gender counts by Subject Area and Decade:

```r
## plot for a all awards by SA and Decade - not stacked, with count values:
BPindiv %>%
  group_by(award_SA, Decade) %>%
  count(Gender) %>%
  ggplot(aes(x = reorder(award_SA, desc(award_SA)),
             y = n,
             fill = Gender)) +
  geom_bar(aes(fill = Gender),
           stat = "identity",
           position = position_dodge(0.8),
           width = 0.8,
           col = "white") + # use , position = "fill" for proportion plot
  facet_wrap(~Decade, scales = "fixed", nrow = 1, ncol = 3) +
  coord_flip() +
  scale_fill_manual(values = c("#CA884C", "#3A5775")) +
  theme_bw() +
  theme(legend.position="top",
        legend.box = "horizontal",
        axis.text = element_text(size = 10)) +
  labs(fill = "Awardee gender:") +
  labs(x = "Scimago Subject Area",
       y = "Awardee count")
```

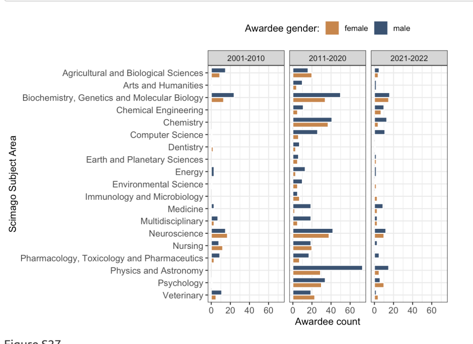


Figure S27.

*Counts of the past awardees assigned as female or male based on their pronouns, photos and names - by Subject Area and Decade. F = female/women, M = male/men.*

#### Affiliation country.

Number of past awardees without affiliation country info: 54 (5%).

table(BPindiv$affiliation_info_source)

34 (3%).

Source of affiliation information:
- Award page: 355 (33%).
- Award announcement: 159 (15%).
- Winning article: 482 (45%).
- Other: 30 (3%).

```r
#count individuals with affiliations:
BPindiv %>%
  filter(!is.na(affiliation_country)) %>%
  nrow() -> affiliation_indiv_count

#count number of unique countries overall:
BPindiv %>%
  filter(!is.na(affiliation_country)) %>%
  summarise(count = n_distinct(affiliation_country)) -> affiliation_country_count

# #count number of unique countries per decade:
# BPindiv %>%
#   filter(!is.na(affiliation_country)) %>%
#   group_by(Decade) %>%
#   summarise(count = n_distinct(affiliation_country))

# #count number of unique countries per year:
# BPindiv %>%
#   filter(!is.na(affiliation_country)) %>%
#   group_by(award_year) %>%
#   summarise(count = n_distinct(affiliation_country))

#count by countries:
BPindiv %>%
  filter(!is.na(affiliation_country)) %>%
  count(affiliation_country) %>%
  arrange(desc(n)) -> country_count

#count for USA only:
BPindiv %>%
  filter(!is.na(affiliation_country)) %>%
  filter(affiliation_country == "USA") %>%
  nrow() -> affiliation_country_USA

# #count countries - view top 20:
# BPindiv %>%
#   filter(!is.na(affiliation_country)) %>%
#   count(affiliation_country) %>%
#   arrange(desc(n)) %>%
#   filter(n>5) %>%
#   View()

#recode top 10 countries + other
BPindiv$Country10 <- recode(BPindiv$affiliation_country,
                            "USA" = "USA",
                            "UK" = "UK",
                            "Australia" = "Australia",
                            "China" = "China",
                            "Germany" = "Germany",
                            "Canada" = "Canada",
                            "Japan" = "Japan",
                            "Netherlands" = "Netherlands",
                            "Italy" = "Italy",
                            "France" = "France",
                            .default = "other")
#table(BPindiv$Country10)

#recode countries with n>5 (20)
#BPindiv$Country20 <- recode(BPindiv$affiliation_country,
                            # "USA" = "USA",
                            # "UK" = "UK",
                            # "Australia" = "Australia",
                            # "China" = "China",
                            # "Germany" = "Germany",
                            # "Canada" = "Canada",
                            # "Japan" = "Japan",
                            # "Netherlands" = "Netherlands",
                            # "Italy" = "Italy",
                            # "France" = "France",
                            # "Austria" = "Austria",
                            # "Spain" = "Spain",
                            # "Sweden" = "Sweden",
                            # "Switzerland" = "Switzerland",
                            # "India" = "India",
                            # "South Korea" = "South Korea",
                            # "Brazil" = "Brazil",
                            # "Singapore" = "Singapore",
                            # "Taiwan" = "Taiwan",
                            # "Belgium" = "Belgium",
                            # .default = "other")
#table(BPindiv$Country20)

## make a simple table with percentages of the total number of Documents
# BPindiv %>%
#   filter(!is.na(affiliation_country)) %>%
#   count(Country10)  %>%
#   arrange(desc(n)) %>%
#   mutate(Country_pct = n/sum(n) * 100)
```

Affiliation country composition summary:

- Out of 1079 extracted individual winners, 1025 (95%) had information on their affiliation country.
- Affiliations represented 42 countries (only first affiliation was used).
- Out of 1025 extracted individual winners with affiliation country information, 487 (48 47.5%) had USA as their affiliation country.

Incorporate data on Global South nations (United Nations FCFSSC) among past awardees:

```r
GS_nations <- c("India",
                "China",
                "Indonesia",
                "Pakistan",
                "Nigeria",
                "Brazil",
                "Bangladesh",
                "Ethiopia",
                "Philippines",
                "Egypt",
                "DR Congo",
                "Vietnam",
                "Iran",
                "Thailand",
                "Tanzania",
                "South Africa",
                "Kenya",
                "Myanmar",
                "Colombia",
                "Uganda",
                "Sudan",
                "Argentina",
                "Algeria",
                "Iraq",
                "Afghanistan",
                "Morocco",
                "Saudi Arabia",
                "Angola",
                "Yemen",
                "Peru",
                "Malaysia",
                "Ghana",
                "Mozambique",
                "Nepal",
                "Madagascar",
                "Ivory Coast",
                "Venezuela",
                "Cameroon",
                "Niger",
                "North Korea",
                "Mali",
                "Burkina Faso",
                "Syria",
                "Sri Lanka",
                "Malawi",
                "Zambia",
                "Chile",
                "Chad",
                "Ecuador",
                "Somalia",
                "Guatemala",
                "Senegal",
                "Cambodia",
                "Zimbabwe",
                "Guinea",
                "Rwanda",
                "Benin",
                "Burundi",
                "Tunisia",
                "Bolivia",
                "Haiti",
                "Jordan",
                "Dominican Republic",
                "Cuba",
                "South Sudan",
                "Honduras",
                "Papua New Guinea",
                "Tajikistan",
                "United Arab Emirates",
                "Togo",
                "Sierra Leone",
                "Laos",
                "Nicaragua",
                "Libya",
                "Paraguay",
                "Turkmenistan",
                "El Salvador",
                "Republic Of The Congo",
                "Singapore",
                "Central African Republic",
                "Liberia",
                "Palestine",
                "Lebanon",
                "Costa Rica",
                "Mauritania",
                "Oman",
                "Panama",
                "Kuwait",
                "Eritrea",
                "Mongolia",
                "Uruguay",
                "Bosnia And Herzegovina",
                "Jamaica",
                "Gambia",
                "Qatar",
                "Botswana",
                "Namibia",
                "Gabon",
                "Lesotho",
                "Guinea Bissau",
                "Equatorial Guinea",
                "Trinidad And Tobago",
                "Bahrain",
                "Timor Leste",
                "Mauritius",
                "Eswatini",
                "Djibouti",
                "Fiji",
                "Comoros",
                "Guyana",
                "Bhutan",
                "Solomon Islands",
                "Suriname",
                "Cape Verde",
                "Maldives",
                "Brunei",
                "Bahamas",
                "Belize",
                "Vanuatu",
                "Barbados",
                "Sao Tome And Principe",
                "Samoa",
                "Saint Lucia",
                "Kiribati",
                "Grenada",
                "Micronesia",
                "Tonga",
                "Seychelles",
                "Saint Vincent And The Grenadines",
                "Antigua And Barbuda",
                "Dominica",
                "Saint Kitts And Nevis",
                "Marshall Islands",
                "Nauru"
)

#count Global South countries in the data set:
# BPindiv %>%
#   filter(!is.na(affiliation_country)) %>%
#   filter(affiliation_country  %in% GS_nations) %>%
#   count(affiliation_country) %>%
#   arrange(desc(n)) #%>% View() #table of GS nations and winner counts

BPindiv %>%
  filter(!is.na(affiliation_country)) %>% nrow() -> n_winners_affil #number of winners with affiliation country

BPindiv %>%
  filter(!is.na(affiliation_country)) %>%
  filter(affiliation_country  %in% GS_nations) %>%
  count(affiliation_country) %>%
  arrange(desc(n)) %>% count() -> n_winners_GS_countries #number of countries from GS

BPindiv %>%
  filter(!is.na(affiliation_country)) %>%
  filter(affiliation_country  %in% GS_nations) %>%
  count(affiliation_country) %>%
  arrange(desc(n)) %>% select(n) %>% sum() -> n_winners_GS_indiv #number of individual winners from GS

# Table of Count of winners by their country of affiliation
# country_count  %>%
#   mutate(Global_South =  ifelse(affiliation_country %in% GS_nations, "yes", "no")) %>%
#   kable("html") %>%
#   kable_styling("striped", position = "left") #%>%
#   #scroll_box(width = "100%", height = "1500px")
```

Summary of awardees from the Global South:
- Out of 1025extracted individual winners with affiiation, 111 (11%) were affiliated to a country classified as the Global South.
- The number of countries classified as the Global South: 12 (29% of the all detected countries).

Plot awardee counts by country affiliations overall for all countries:

```r
#all countries as a simple barplot
BPindiv %>%
  filter(!is.na(affiliation_country)) %>%
  count(affiliation_country) %>%
  arrange((n)) %>%
  mutate(Global_South =  ifelse(affiliation_country %in% GS_nations,
                               "yes",
                               "no")) %>%
  ggplot(aes(x = reorder(affiliation_country, n),
             y = n,
             fill = Global_South)) +
  geom_bar(stat = "identity",
           position = position_dodge(0.9)) +
  theme_minimal() +
  geom_text(aes(label = n),
            vjust = 0.5,
            hjust = 0,
            nudge_y = 5) + #add counts #add GS status
  scale_y_continuous(limits = c(0, 550)) +
  coord_flip() +
  theme(legend.position="top",
        legend.box = "horizontal",
        axis.text = element_text(size = 10)) +
  labs(fill = "Global South country: ") +
  labs(x = "Affiliation country",
       y = "Awardee count")
```

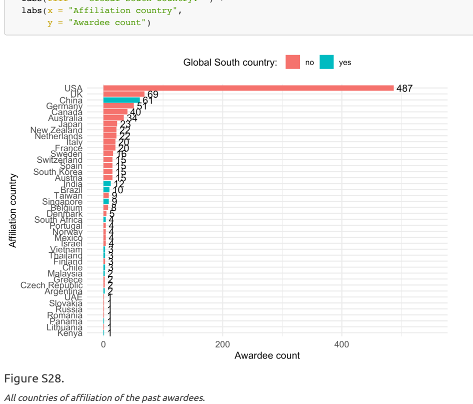


Figure S28.

*All countries of affiliation of the past awardees.*

Plot awardee counts for only top 10 countries with most awardees:

```r
#top 10 countries - two-colour barplot
BPindiv %>%
  filter(!is.na(Country10)) %>%
  count(Country10) %>%
  arrange((n)) %>%
  ggplot(aes(x = reorder(Country10, n), y = n)) +
  geom_bar(aes(fill= Country10 == "other"),
           stat = "identity",
           position = position_dodge(0.9)) +
  scale_fill_manual(values = c("#DCDCDCFF", "#000000")) +
  coord_flip() +
  theme_bw() +
  theme(legend.position="none",
        axis.text = element_text(size = 10)) +
  labs(x = "Affiliation country",
       y = "Awardee count")
```

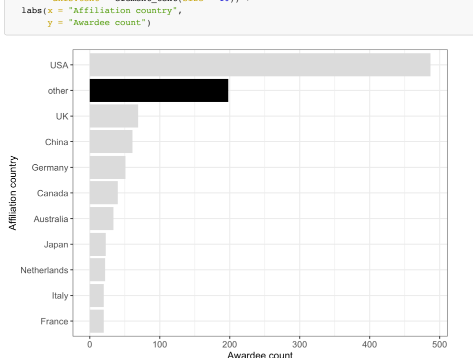


Figure S29.

*Countries of first affiliation of the past awardees - top 10 countries only (remaining countries grouped as "other" and coloured black).*

Plot country of affiliation counts by Subject Area:

```r
#define 11 colour blind-friendly colors from https://rdrr.io/cran/colorBlindness/src/R/palette.R:
mycolors <- rev(c("#FFBF7F", "#FF7F00", "#000000", "#FFFF32",
                  "#B2FF8C", "#A5EDFF", "#19B2FF",
                  "#CCBFFF", "#654CFF", "#FF99BF", "#E51932"))

#top 10 countries by SA
BPindiv %>%
  filter(!is.na(Country10)) %>%
  count(award_SA, Country10) %>%
  ggplot(aes(y = reorder(award_SA, desc(award_SA)),
             x = n,
             fill = Country)) +
  #geom_bar(stat = "identity")
  geom_bar(aes(fill = Country10),
           stat = "identity") +
  scale_fill_manual(values = mycolors) +
  theme_bw() +
  theme(legend.position="top",
        axis.text = element_text(size = 10),
        legend.box = "horizontal",
        legend.margin = margin()) +
  labs(fill = "Awardee affiliation country:") +
  labs(x = "Scimago Subject Area",
       y = "Awardee count")
```

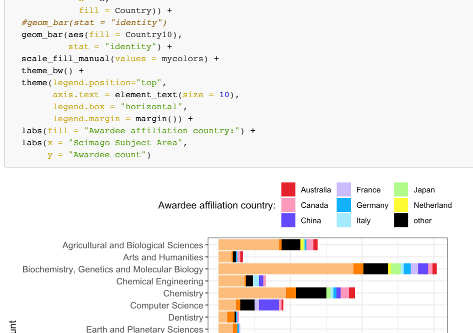


Figure S30.

*Countries of first affiliation of the past awardees - by Subject Area (remaining countries grouped as "other").*

##### Figure 5C - plot of counts of top 10 country affiliations by decade:

```r
#top 10 countries by Decade - horizontal, stacked, with count values
figure5C <- BPindiv %>%
  filter(!is.na(Country10)) %>%
  count(Decade, Country10) %>%
  ggplot(aes(y = Decade,
             x = n,
             fill = Country10)) +
  geom_bar(aes(fill = Country10),
           stat = "identity") + # use , position = "fill" for proportion plot
  scale_fill_manual(values = mycolors) +
  theme_bw() +
  theme(legend.position="top",
        legend.text = element_text(size = 8),
        legend.title = element_text(size = 8),
        axis.text = element_text(size = 10),
        legend.box = "horizontal",
        legend.margin = margin()) +
  guides(fill = guide_legend(nrow = 2, byrow = TRUE, reverse = TRUE)) +
  labs(fill = "Awardee affiliation country:") +
  labs(y = "Decade",
       x = "Awardee counts")
```

##### Figure 5D - proportions of top 10 affiliation countries by decade:

```r
#top 10 countries by Decade - horizontal, stacked, with proportion values
figure5D <- BPindiv %>%
  filter(!is.na(Country10)) %>%
  count(Decade, Country10) %>%
  ggplot(aes(y = Decade,
             x = n,
             fill = Country10)) +
  geom_bar(aes(fill = Country10),
           stat = "identity",
           position = "fill") + # use , position = "fill" for proportion plot
  scale_fill_manual(values = mycolors) +
  theme_bw() +
  theme(legend.position = "none",
        axis.text = element_text(size = 10),
        legend.box = "horizontal",
        legend.margin = margin()) +
  labs(fill = "Awardee affiliation country:") +
  labs(y = "Decade",
       x = "Awardee proportion")
```

Plot awardee country of affiliation counts by decade and by Subject Area:

```r
#top 10 countries by SA and decade
BPindiv %>%
  filter(!is.na(Country10)) %>%
  count(award_SA, Decade, Country10) %>%
  ggplot(aes(y = reorder(award_SA, desc(award_SA)),
             x = n,
             fill = Country10)) +
  geom_bar(aes(fill = Country10),
           stat = "identity") +
  scale_fill_manual(values = mycolors) +
  facet_wrap(~Decade, scales = "fixed", nrow = 1, ncol = 3) +
  theme_bw() +
  theme(legend.position="top",
        axis.text = element_text(size = 10),
        legend.box = "horizontal",
        legend.margin = margin()) +
  labs(fill = "Awardee affiliation country:") +
  labs(x = "Awardee count",
       y = "Scimago Subject Area")
```

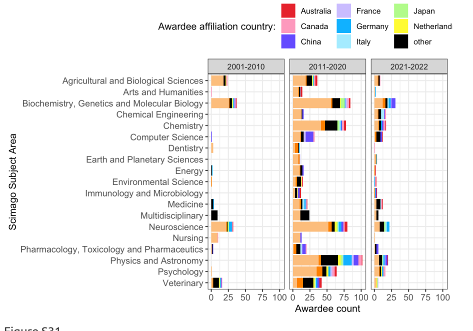


Figure S31.

*Counts of awardees countries of first affiliation - by decade and Subject Area.*

##### Figure 5 - combine 4 panels and save:

```r
#assemble the panels using patchwork package
figure5 <- figure5A / figure5B / figure5C / figure5D +
  plot_layout(ncol = 1, nrow = 4) +
  plot_annotation(tag_levels = "A")
#ggsave(plot = figure5, here("plots", "Fig5ABCD_gender_countries_v1.png"), width = 18, height = 14, units = "cm", d
#ggsave(plot = figure5, here("plots", "Fig5ABCD_gender_countries_v1.pdf"), width = 18, height = 14, units = "cm", s
```

#### Country productivity context.

Load and process SCImago Country document productivity data from 2021:

```r
COprod <- read_csv(here("data", "scimagojr country rank 2021.csv"), skip = 0, show_col_types = FALSE) #load data

#top10 contries
COprod$Country %in% unique(BPindiv$Country10) %>% sum() #check overlap - 8
```

```
## [1] 8
```

```r
#setdiff(unique(BPindiv$Country10), COprod$Country) #missing: "United States" and "United Kingdom"
COprod$Country <-  gsub("United States", "USA", COprod$Country)  #replace with matching name
COprod$Country <-  gsub("United Kingdom", "UK", COprod$Country)  #replace with a matching name
#all countries
COprod$Country %in% unique(BPindiv$affiliation_country) %>% sum() #check overlap - 39
```

```
## [1] 39
```

```
setdiff(unique(BPindiv$affiliation_country), COprod$Country) #missing: "Vietnam" "UAE" "Russia"
```

```
## [1] "Vietnam" NA        "UAE"     "Russia"
```

```
## [1] "Vietnam" NA        "UAE"     "Russia"
COprod$Country <-  gsub("Viet Nam", "Vietnam", COprod$Country)  #replace with matching name
COprod$Country <-  gsub("United Arab Emirates", "UAE", COprod$Country)  #replace with matching name
COprod$Country <-  gsub("Russian Federation", "Russia", COprod$Country)  #replace with a matching name

## Plot for all countries
#recode top 10 countries + other
COprod$Country_affil <- recode(COprod$Country,
                              "USA" = "USA",
                              "UK" = "UK",
                              "Australia" = "Australia",
                              "China" = "China",
                              "Germany" = "Germany",
                              "Canada" = "Canada",
                              "Japan" = "Japan",
                              "Netherlands" = "Netherlands",
                              "Italy" = "Italy",
                              "France" = "France",
                              .default = "other")
#table(COprod_affil$Country_affil)

## make a simple table with percentages of the total number of Documents
# COprod %>%
#   group_by(Country_affil) %>%
#   summarise(Documents = sum(Documents)) %>%
#   mutate(Documents_pct = Documents/sum(Documents) * 100)
```

Plot for the top 10 most productive countries:

```
## Plot productivity of top 10 countries and other - color productivity (Documents) for Countries that are in BPind
COprod %>%
  ggplot(aes(x = 1,
             y = Documents,
             fill = Country_affil)) +
  geom_bar(stat = "identity",
           position = "fill") +
  coord_flip() +
  scale_fill_manual(values = mycolors) +
  theme_bw() +
  theme(legend.position="top",
        axis.text = element_text(size = 10),
        legend.box = "horizontal",
        legend.margin = margin()) +
  labs(x = "Countries",
       y = "Documents",
       fill = "Awardee affiliation country:") +
  scale_x_discrete(labels = NULL)  #used to remove vertical labels, also breaks = NULL
```

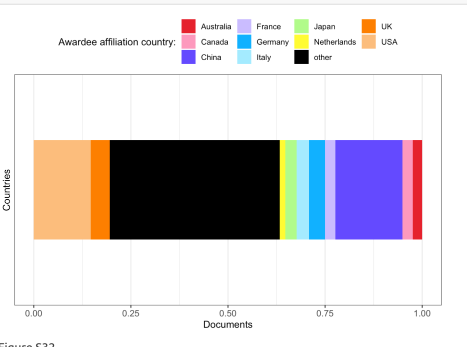


Figure S32.

*SCImago Country document productivity distribution for top 10 countries in the data set (remaining countries are grouped as "other").*

Plot of awardee gender counts by country affiliation by decade:

```
#top 10 countries by SA and decade
BPindiv %>%
  filter(!is.na(Country10)) %>%
  count(Gender, Decade, Country10) %>%
  ggplot(aes(y = reorder(Country10, desc(Country10)),
             x = n,
             fill = Gender)) +
  #geom_bar(stat = "identity")
  geom_bar(aes(fill = Gender),
           stat = "identity") + # use , position = "fill" for proportion plot
  scale_fill_manual(values = c("#CA884C", "#3A5775")) +
  facet_wrap(~Decade, scales = "fixed", nrow = 1, ncol = 3) +
  theme_bw() +
  theme(legend.position="bottom",
        axis.text = element_text(size = 10),
        legend.box = "horizontal",
        legend.margin = margin()) +
  labs(fill = "Awardee gender:") +
  labs(x = "Awardee count",
       y = "Affiliation country")
```

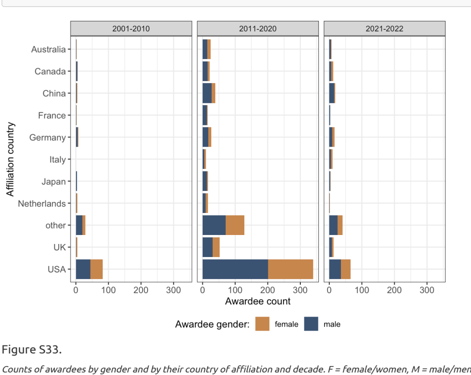


Figure S33.

*Counts of awardees by gender and by their country of affiliation and decade. F = female/women, M = male/men.*

#### Analyses of past awardees first names.

Find the most common first names of the past award winners:

```
BPindiv$first_names <- word(BPindiv$awardee_name, 1) #extract first word
#length(unique(BPindiv$first_names)) #815 unique first names, note some are initials

# #count number of unique first names per decade:
# BPindiv %>%
#   filter(!is.na(first_names)) %>%
#   group_by(Decade) %>%
#   summarise(count = n_distinct(first_names))

# #count number of unique first names per year:
# BPindiv %>%
#   filter(!is.na(first_names)) %>%
#   group_by(award_year) %>%
#   summarise(count = n_distinct(first_names)) %>% View()

#count first_names - view 16 names with >5 counts (16 names):
# BPindiv %>%
#    #filter(!is.na(first_names)) %>%
#    count(first_names) %>%
#    arrange(desc(n)) %>%
#    filter(n > 5)
```

Total number of unique past awardee first names in the dataset: 815.

Plot counts of first awardees names that appear more than 5 times:

```
#plot first names that appear more than 5 times
BPindiv %>%
  #filter(!is.na(first_names)) %>%
  count(first_names) %>%
  arrange(desc(n)) %>%
  filter(n > 5) %>%
  ggplot(aes(x = reorder(first_names, n),
             y = n)) +
  geom_bar(stat = "identity",
           position = position_dodge(0.9)) +
  coord_flip() +
  theme_bw() +
  labs(x = "Winners first name", y = "Count")
```

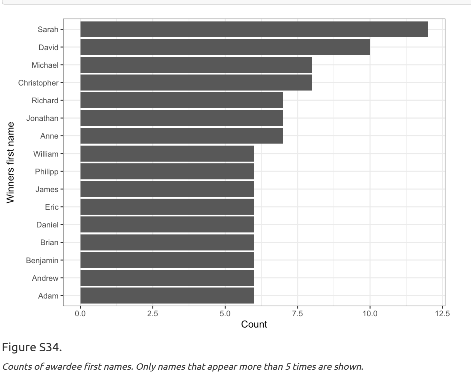


Figure S34.

*Counts of awardee first names. Only names that appear more than 5 times are shown.*

#### Affiliation institutions of past awardees.

Process and find most common affiliation institutions:

```
#length(unique(BPindiv$affiliation_institution)) #672 unique  names, note some require cleaning from extra informat

#do some  cleaning
BPindiv$institution <- BPindiv$affiliation_institution
BPindiv$institution <- gsub(",.*", "", BPindiv$affiliation_institution) #remove everything (usually a city) after f

# #count number of unique institution names per decade:
# BPindiv %>%
#   filter(!is.na(institution)) %>%
#   group_by(Decade) %>%
#   summarise(count = n_distinct(institution))

# #count number of unique institution names per year:
# BPindiv %>%
#   filter(!is.na(institution)) %>%
#   group_by(award_year) %>%
#   summarise(count = n_distinct(institution)) %>% View()

# #count institutions - view institution names with >5 counts (16 names):
# BPindiv %>%
#    filter(!is.na(institution)) %>%
#    count(institution) %>%
#    arrange(desc(n)) %>%
#    filter(n > 5)
```

Total number of unique affiliation institution names in the dataset: 583.

Plot counts of institution names that appear more than 5 times:

```
#plot institution names that appear more than 5 times
BPindiv %>%
  filter(!is.na(institution)) %>%
  count(institution) %>%
  arrange(desc(n)) %>%
  filter(n > 5) %>%
  ggplot(aes(x = reorder(institution, n),
             y = n)) +
  geom_bar(stat = "identity",
           position = position_dodge(0.9)) +
  coord_flip() +
  theme_bw() +
  labs(x = "Winners affiliation institution",
       y = "Count")
```

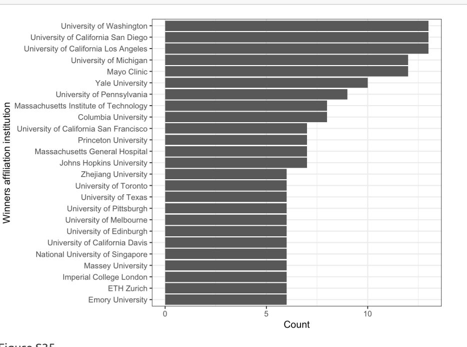


Figure S35.

*Counts of awardee first affiliation institutions. Only institutions that appear more than 5 times are shown.*

#### Cash and perks for the winners.

Cash award mentioned in 112 (50%) award descriptions.

##### Figure 6A - mentions of cash awards:

```
#table(BPdata$Award_cash) #103 no, 111 yes

figure6A <- BPdata %>%
    mutate(Award_cash = factor(Award_cash, levels = (c("yes",
                                                      "not available",
                                                      "no")))) %>% #reorder value levels
    mutate(Award_cash = recode(Award_cash,
                               'no' = 'no',
                               'yes' = 'yes',
                               'not available' = 'no description')) %>% #change level names
    count(Award_cash) %>%
    ggplot(aes(x = 1,
               y = n,
               fill = Award_cash,
               pattern = Award_cash,
               pattern_angle = Award_cash)) +
    geom_bar_pattern(stat = "identity",
    pattern_density = 0.1,
    pattern_size = 0.5,
    pattern_spacing = 0.1,
    pattern_key_scale_factor = 0.2,
    pattern_alpha = 0.5,
    pattern_fill    = 'white',
    pattern_colour  = 'white') +
    coord_flip() +
    scale_y_continuous(breaks = c(0, 50, 100, 150, 200, 250)) +
    theme_classic() +
    scale_fill_manual(values = c("#748A52", "#CCBA98", "#8A6172")) +
    labs(x = "Cash",
         y = "Award count") +
    scale_x_discrete(labels = NULL) + #used to remove vertical labels, also breaks = NULL
    theme(legend.position = "top",
          axis.title.x = element_text(size = 10))
```

Plot of cash awards mentioned per Subject Area:

```
#table(BPdata$Scimago_Subject_Area, BPdata$Award_cash) #as a table by SA

BPdata %>%
    mutate(Award_cash = factor(Award_cash, levels = (c("yes",
                                                      "not available",
                                                      "no")))) %>% #reorder value levels
    mutate(Award_cash = recode(Award_cash,
                               'no' = 'no',
                               'yes' = 'yes',
                               'not available' = 'no description')) %>% #change level names
    count(Scimago_Subject_Area, Award_cash) %>%
    ggplot(aes(x = reorder(Scimago_Subject_Area, desc(Scimago_Subject_Area)),
               y = n,
               fill = Award_cash)) +
    geom_col(width = 0.8,
             position = position_stack(reverse = TRUE)) +
    coord_flip() +
    scale_y_continuous(breaks = c(0, 5, 10)) +
    theme_classic() +
    scale_fill_manual(values = c("#748A52", "#CCBA98", "#8A6172")) +
    labs(x = "Scimago Subject Area",
         y = "Award count",
         fill = "Cash award mentioned:") +
    theme(legend.position = "top",
          axis.title.x = element_text(size = 10))
```

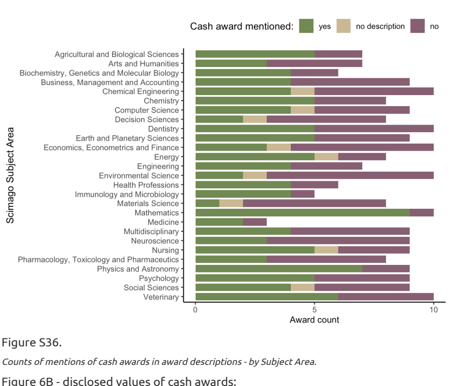


Figure S36.

*Counts of mentions of cash awards in award descriptions - by Subject Area.*

##### Figure 6B - disclosed values of cash awards:

```
#table(!is.na(BPdata$Award_cash_max_USD_pperson)) #96 values available
#median(BPdata$Award_cash_max_USD_pperson[!is.na(BPdata$Award_cash_max_USD_pperson)]) #1086 median
#table(!is.na(BPdata$Award_cash_max_USD_pperson), BPdata$Award_individual) #of 96 values available, 39 are for indi

#wacolors$uw #palette to be used

#plot overall - histogram
figure6B <- BPdata %>%
  filter(!is.na(Award_cash_max_USD_pperson)) %>%
  ggplot(aes(y = Award_cash_max_USD_pperson)) +
  geom_histogram(binwidth = 500,
                 fill = "#748A52",
                 col = "white") +
  coord_flip() +
  theme_bw() +
  labs(y = "Award cash max. USD/person",
       x = "Award count") +
  scale_y_continuous(breaks = c(seq(0, 30000, by = 5000))) +
  theme(legend.position = "none",
        axis.title.x = element_text(size = 10))
```

##### Figure 6 - Combine 2 panels and save:

```
#assemble the panels using patchwork package
figure6 <- figure6A / figure6B +
  plot_layout(nrow = 2, heights = c(1, 3)) +
  plot_annotation(tag_levels = "A")
#ggsave(plot = figure6, here("plots", "Fig6AB_perks_v1.png"), width = 18, height = 8, units = "cm", dpi = "retina",
#ggsave(plot = figure6, here("plots", "Fig6AB_perks_v1.pdf"), width = 18, height = 8, units = "cm", scale = 1.2)
```

Plot of awards with highest cash amounts:

```
BPdata %>%
  filter(!is.na(Award_cash_max_USD_pperson)) %>%
  filter(Award_cash_max_USD_pperson >= 5000) %>%
  ggplot(aes(x = Award_name,
             y = Award_cash_max_USD_pperson)) +
  geom_col(width = 0.8,
           position = position_stack(reverse = TRUE)) +
  coord_flip() +
  theme_bw() +
  labs(y = "Award cash max. USD/person",
       x = "Award name")
```

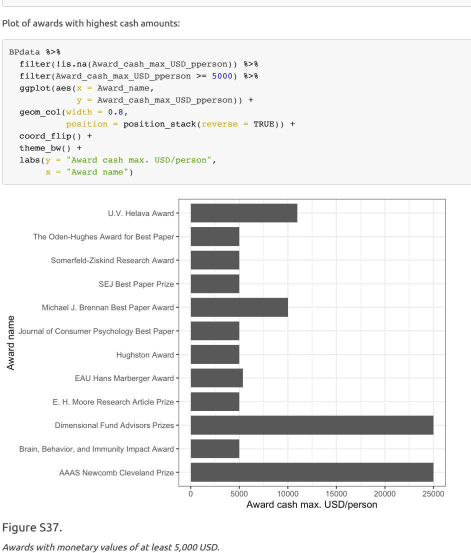


Figure S37.

*Awards with monetary values of at least 5,000 USD.*

Plot disclosed cash amounts by Subject Area:

```
BPdata %>%
  filter(!is.na(Award_cash_max_USD_pperson)) %>%
  ggplot(aes(x = reorder(Scimago_Subject_Area,
                         desc(Scimago_Subject_Area)),
             y = Award_cash_max_USD_pperson)) +
  geom_beeswarm(color = "grey30",
                alpha = 0.5) +
  coord_flip() +
  theme_bw() +
  labs(y = "Award cash max. USD/person",
       x = "Scimago Subject Area") +
  theme(legend.position = "none",
        axis.title.x = element_text(size = 10))
```

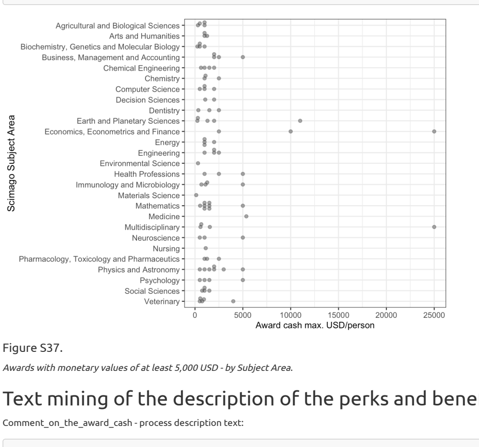


Figure S37.

*Awards with monetary values of at least 5,000 USD - by Subject Area.*

#### Text mining of the description of the perks and benefits.

Comment_on_the_award_cash - process description text:

```
#Using packages tidytext and stopwords
Comment_on_the_award_cash_txt <- tibble(txt = tolower(BPdata$Comment_on_the_award_cash))
Comment_on_the_award_cash_txt <- Comment_on_the_award_cash_txt %>% unnest_tokens(output = word, input = txt, token
Comment_on_the_award_cash_txt <- Comment_on_the_award_cash_txt %>% anti_join(get_stopwords()) #remove stop words
```

```
## Joining with `by = join_by(word)`
```

```
## Joining with `by = join_by(word)`
Comment_on_the_award_cash_txt$word <- tokenize_word_stems(Comment_on_the_award_cash_txt$word) #make all word stems
word.freq <- Comment_on_the_award_cash_txt %>% count(word, sort = TRUE) #count words
word.freq$word <-  gsub("[[:digit:]]", "", word.freq$word)  #remove numbers
word.freq$word <- gsub("[[:punct:][:blank:]]+", " ", word.freq$word) #remove punctuation
word.freq <-  word.freq %>% drop_na() %>% filter(word != "")
```

Count mentions of specific words (stemmed) in Comment_on_the_award_cash:

```
#create list of specific words (stemmed) to count within strings
specific.words2 <- c("certificate", "grant", "attend", "expens", "plaque", "talk", "invit", "registr", "meet", "con

#prepare award descriptions as a single lowercase string
descriptions2 <- BPdata %>%
  filter(!is.na(Comment_on_the_award_cash)) %>%
  select(Comment_on_the_award_cash) %>%
  tolower() #single lowercase string

#sum of all mentions for each word
specific.words.mentions2 <- specific.words2 %>%
  map_int(~ str_count(tolower(descriptions2), .x))

#prepare award descriptions while keeping them separate for each award
descriptions3 <- tolower(BPdata$Comment_on_the_award_cash) #vector of lowercase strings

#sum of mentions per award for each word (counts only one mention per award)
specific.words.mentions3 <- specific.words2 %>%
  map_int(~ sum(str_detect(descriptions3, .x), na.rm = TRUE))

# ## doing the same as above, but manually:
# #count all mentions of words(parts) individually, e.g.:
# sum(str_count(tolower(BPdata$Comment_on_the_award_cash), "certificate"), na.rm = TRUE)
# #counting once per award, e.g.
# sum(str_detect(BPdata$Comment_on_the_award_cash, "certificate"), na.rm = TRUE)
```

Plot frequencies of specific words - all mentions in award cash descriptions:

```
words.df <- tibble(Words = specific.words2,
                   Count_all = specific.words.mentions2,
                   Count_once = specific.words.mentions3)

words.df %>%
    ggplot(aes(x = reorder(Words, Count_all),
               y = Count_all)) +
    geom_col(width = 0.8,
             fill = "#838B83") +
    coord_flip() +
    scale_y_continuous(breaks = c(0, 5, 10, 15, 20, 25, 30, 35, 40, 45, 50, 55),
                       limits = c(0, 55)) +
    theme_bw() +
    labs(x = "Word stem",
         y = "Count of all mentions") +
    theme(legend.position = "none",
          axis.title.x = element_text(size = 10))
```

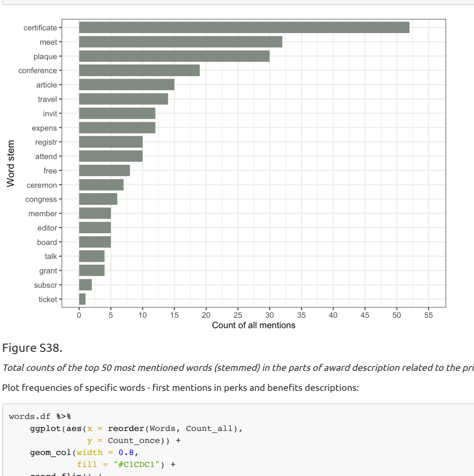


Figure S38.

*Total counts of the top 50 most mentioned words (stemmed) in the parts of award description related to the prize value and related perks.*

Plot frequencies of specific words - first mentions in perks and benefits descriptions:

```
words.df %>%
    ggplot(aes(x = reorder(Words, Count_all),
               y = Count_once)) +
    geom_col(width = 0.8,
             fill = "#C1CDC1") +
    coord_flip() +
    scale_y_continuous(breaks = c(0, 5, 10, 15, 20, 25, 30, 35, 40, 45, 50, 55),
                       limits = c(0, 55)) +
    theme_bw() +
    labs(x = "Word stem",
         y = "Count of first mention per award") +
    #scale_x_discrete(labels = NULL) + labs(x = "")  + #used to remove vertical labels, also breaks = NULL
    theme(legend.position = "none",
          axis.title.x = element_text(size = 10))
```

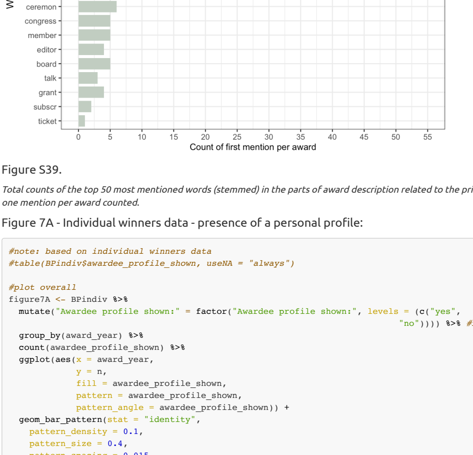


Figure S39.

*Total counts of the top 50 most mentioned words (stemmed) in the parts of award description related to the prize value and related perks - only one mention per award counted.*

##### Figure 7A - Individual winners data - presence of a personal profile:

```
#note: based on individual winners data
#table(BPindiv$awardee_profile_shown, useNA = "always")

#plot overall
figure7A <- BPindiv %>%
  mutate("Awardee profile shown:" = factor("Awardee profile shown:", levels = (c("yes",
                                                                                "no")))) %>% #reorder value levels
  group_by(award_year) %>%
  count(awardee_profile_shown) %>%
  ggplot(aes(x = award_year,
             y = n,
             fill = awardee_profile_shown,
             pattern = awardee_profile_shown,
             pattern_angle = awardee_profile_shown)) +
  geom_bar_pattern(stat = "identity",
    pattern_density = 0.1,
    pattern_size = 0.4,
    pattern_spacing = 0.015,
    pattern_key_scale_factor = 0.5,
    pattern_alpha = 0.5,
    pattern_fill    = 'white',
    pattern_colour  = 'white') +
  scale_fill_manual(values = c("#8A6172", "#748A52")) +
  theme_bw() +
  labs(fill = "Awardee profile shown:",
       pattern = "Awardee profile shown:",
       pattern_angle = "Awardee profile shown:",
       x = "Year",
       y = "Awardee count") +
  theme(legend.position = "top",
        axis.title.x = element_text(size = 10))
```

##### Figure 7B - Individual winners data - presence of a personal photo:

```
#table(BPindiv$awardee_photo_shown, useNA = "always")
#table(BPindiv$awardee_photo_shown, BPindiv$award_year, useNA = "always")

#plot overall
figure7B <- BPindiv %>%
  mutate(awardee_photo_shown = factor(awardee_photo_shown, levels = (c("no",
                                                                      "yes")))) %>% #reorder value levels
  group_by(award_year) %>%
  count(awardee_photo_shown) %>%
  ggplot(aes(x = award_year,
             y = n,
             fill = awardee_photo_shown,
             pattern = awardee_photo_shown,
             pattern_angle = awardee_photo_shown)) +
  geom_bar_pattern(stat = "identity",
    pattern_density = 0.1,
    pattern_size = 0.4,
    pattern_spacing = 0.015,
    pattern_key_scale_factor = 0.5,
    pattern_alpha = 0.5,
    pattern_fill    = 'white',
    pattern_colour  = 'white') +
  scale_fill_manual(values = c("#8A6172", "#748A52")) +
  theme_bw() +
  labs(fill = "Awardee photo shown:",
       pattern = "Awardee photo shown:",
       pattern_angle = "Awardee photo shown:",
       x = "Year",
       y = "Awardee count") +
  theme(legend.position = "top",
        axis.title.x = element_text(size = 10))
```

##### Figure 7 - combine 2 panels and save:

```
#assemble the panels using patchwork package
figure7 <- figure7A / figure7B +
  plot_layout(ncol = 2, nrow = 1) +
  plot_annotation(tag_levels = "A")
#ggsave(plot = figure7, here("plots", "Fig7AB_perks_v1.png"), width = 18, height = 8, units = "cm", dpi = "retina",
#ggsave(plot = figure7, here("plots", "Fig7AB_perks_v1.pdf"), width = 18, height = 8, units = "cm", scale = 1.2)
```

#### Statistical analyses and visualisations of awardees data - using award years.

```
#Scale the award_year variable to have the mean of 0 and SD of 1
BPindiv$Award_year_scaled <- scale(BPindiv$award_year)
```

```
#table(BPindiv$awardee_gender, useNA = "always")
BPindiv %>%
  mutate(Gender = case_when(
    endsWith(awardee_gender, "F") ~ 1,
    endsWith(awardee_gender, "M") ~ 0
    )) -> BPindiv
#table(BPindiv$Gender)

#Fit generalised mixed model with binomial error family and with logit link function, award as a random effect (usi

#without year
model_gender <- glmer(Gender ~ 1 + (1|award_SA) + (1|award_name),
                      family = "binomial",
                      data = BPindiv) #without year as a predictor
summary(model_gender)
```

```
## Generalized linear mixed model fit by maximum likelihood (Laplace
##   Approximation) [glmerMod]
##  Family: binomial  ( logit )
## Formula: Gender ~ 1 + (1 | award_SA) + (1 | award_name)
##    Data: BPindiv
##
##      AIC      BIC   logLik deviance df.resid
##   1422.4   1437.3   -708.2   1416.4     1076
##
## Scaled residuals:
##     Min      1Q  Median      3Q     Max
## -1.4256 -0.7983 -0.6339  1.0853  2.0280
##
## Random effects:
##  Groups     Name        Variance Std.Dev.
##  award_name (Intercept) 0.2275   0.4770
##  award_SA   (Intercept) 0.1647   0.4058
## Number of obs: 1079, groups:  award_name, 61; award_SA, 19
##
```

```
## Fixed effects:
##             Estimate Std. Error z value Pr(>|z|)
## (Intercept)  -0.5093     0.1454  -3.503 0.000461 ***
## ---
## Signif. codes:  0 '***' 0.001 '**' 0.01 '*' 0.05 '.' 0.1 ' ' 1
```

```
plogis(summary(model_gender)$coef[1,1]) *100 #calculate % difference between genders at the intercept: 38% female na
```

```
## [1] 37.53664
```

```
#with year
model_gender <- glmer(Gender ~ Award_year_scaled + (1|award_SA) + (1|award_name),
                      family = "binomial",
                      data = BPindiv) #with year as a predictor
summary(model_gender) #year not significant
```

```
## Generalized linear mixed model fit by maximum likelihood (Laplace
##   Approximation) [glmerMod]
##  Family: binomial  ( logit )
## Formula: Gender ~ Award_year_scaled + (1 | award_SA) + (1 | award_name)
##    Data: BPindiv
##
##      AIC      BIC   logLik deviance df.resid
##   1423.5   1443.4   -707.7   1415.5     1075
##
## Scaled residuals:
##     Min      1Q  Median      3Q     Max
## -1.4546 -0.8070 -0.6205  1.0708  2.1288
##
## Random effects:
##  Groups     Name        Variance Std.Dev.
##  award_name (Intercept) 0.2133   0.4618
##  award_SA   (Intercept) 0.1824   0.4271
## Number of obs: 1079, groups:  award_name, 61; award_SA, 19
##
## Fixed effects:
##                   Estimate Std. Error z value Pr(>|z|)
## (Intercept)       -0.52084    0.14784  -3.523 0.000427 ***
## Award_year_scaled  0.06841    0.07214   0.948 0.342979
## ---
## Signif. codes:  0 '***' 0.001 '**' 0.01 '*' 0.05 '.' 0.1 ' ' 1
##
## Correlation of Fixed Effects:
##             (Intr)
## Awrd_yr_scl -0.082
```

```
plogis(summary(model_gender)$coef[1,1])*100 #calculate % difference between genders at the intercept: 37% female na
```

```
## [1] 37.26562
```

Logistic plot for awardee gender by year:

```
BPindiv %>%
  ggplot(aes(x = award_year,
             y = Gender)) +
  geom_point(alpha = 0.1) +
  stat_smooth(method = "glm",
              method.args = list(family = binomial),
              se = TRUE) +
  xlab("Award year") +
  ylab("Awardee gender") +
  theme_minimal()
```

```
## `geom_smooth()` using formula = 'y ~ x'
```

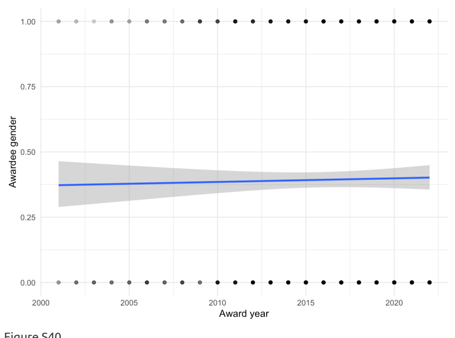


Figure S40.

*Plot of awardee gender, by award year.*

Models for individual winners online profile across years:

```
#table(BPindiv$awardee_profile_shown, useNA = "always")
BPindiv %>%
  mutate(Profile = case_when(
    endsWith(awardee_profile_shown, "yes") ~ 1,
    endsWith(awardee_profile_shown, "no") ~ 0
    )) -> BPindiv

#Fit generalised mixed model with binomial error family and with logit link function, award as a random effect (usi
model_profile <- glmer(Profile ~ Award_year_scaled + (1|award_name),
                       family = "binomial",
                       data = BPindiv) #with year as a predictor
summary(model_profile) #slope 1.11 signif
```

```
## Generalized linear mixed model fit by maximum likelihood (Laplace
##   Approximation) [glmerMod]
##  Family: binomial  ( logit )
## Formula: Profile ~ Award_year_scaled + (1 | award_name)
##    Data: BPindiv
##
##      AIC      BIC   logLik deviance df.resid
##    383.9    398.8   -188.9    377.9     1076
##
## Scaled residuals:
##     Min      1Q  Median      3Q     Max
## -3.7555 -0.0157 -0.0097 -0.0052 10.5452
##
## Random effects:
##  Groups     Name        Variance Std.Dev.
##  award_name (Intercept) 94.32    9.712
## Number of obs: 1079, groups:  award_name, 61
##
## Fixed effects:
##                   Estimate Std. Error z value Pr(>|z|)
## (Intercept)        -9.0425     1.4326  -6.312 2.75e-10 ***
## Award_year_scaled   1.1087     0.2231   4.970 6.70e-07 ***
## ---
## Signif. codes:  0 '***' 0.001 '**' 0.01 '*' 0.05 '.' 0.1 ' ' 1
##
## Correlation of Fixed Effects:
##             (Intr)
## Awrd_yr_scl -0.155
```

Logistic plot for awardee online profile by year:

```
BPindiv %>%
  ggplot(aes(x = award_year,
             y = Profile)) +
  geom_point(alpha = 0.1) +
  stat_smooth(method = "glm",
              method.args = list(family = binomial),
              se = TRUE) +
  xlab("Award year") +
  ylab("Awardee online profile") +
  theme_minimal()
```

```
## `geom_smooth()` using formula = 'y ~ x'
```

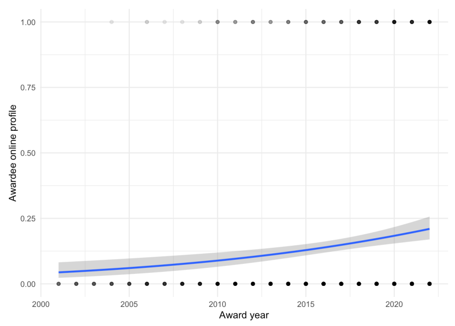


Figure S41.

*Plot of awardee online profile, by award year.*

Models for individual winners online photo across years:

```
#table(BPindiv$awardee_photo_shown, useNA = "always")
BPindiv %>%
  mutate(Photo = case_when(
    endsWith(awardee_photo_shown, "yes") ~ 1,
    endsWith(awardee_photo_shown, "no") ~ 0
    )) -> BPindiv

#Fit generalised mixed model with binomial error family and with logit link function, award as a random effect (usi
model_photo <- glmer(Photo ~ Award_year_scaled + (1|award_name),
                     family = "binomial",
                     data = BPindiv) #with year as a predictor
summary(model_photo) #slope 1.18 signif
```

```
## Generalized linear mixed model fit by maximum likelihood (Laplace
##   Approximation) [glmerMod]
##  Family: binomial  ( logit )
## Formula: Photo ~ Award_year_scaled + (1 | award_name)
##    Data: BPindiv
##
##      AIC      BIC   logLik deviance df.resid
##    426.5    441.5   -210.3    420.5     1076
##
## Scaled residuals:
##      Min       1Q   Median       3Q      Max
## -13.2419  -0.0148  -0.0037   0.0755   2.0921
##
## Random effects:
##  Groups     Name        Variance Std.Dev.
##  award_name (Intercept) 151.6    12.31
## Number of obs: 1079, groups:  award_name, 61
##
## Fixed effects:
##                   Estimate Std. Error z value Pr(>|z|)
## (Intercept)        -8.9908     1.5487  -5.806 6.41e-09 ***
## Award_year_scaled   1.2244     0.2215   5.528 3.24e-08 ***
## ---
## Signif. codes:  0 '***' 0.001 '**' 0.01 '*' 0.05 '.' 0.1 ' ' 1
##
## Correlation of Fixed Effects:
##             (Intr)
## Awrd_yr_scl -0.170
```

Logistic plot for awardee online photo by year:

```
BPindiv %>%
  ggplot(aes(x = award_year,
             y = Photo)) +
  geom_point(alpha = 0.1) +
  stat_smooth(method = "glm",
              method.args = list(family = binomial),
              se = TRUE) +
  xlab("Award year") +
  ylab("Awardee online photo") +
  theme_minimal()
```

```
## `geom_smooth()` using formula = 'y ~ x'
```

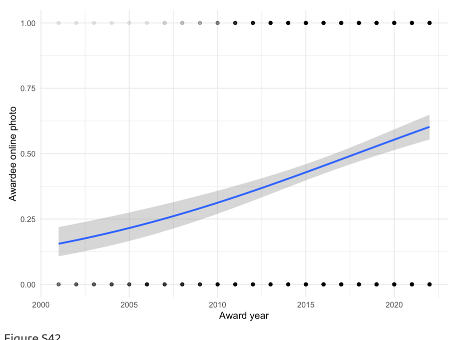


Figure S42.

*Plot of awardee online photo, by award year.*

### Analyses of awards - using award age.

NOTE: Using the year of oldest listed past winner as a proxy of award establishment year.

Award_individual vs. award age:

```
#Scale the Awardee_list_earliest_year variable to have the mean of 0 and SD of 1
BPdata$Awardee_list_earliest_year_scaled <- scale(BPdata$Awardee_list_earliest_year)
#table(!is.na(BPdata$Awardee_list_earliest_year)) # 203 values available

#table
# table(BPdata$Award_individual, useNA = "always")

#filter and make binomial
BPdata %>%
  filter(!is.na(Awardee_list_earliest_year)) %>%
  mutate(Award_individual_num = recode(Award_individual,
                                       yes = 1,
                                       no = 0)) -> BPdata4
```

Logistic plot for individual awards relatively to award age:

```
BPdata4 %>%
  ggplot(aes(x = Awardee_list_earliest_year,
             y = Award_individual_num)) +
  geom_point(alpha = 0.2) +
  stat_smooth(method = "glm",
              method.args = list(family = binomial),
              se = TRUE) +
  xlab("Year of first listed awardee") +
  ylab("Probability of Award_individual") +
  theme_minimal()
```

```
## `geom_smooth()` using formula = 'y ~ x'
```

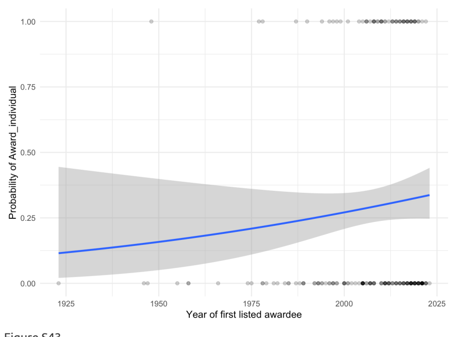


Figure S43.

*Plot of whether award is individual-focused, by award age.*

Models:

```
#model - fit a generalised mixed model with binomial error family and with logit link function, award as a random e
model_Award_individual <- glmer(Award_individual_num ~ Awardee_list_earliest_year_scaled + (1|Scimago_Subject_Area)
                                family = "binomial",
                                data = BPdata4) #with year as a predictor
summary(model_Award_individual) #slope ns
```

```
## Generalized linear mixed model fit by maximum likelihood (Laplace
##   Approximation) [glmerMod]
##  Family: binomial  ( logit )
## Formula: Award_individual_num ~ Awardee_list_earliest_year_scaled + (1 |
##     Scimago_Subject_Area) + (1 | Awarding_society)
##    Data: BPdata4
##
##      AIC      BIC   logLik deviance df.resid
##    229.9    243.1   -110.9    221.9      199
##
## Scaled residuals:
##     Min      1Q  Median      3Q     Max
## -1.3911 -0.5661 -0.2933  0.6713  2.0516
##
## Random effects:
##  Groups               Name        Variance Std.Dev.
##  Awarding_society     (Intercept) 0.2334   0.4831
##  Scimago_Subject_Area (Intercept) 2.3012   1.5170
## Number of obs: 203, groups:  Awarding_society, 118; Scimago_Subject_Area, 27
##
## Fixed effects:
##                                   Estimate Std. Error z value Pr(>|z|)
## (Intercept)                        -1.0512     0.3854  -2.727  0.00639 **
## Awardee_list_earliest_year_scaled   0.2523     0.2318   1.088  0.27650
## ---
## Signif. codes:  0 '***' 0.001 '**' 0.01 '*' 0.05 '.' 0.1 ' ' 1
##
## Correlation of Fixed Effects:
##             (Intr)
## Awrd_lst___ 0.040
```

```
##alternative model reporting (as a table)
#library(sjPlot)
#tab_model(model_Award_individual)
```

Flexible_eligibility vs. award age:

```
#table
# table(BPdata$Flexible_eligibility, useNA = "always")

#filter and make binomial
BPdata %>%
  filter(!is.na(Awardee_list_earliest_year)) %>%
  filter(Flexible_eligibility == "yes" | Flexible_eligibility == "no") %>%
  mutate(Flexible_eligibility_num = recode(Flexible_eligibility,
                                           yes = 1,
                                           no = 0)) -> BPdata4
```

Logistic plot for award flexibility relatvely to award age:

```
BPdata4 %>%
  ggplot(aes(x = Awardee_list_earliest_year,
             y = Flexible_eligibility_num)) +
  geom_point(alpha = 0.2) +
  stat_smooth(method = "glm",
              method.args = list(family = binomial),
              se = TRUE) +
  xlab("Year of first listed awardee") +
  ylab("Probability of Flexible_eligibility") +
  theme_minimal()
```

```
## `geom_smooth()` using formula = 'y ~ x'
```

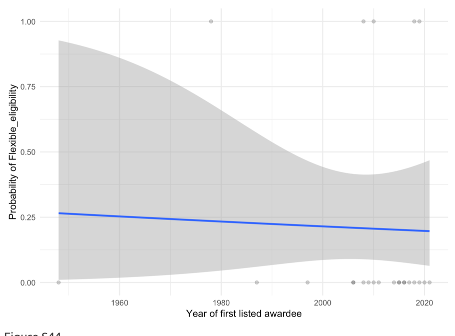


Figure S44.

*Plot of whether award has flexible eligibility, by award age.*

Models:

```
#model - fit a generalised mixed model with binomial error family and with logit link function, award as a random e

# model_Flexible_eligibility <- glmer(Flexible_eligibility_num ~ Awardee_list_earliest_year_scaled + (1|Scimago_Sub

model_Flexible_eligibility <- glm(Flexible_eligibility_num ~ Awardee_list_earliest_year_scaled,
                                  family = "binomial",
                                  data = BPdata4) #with year as a predictor, without random effects included
summary(model_Flexible_eligibility) #slope ns
```

```
##
## Call:
## glm(formula = Flexible_eligibility_num ~ Awardee_list_earliest_year_scaled,
##     family = "binomial", data = BPdata4)
##
## Coefficients:
##                                   Estimate Std. Error z value Pr(>|z|)
## (Intercept)                       -1.32990    0.50324  -2.643  0.00823 **
## Awardee_list_earliest_year_scaled -0.08686    0.48362  -0.180  0.85747
## ---
## Signif. codes:  0 '***' 0.001 '**' 0.01 '*' 0.05 '.' 0.1 ' ' 1
##
## (Dispersion parameter for binomial family taken to be 1)
##
##     Null deviance: 24.564  on 23  degrees of freedom
## Residual deviance: 24.532  on 22  degrees of freedom
## AIC: 28.532
##
## Number of Fisher Scoring iterations: 4
```

Assessors_transparency vs. award age:

```
#table
# table(BPdata$Assessors_transparency, useNA = "always")

#filter and make binomial
BPdata %>%
  filter(!is.na(Awardee_list_earliest_year)) %>%
  filter(Assessors_transparency == "yes" | Assessors_transparency == "no") %>%
  mutate(Assessors_transparency_num = recode(Assessors_transparency,
                                             yes = 1,
                                             no = 0)) -> BPdata4
```

Logistic plot of assesor transparency versus award age:

```
BPdata4 %>%
  ggplot(aes(x = Awardee_list_earliest_year,
             y = Assessors_transparency_num)) +
  geom_point(alpha = 0.2) +
  stat_smooth(method = "glm",
              method.args = list(family = binomial),
              se = TRUE) +
  xlab("Year of first listed awardee") +
  ylab("Probability of Assessors_transparency") +
  theme_minimal()
```

```
## `geom_smooth()` using formula = 'y ~ x'
```

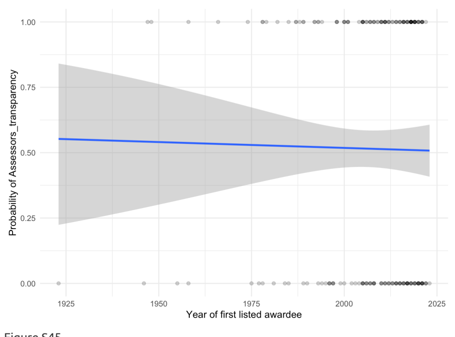


Figure S45.

*Plot of whether award has assessor transparency, by award age.*

Models:

```
#model - fit a generalised mixed model with binomial error family and with logit link function, award as a random e
model_Assessors_transparency <- glmer(Assessors_transparency_num ~ Awardee_list_earliest_year_scaled + (1|Scimago_S
                                      family = "binomial",
                                      data = BPdata4) #with year as a predictor, + (1|Awarding_society) causes "bou
summary(model_Assessors_transparency) #slope ns
```

```
## Generalized linear mixed model fit by maximum likelihood (Laplace
##   Approximation) [glmerMod]
##  Family: binomial  ( logit )
## Formula: Assessors_transparency_num ~ Awardee_list_earliest_year_scaled +
##     (1 | Scimago_Subject_Area)
##    Data: BPdata4
##
##      AIC      BIC   logLik deviance df.resid
##    272.5    282.3   -133.3    266.5      191
##
## Scaled residuals:
##     Min      1Q  Median      3Q     Max
## -1.4059 -0.9351  0.7128  0.9220  1.2300
##
## Random effects:
##  Groups               Name        Variance Std.Dev.
##  Scimago_Subject_Area (Intercept) 0.2852   0.534
## Number of obs: 194, groups:  Scimago_Subject_Area, 27
##
## Fixed effects:
##                                   Estimate Std. Error z value Pr(>|z|)
## (Intercept)                        0.06852    0.18168   0.377    0.706
## Awardee_list_earliest_year_scaled  0.02803    0.15627   0.179    0.858
##
## Correlation of Fixed Effects:
##             (Intr)
## Awrd_lst___ 0.012
```

Award_integrity_mentioned vs. award age:

```
# table(BPdata$Award_integrity_mentioned, useNA = "always")

#filter and make binomial
BPdata %>%
  filter(!is.na(Awardee_list_earliest_year)) %>%
  filter(Award_integrity_mentioned == "yes" | Award_integrity_mentioned == "no") %>%
  mutate(Award_integrity_mentioned_num = recode(Award_integrity_mentioned,
                                                yes = 1,
                                                no = 0)) -> BPdata4
```

Logistic plot of award integrity vrelatively to award age:

```
BPdata4 %>%
  ggplot(aes(x = Awardee_list_earliest_year,
             y = Award_integrity_mentioned_num)) +
  geom_point(alpha = 0.2) +
  stat_smooth(method = "glm",
              method.args = list(family = binomial),
              se = TRUE) +
  xlab("Year of first listed awardee") +
  ylab("Probability of Award_integrity_mentioned") +
  theme_minimal()
```

```
## `geom_smooth()` using formula = 'y ~ x'
```

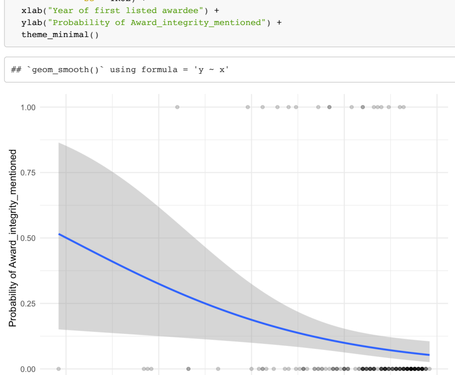


Figure S46.

*Plot of whether award mentions assessment integrity, by award age.*

Models:

```
#model - fit a generalised mixed model with binomial error family and with logit link function, award as a random e
model_Award_integrity_mentioned <- glmer(Award_integrity_mentioned_num ~ Awardee_list_earliest_year_scaled + (1|Awa
                                         family = "binomial",
                                         data = BPdata4) #with year as a predictor, + (1|Scimago_Subject_Area) caus
summary(model_Award_integrity_mentioned) #slope ns, but close
```

```
## Generalized linear mixed model fit by maximum likelihood (Laplace
##   Approximation) [glmerMod]
##  Family: binomial  ( logit )
## Formula: Award_integrity_mentioned_num ~ Awardee_list_earliest_year_scaled +
##     (1 | Awarding_society)
##    Data: BPdata4
##
##      AIC      BIC   logLik deviance df.resid
##     85.6     95.4    -39.8     79.6      190
##
## Scaled residuals:
##     Min      1Q  Median      3Q     Max
## -1.1129 -0.0979 -0.0037 -0.0023  4.6810
##
## Random effects:
##  Groups           Name        Variance Std.Dev.
##  Awarding_society (Intercept) 559.1    23.64
## Number of obs: 193, groups:  Awarding_society, 115
##
## Fixed effects:
##                                   Estimate Std. Error z value Pr(>|z|)
## (Intercept)                       -11.6213     2.6581  -4.372 1.23e-05 ***
## Awardee_list_earliest_year_scaled  -1.1068     0.7514  -1.473    0.141
## ---
## Signif. codes:  0 '***' 0.001 '**' 0.01 '*' 0.05 '.' 0.1 ' ' 1
##
## Correlation of Fixed Effects:
##             (Intr)
## Awrd_lst___ 0.657
```

Self_nomination allowed vs. award age:

```
# table(BPdata$Self_nomination, useNA = "always")

#filter and make binomial
BPdata %>%
  filter(!is.na(Awardee_list_earliest_year)) %>%
  filter(Self_nomination == "yes" | Self_nomination == "no") %>%
  mutate(Self_nomination_num = recode(Self_nomination,
                                      yes = 1,
                                      no = 0)) -> BPdata4
```

Logistic plot of self nominations relatively to award age:

```
BPdata4  %>%
  ggplot(aes(x = Awardee_list_earliest_year,
             y = Self_nomination_num)) +
  geom_point(alpha = 0.2) +
  stat_smooth(method = "glm",
              method.args = list(family = binomial),
              se = TRUE) +
  xlab("Year of first listed awardee") +
  ylab("Probability of Self_nomination") +
  theme_minimal()
```

```
## `geom_smooth()` using formula = 'y ~ x'
```

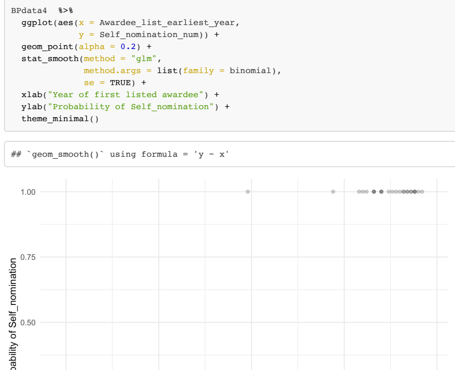


Figure S47.

*Plot of whether award explicitly allows self-nominations, by award age.*

Models:

```
#model - fit a generalised mixed model with binomial error family and with logit link function, award as a random e
model_Self_nomination <- glmer(Self_nomination_num ~ Awardee_list_earliest_year_scaled + (1|Scimago_Subject_Area),
                               family = "binomial",
                               data = BPdata4) #with year as a predictor
summary(model_Self_nomination) #slope ns, but trend for newer awards to mention self-nominations
```

```
## Generalized linear mixed model fit by maximum likelihood (Laplace
##   Approximation) [glmerMod]
##  Family: binomial  ( logit )
## Formula: Self_nomination_num ~ Awardee_list_earliest_year_scaled + (1 |
##     Scimago_Subject_Area)
##    Data: BPdata4
##
##      AIC      BIC   logLik deviance df.resid
##    154.0    163.8    -74.0    148.0      190
##
## Scaled residuals:
##     Min      1Q  Median      3Q     Max
## -0.6315 -0.4125 -0.3501 -0.2592  4.8781
##
## Random effects:
##  Groups               Name        Variance Std.Dev.
##  Scimago_Subject_Area (Intercept) 0.3627   0.6023
## Number of obs: 193, groups:  Scimago_Subject_Area, 27
##
## Fixed effects:
##                                   Estimate Std. Error z value Pr(>|z|)
## (Intercept)                        -2.0653     0.3095  -6.673 2.51e-11 ***
## Awardee_list_earliest_year_scaled   0.5658     0.3225   1.754   0.0793 .
## ---
## Signif. codes:  0 '***' 0.001 '**' 0.01 '*' 0.05 '.' 0.1 ' ' 1
##
## Correlation of Fixed Effects:
##             (Intr)
## Awrd_lst___ -0.343
```

Letter_required vs. award age:

```
# table(BPdata$Letter_required, useNA = "always")

#filter and make binomial
BPdata %>%
```

```
  filter(!is.na(Awardee_list_earliest_year)) %>%
  filter(Letter_required == "yes" | Letter_required == "no") %>%
  mutate(Letter_required_num = recode(Letter_required,
                                      yes = 1,
                                      no = 0)) -> BPdata4
```

Logistic plot for nomination letters relatively to award age:

```
BPdata4  %>%
  ggplot(aes(x = Awardee_list_earliest_year,
             y = Letter_required_num)) +
  geom_point(alpha = 0.2) +
  stat_smooth(method = "glm",
              method.args = list(family = binomial),
              se = TRUE) +
  xlab("Year of first listed awardee") +
  ylab("Probability of Letter_required") +
  theme_minimal()
```

```
## `geom_smooth()` using formula = 'y ~ x'
```

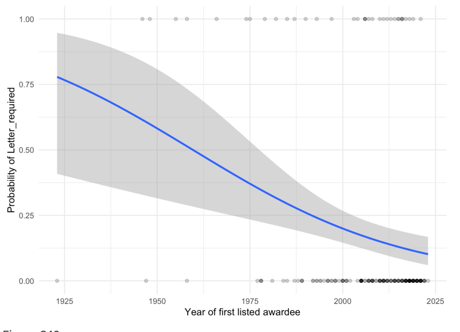


Figure S48.

*Plot of whether award nomination requires a letter, by award age.*

Models:

```
#model - fit a generalised mixed model with binomial error family and with logit link function, award as a random e
model_Letter_required <- glmer(Letter_required_num ~ Awardee_list_earliest_year_scaled + (1|Scimago_Subject_Area) +
                              family = "binomial",
                              data = BPdata4) #with year as a predictor
summary(model_Letter_required) #slope significant negative -0.5431 - less common over the years
```

```
## Generalized linear mixed model fit by maximum likelihood (Laplace
##   Approximation) [glmerMod]
##  Family: binomial  ( logit )
## Formula: Letter_required_num ~ Awardee_list_earliest_year_scaled + (1 |
##     Scimago_Subject_Area) + (1 | Awarding_society)
##    Data: BPdata4
##
##      AIC      BIC   logLik deviance df.resid
##    176.8    189.9    -84.4    168.8      189
##
## Scaled residuals:
##     Min      1Q  Median      3Q     Max
## -1.6478 -0.4311 -0.3374 -0.2687  2.6569
##
## Random effects:
##  Groups               Name        Variance Std.Dev.
##  Awarding_society     (Intercept) 0.1654   0.4067
##  Scimago_Subject_Area (Intercept) 0.5556   0.7454
## Number of obs: 193, groups:  Awarding_society, 115; Scimago_Subject_Area, 27
##
## Fixed effects:
##                                   Estimate Std. Error z value Pr(>|z|)
## (Intercept)                        -1.7578     0.3110  -5.653 1.58e-08 ***
## Awardee_list_earliest_year_scaled  -0.5425     0.1929  -2.812  0.00492 **
## ---
## Signif. codes:  0 '***' 0.001 '**' 0.01 '*' 0.05 '.' 0.1 ' ' 1
##
## Correlation of Fixed Effects:
##             (Intr)
## Awrd_lst___ 0.208
```

Award_contact_provided vs. award age:

```
# table(BPdata$Award_contact_provided, useNA = "always")

#filter and make binomial
BPdata %>%
  filter(!is.na(Awardee_list_earliest_year)) %>%
  filter(Award_contact_provided == "yes" | Award_contact_provided == "no") %>%
  mutate(Award_contact_provided_num = recode(Award_contact_provided,
                                             yes = 1,
                                             no = 0)) -> BPdata4
```

Logistic plot of award contact details relatively to award age:

```
BPdata4  %>%
  ggplot(aes(x = Awardee_list_earliest_year,
             y = Award_contact_provided_num)) +
  geom_point(alpha = 0.2) +
  stat_smooth(method = "glm",
              method.args = list(family = binomial),
              se = TRUE) +
  xlab("Year of first listed awardee") +
  ylab("Probability of Award_individual") +
  theme_minimal()
```

```
## `geom_smooth()` using formula = 'y ~ x'
```

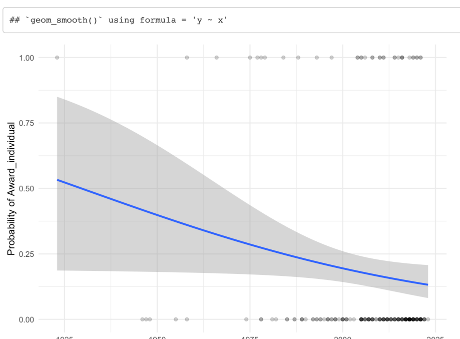


Figure S49.

*Plot of whether award contact details are provided, by award age.*

Models:

```
#model - fit a generalised mixed model with binomial error family and with logit link function, award as a random e
model_Award_contact_provided <- glm(Award_contact_provided_num ~ Awardee_list_earliest_year_scaled,
                                   family = "binomial",
                                   data = BPdata4) #with year as a predictor - but fails with random effects
summary(model_Award_contact_provided) #slope -0.3314
```

```
##
## Call:
## glm(formula = Award_contact_provided_num ~ Awardee_list_earliest_year_scaled,
##     family = "binomial", data = BPdata4)
##
## Coefficients:
##                                   Estimate Std. Error z value Pr(>|z|)
## (Intercept)                        -1.5473     0.1923  -8.048 8.41e-16 ***
## Awardee_list_earliest_year_scaled  -0.3283     0.1625  -2.021   0.0433 *
## ---
## Signif. codes:  0 '***' 0.001 '**' 0.01 '*' 0.05 '.' 0.1 ' ' 1
##
## (Dispersion parameter for binomial family taken to be 1)
##
##     Null deviance: 182.74  on 192  degrees of freedom
## Residual deviance: 178.87  on 191  degrees of freedom
## AIC: 182.87
##
## Number of Fisher Scoring iterations: 4
```

Criteria_transparency vs. award age:

```
# table(BPdata$Criteria_transparency, useNA = "always")

#filter and make binomial
BPdata %>%
filter(!is.na(Awardee_list_earliest_year)) %>%
  filter(Criteria_transparency == "yes" | Criteria_transparency == "no") %>%
  mutate(Criteria_transparency_num = recode(Criteria_transparency,
                                            yes = 1,
                                            no = 0)) -> BPdata4
```

Logistic plot of criteria transparency vs. award age:

```
BPdata4 %>%
  ggplot(aes(x = Awardee_list_earliest_year,
             y = Criteria_transparency_num)) +
  geom_point(alpha = 0.2) +
  stat_smooth(method = "glm",
              method.args = list(family = binomial),
              se = TRUE) +
  xlab("Year of first listed awardee") +
  ylab("Probability of Criteria_transparency") +
  theme_minimal()
```

```
## `geom_smooth()` using formula = 'y ~ x'
```

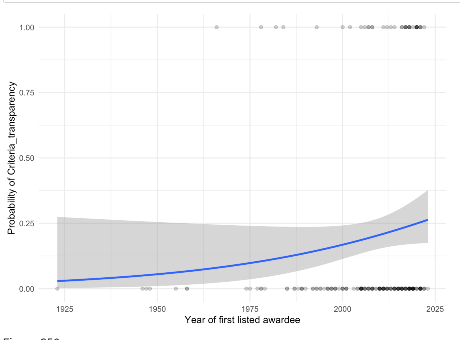


Figure S50.

*Plot of whether award has transparent assessment criteria, by award age.*

Models:

```
#model - fit a generalised mixed model with binomial error family and with logit link function, award as a random e
model_Criteria_transparency <- glm(Criteria_transparency_num ~ Awardee_list_earliest_year_scaled,
                                  family = "binomial",
                                  data = BPdata4) #with year as a predictor
summary(model_Criteria_transparency) #slope ns
```

```
##
## Call:
## glm(formula = Criteria_transparency_num ~ Awardee_list_earliest_year_scaled,
##     family = "binomial", data = BPdata4)
##
## Coefficients:
##                                   Estimate Std. Error z value Pr(>|z|)
## (Intercept)                        -1.4415     0.1891  -7.623 2.48e-14 ***
## Awardee_list_earliest_year_scaled   0.4048     0.2412   1.678   0.0934 .
## ---
## Signif. codes:  0 '***' 0.001 '**' 0.01 '*' 0.05 '.' 0.1 ' ' 1
##
## (Dispersion parameter for binomial family taken to be 1)
##
##     Null deviance: 191.04  on 191  degrees of freedom
## Residual deviance: 187.57  on 190  degrees of freedom
## AIC: 191.57
##
## Number of Fisher Scoring iterations: 4
```

Award_impact_metrics_mentioned vs. award age:

```
# table(BPdata$Award_impact_metrics_mentioned, useNA = "always")
BPdata %>%
  filter(!is.na(Awardee_list_earliest_year)) %>%
  filter(Award_impact_metrics_mentioned == "yes" | Award_impact_metrics_mentioned == "no") %>%
  mutate(Award_impact_metrics_mentioned_num = recode(Award_impact_metrics_mentioned,
                                                     yes = 1,
                                                     no = 0)) -> BPdata4
```

Logistic plot of mentios of impact metrics relatively to teh award age:

```
BPdata4 %>%
  ggplot(aes(x = Awardee_list_earliest_year,
             y = Award_impact_metrics_mentioned_num)) +
  geom_point(alpha = 0.2) +
  stat_smooth(method = "glm",
              method.args = list(family = binomial),
              se = TRUE) +
  xlab("Year of first listed awardee") +
  ylab("Probability of Award_impact_metrics_mentioned") +
  theme_minimal()
```

```
## `geom_smooth()` using formula = 'y ~ x'
```

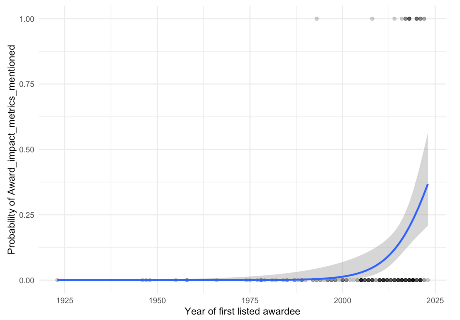


Figure S51.

*Plot of whether award description mentions impact metrics, by award age.*

Models:

```
#model - fit a generalised mixed model with binomial error family and with logit link function, award as a random e
model_Award_impact_metrics_mentioned <- glmer(Award_impact_metrics_mentioned_num ~ Awardee_list_earliest_year_scale
                                             family = "binomial",
                                             data = BPdata4) #with year as a predictor
summary(model_Award_impact_metrics_mentioned) #slope signif 4.930 - recently mentioned more often
```

```
## Generalized linear mixed model fit by maximum likelihood (Laplace
##   Approximation) [glmerMod]
##  Family: binomial  ( logit )
## Formula:
## Award_impact_metrics_mentioned_num ~ Awardee_list_earliest_year_scaled +
##     (1 | Awarding_society)
##    Data: BPdata4
##
##      AIC      BIC   logLik deviance df.resid
##     90.1     99.9    -42.1     84.1      190
##
## Scaled residuals:
##      Min       1Q   Median       3Q      Max
## -0.99218 -0.01514 -0.00158 -0.00009  2.46961
##
## Random effects:
##  Groups           Name        Variance Std.Dev.
##  Awarding_society (Intercept) 916.8    30.28
## Number of obs: 193, groups:  Awarding_society, 115
##
## Fixed effects:
##                                   Estimate Std. Error z value Pr(>|z|)
## (Intercept)                        -14.264      3.262  -4.373 1.23e-05 ***
## Awardee_list_earliest_year_scaled    4.870      2.256   2.158   0.0309 *
## ---
## Signif. codes:  0 '***' 0.001 '**' 0.01 '*' 0.05 '.' 0.1 ' ' 1
##
## Correlation of Fixed Effects:
##             (Intr)
## Awrd_lst___ -0.570
```

Award_impact_metrics_only vs. award age:

```
# table(BPdata$Award_impact_metrics_only, useNA = "always")

#filter and make binomial
BPdata %>%
  filter(!is.na(Awardee_list_earliest_year)) %>%
  filter(Award_impact_metrics_only == "yes" | Award_impact_metrics_only == "no") %>%
  mutate(Award_impact_metrics_only_num = recode(Award_impact_metrics_only,
                                                yes = 1,
                                                no = 0)) -> BPdata4
```

Logistic plot of impact metrics as only criteria relatively to award age:

```
BPdata4 %>%
  ggplot(aes(x = Awardee_list_earliest_year,
             y = Award_impact_metrics_only_num)) +
  geom_point(alpha = 0.2) +
  stat_smooth(method = "glm",
              method.args = list(family = binomial),
              se = TRUE) +
  xlab("Year of first listed awardee") +
  ylab("Probability of Award_impact_metrics_only") +
  theme_minimal()
```

```
## `geom_smooth()` using formula = 'y ~ x'
```

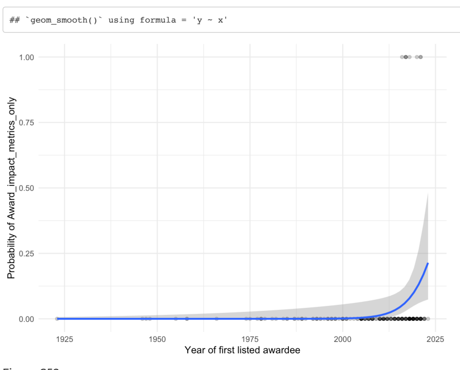


Figure S52.

*Plot of whether award description mentions impact metrics as only criteria, by award age.*

Models:

```
#model - fit a generalised mixed model with binomial error family and with logit link function, award as a random e
model_Award_impact_metrics_only <- glm(Award_impact_metrics_only_num ~ Awardee_list_earliest_year_scaled,
                                      family = "binomial",
                                      data = BPdata4) #with year as a predictor, fails with random effects
summary(model_Award_impact_metrics_only) #slope signif 3.782 - more common recently
```

```
##
## Call:
## glm(formula = Award_impact_metrics_only_num ~ Awardee_list_earliest_year_scaled,
##     family = "binomial", data = BPdata4)
##
## Coefficients:
##                                   Estimate Std. Error z value Pr(>|z|)
## (Intercept)                         -5.154      1.289  -3.998  6.4e-05 ***
## Awardee_list_earliest_year_scaled    3.784      1.705   2.219   0.0265 *
## ---
## Signif. codes:  0 '***' 0.001 '**' 0.01 '*' 0.05 '.' 0.1 ' ' 1
##
## (Dispersion parameter for binomial family taken to be 1)
##
##     Null deviance: 66.596  on 192  degrees of freedom
## Residual deviance: 54.872  on 191  degrees of freedom
## AIC: 58.872
##
## Number of Fisher Scoring iterations: 8
```

Open_science vs. award age:

```
# table(BPdata$Open_science, useNA = "always")

#filter and make binomial
BPdata %>%
  filter(!is.na(Awardee_list_earliest_year)) %>%
  filter(Open_science == "yes" | Open_science == "no") %>%
  mutate(Open_science_num = recode(Open_science,
                                   yes = 1,
                                   no = 0)) -> BPdata4
```

Logistic plot of valueing Open Science practices relatively to award age:

```
BPdata4 %>%
  ggplot(aes(x = Awardee_list_earliest_year, y = Open_science_num)) +
  geom_point(alpha = 0.2) +
  stat_smooth(method = "glm",
              method.args = list(family = binomial),
              se = TRUE) +
  xlab("Year of first listed awardee") +
  ylab("Probability of Open_science") +
  theme_minimal()
```

```
## `geom_smooth()` using formula = 'y ~ x'
```

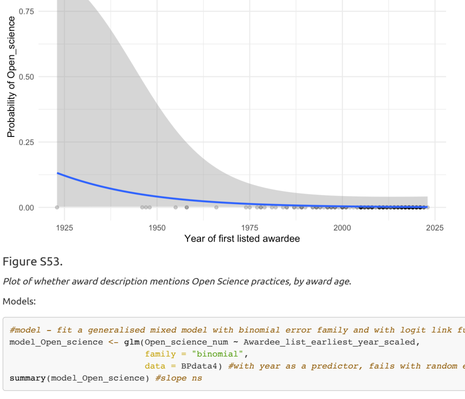


Figure S53.

*Plot of whether award description mentions Open Science practices, by award age.*

Models:

```
#model - fit a generalised mixed model with binomial error family and with logit link function, award as a random e
model_Open_science <- glm(Open_science_num ~ Awardee_list_earliest_year_scaled,
                         family = "binomial",
                         data = BPdata4) #with year as a predictor, fails with random effects
summary(model_Open_science) #slope ns
```

```
##
## Call:
## glm(formula = Open_science_num ~ Awardee_list_earliest_year_scaled,
##     family = "binomial", data = BPdata4)
##
## Coefficients:
##                                   Estimate Std. Error z value Pr(>|z|)
## (Intercept)                        -5.7776     1.3438  -4.299 1.71e-05 ***
## Awardee_list_earliest_year_scaled  -0.7611     0.5281  -1.441     0.15
## ---
## Signif. codes:  0 '***' 0.001 '**' 0.01 '*' 0.05 '.' 0.1 ' ' 1
##
## (Dispersion parameter for binomial family taken to be 1)
##
##     Null deviance: 12.520  on 192  degrees of freedom
## Residual deviance: 10.936  on 191  degrees of freedom
## AIC: 14.936
##
## Number of Fisher Scoring iterations: 8
```

```
# report::report(model_Open_science) #use the reports package (Makowski et al. 2021) to summarize the analysis
```

### Author contributions

```
#save vector for  reordering in the plot
position_order <- AUcontr$author_order

#reorder in reverse author position
AUcontr <- AUcontr %>% arrange(desc(position_order))

#make names a factor
AUcontr$name <- factor(AUcontr$name, levels = AUcontr$name)

#change to long format
AUcontr_long <- gather(AUcontr, contribution, counts, piloting_documentation:finalising_manuscript_checks_feedback,

#remove rows with 0 counts
AUcontr_long <- filter_if(AUcontr_long, is.numeric, all_vars((.) != 0))

# #reinforce factor orders
AUcontr_long$name <- factor(AUcontr_long$name, levels = AUcontr$name)

#make bubble plot
ggplot(AUcontr_long,
       aes(x = fct_inorder(contribution),
           y = name,
           colour = counts,
           size = counts)) +
  geom_point() +
  geom_text(data = subset(AUcontr_long, grepl("(SA)", AUcontr_long$contribution)),
            aes(label = counts),
            colour = "white",
            size = 3) +
  scale_x_discrete(position = "top", expand = expansion(mult = 0.1)) +
  scale_y_discrete(expand = expansion(mult = 0.1)) +
  theme_minimal() +
  theme(axis.text.x = element_text(angle = 90, hjust = 0, vjust = 0)) +
  scale_size_continuous(range = c(2, 6)) +
  labs(x = NULL, y = NULL) +
  theme(legend.position = "none")
```

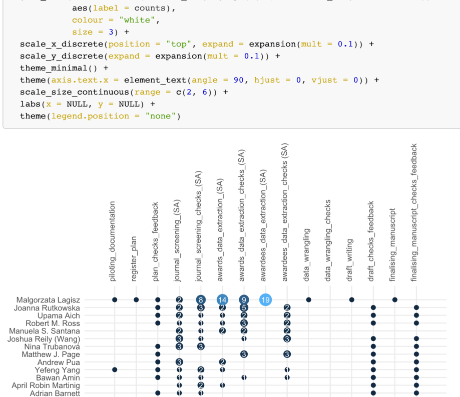


Figure S54.

*Plot of individual author contributions. Authors are listed in the manuscript authorship order. Contributions are listed in chronological order. Dots without numbers indicate contributions. Numbers within dots show numbers of completed Subject Areas, as relevant.*

### Session Info

```
sessionInfo()
```

```
## R version 4.3.1 (2023-06-16)
## Platform: x86_64-apple-darwin20 (64-bit)
## Running under: macOS Ventura 13.5.2
##
## Matrix products: default
## BLAS:   /Library/Frameworks/R.framework/Versions/4.3-x86_64/Resources/lib/libRblas.0.dylib
## LAPACK: /Library/Frameworks/R.framework/Versions/4.3-x86_64/Resources/lib/libRlapack.dylib;  LAPACK version 3.
11.0
##
## locale:
## [1] en_US.UTF-8/en_US.UTF-8/en_US.UTF-8/C/en_US.UTF-8/en_US.UTF-8
##
## time zone: Australia/Sydney
## tzcode source: internal
##
## attached base packages:
## [1] stats     graphics  grDevices utils     datasets  methods   base
##
## other attached packages:
##  [1] ggpattern_1.1.0-0 sjPlot_2.8.15     lme4_1.1-34       Matrix_1.6-1.1
##  [5] wacolors_0.3.1    tokenizers_0.3.0  stopwords_2.3     tidytext_0.4.1
##  [9] ggbeeswarm_0.7.2  patchwork_1.1.3   ggupset_0.3.0     readxl_1.4.3
## [13] scales_1.2.1      ggcharts_0.2.1    ggimage_0.3.3     knitr_1.44
## [17] DT_0.29           here_1.0.1        lubridate_1.9.2   forcats_1.0.0
## [21] stringr_1.5.0     dplyr_1.1.3       purrr_1.0.2       readr_2.1.4
## [25] tidyr_1.3.0       tibble_3.2.1      ggplot2_3.4.3     tidyverse_2.0.0
##
## loaded via a namespace (and not attached):
##  [1] rlang_1.1.1        magrittr_2.0.3     compiler_4.3.1     mgcv_1.8-42
##  [5] vctrs_0.6.3        pkgconfig_2.0.3    crayon_1.5.2       fastmap_1.1.1
##  [9] backports_1.4.1    magick_2.7.5       labeling_0.4.3     utf8_1.2.3
## [13] rmarkdown_2.25     tzdb_0.4.0         nloptr_2.0.3       bit_4.0.5
## [17] xfun_0.40          cachem_1.0.8       jsonlite_1.8.7     highr_0.10
## [21] SnowballC_0.7.1    sjmisc_2.8.9       ggeffects_1.3.1    parallel_4.3.1
## [25] broom_1.0.5        R6_2.5.1           bslib_0.5.1        stringi_1.7.12
## [29] boot_1.3-28.1      jquerylib_0.1.4    cellranger_1.1.0   estimability_1.4.1
## [33] Rcpp_1.0.11        modelr_0.1.11      pacman_0.5.1       splines_4.3.1
## [37] timechange_0.2.0   tidyselect_1.2.0   rstudioapi_0.15.0  yaml_2.3.7
## [41] sjlabelled_1.2.0   lattice_0.21-8     withr_2.5.1        bayestestR_0.13.1
## [45] evaluate_0.22      gridGraphics_0.5-1 pillar_1.9.0       janeaustenr_1.0.0
## [49] insight_0.19.5     ggfun_0.1.3        generics_0.1.3     vroom_1.6.3
## [53] rprojroot_2.0.3    hms_1.1.3          munsell_0.5.0      minqa_1.2.6
## [57] xtable_1.8-4       glue_1.6.2         emmeans_1.8.8      tools_4.3.1
## [61] fs_1.6.3           mvtnorm_1.2-3      grid_4.3.1         colorspace_2.1-0
## [65] nlme_3.1-163       performance_0.10.5 beeswarm_0.4.0     vipor_0.4.5
## [69] cli_3.6.1          fansi_1.0.4        sjstats_0.18.2     gtable_0.3.4
## [73] yulab.utils_0.1.0  sass_0.4.7         digest_0.6.33      ggplotify_0.1.2
## [77] farver_2.1.1       htmlwidgets_1.6.2  memoise_2.0.1      htmltools_0.5.6
## [81] lifecycle_1.0.3    bit64_4.0.5        MASS_7.3-60
```
